# Diversity and distribution of landscape types in Norway

**DOI:** 10.1101/2020.09.28.316331

**Authors:** Trond Simensen, Lars Erikstad, Rune Halvorsen

## Abstract

Norwegian landscapes are changing at an increasingly rapid rate, and systematically structured information about observable landscape variation is required for knowledge-based management of landscape diversity. Here we present the first version of a complete, area-covering, evidence-based landscape-type map of Norway, simultaneously addressing geo-ecological, bio-ecological and land-use related variation at the landscape level. We do so by applying map algebra operations on publicly available geographical data sets with full areal coverage for Norway. The type system used in the mapping is supported by systematically structured empirical evidence. We present the results of the mapping procedure, including the geographical distribution and descriptive statistics (abundance and areal coverage) for each of the identified landscape types. We identify nine major landscape types based on coarse-scale landform variation and, within the six inland and coastal major types, 284 minor landscape types are defined based on the composition of geo-ecological, bio-ecological, and land use-related landscape properties. The results provide new insights into the geography of Norwegian marine, coastal and inland landscapes. We discuss potential errors, uncertainties and limitations of the landscape-type maps, and address the potential value of this new tool for research, management and planning purposes.

## Introduction

### Landscape characterisation and mapping in Norway

Norway is widely recognised as a ‘hot spot’ of European landscape diversity (Mücher et al. 2008; Ciglič & Perko 2013), comprising an exceptional range of variation in climatic conditions, bedrock, landforms, vegetation and land-use within a relatively small area (Moen 1999; Bakkestuen et al. 2008). Like in most other European countries (Plieninger et al. 2018), landscapes in Norway are changing at an increasingly rapid rate (Eiter & Pothoff 2007). Since the development of land-use policies often implies choices between irreconcilable views on the desired utilisation of a landscape, there is a growing demand for planning and management strategies that combine the protection of landscape diversity with sustainable use of landscape resources. Systematically structured information about observable landscape variation is a prerequisite for knowledge-based spatial planning and landscape management (Marsh 2005) and essential for fulfilment of obligations set by international conventions such as the European Landscape Convention (Council of Europe 2000; ratified by Norway 2004). Moreover, the aim of the Norwegian Nature Diversity Act (2009) to protect ‘biological, geological and landscape diversity’ and promote conservation and sustainable use of the ‘full range of variation of habitats and landscape types’ throughout the nation presupposes knowledge about the abundance and spatial distribution of ‘landscape types’.

The conceptual idea of assigning landscapes to types is rooted in the tradition of systematic physical geography or ‘landscape geography’, the aim of which is to present and explain typologies of similar landscapes based on their material content (Holt-Jensen 2018). Because geomorphological, ecological and human land-use related processes are tightly intertwined, many landscape elements tend co-vary in predictable and recurrent patterns throughout larger regions (Bailey 2009). This is exemplified by a mountain plain in Hardangervidda (Buskerud) which is more similar to a distant mountain plain in Valdresflye (Oppland; > 100 km away) or even in Finnmarksvidda (Finnmark; > 1000 km away) than to any of the valleys surrounding the plain, with respect to its content of landforms, ecosystems and other landscape elements. Accordingly, grouping of similar landscapes into types is a powerful way to communicate landscape information effectively, because affiliation to type alone will provide an extensive amount of information about any singular individual of that particular type. Since a type in a type system comprises a predictable, ‘normal’ amount of landscape variation, affiliation to type is also a useful reference and a good starting point for assessment of the unique character and properties of individual landscapes (see e.g. Phillips 2007).

The concept of landscape types has been used informally in geographical and ecological literature in Norway since the early 20^th^ century (e.g. Sømme 1938, Nordhagen 1943), but comprehensive and evidence-based landscape type maps have until now not been established for Norway. National landscape characterisation efforts have followed a regional geographic tradition by identifying and describing the individual character of particular landscape areas, i.e. by identifying landscapes or regions with a high degree of internal similarity. Notable earlier efforts within the regional geographic tradition include interpretative and holistic approaches (e.g. Nordisk Ministerråd 1987; Puschmann 2005) as well as more observer-independent, data-driven methods (e.g. Strand 2011; Krøgli et al. 2015). Regional geographic landscape analyses have served as a useful framework for several applied purposes (Strand 2011). Nevertheless, regional maps lack the thematic and spatial resolution necessary to serve as a relevant knowledge base for land-use policies and environmental impact assessments (EIAs) at the local regional scale (i.e. ~1:50 000, see, e.g. Helland et al. 2015). Most governmental guidelines for landscape analysis recommend a largely value-neutral description of the observable properties of a landscape as a starting point for character-, value- and suitability assessments based on context-specific criteria (see e.g. Helland et al. 2015, Norwegian Public Roads Administration 2018). To ensure better quality and consistency of general landscape descriptions, several Norwegian scientists have called for a more systematic, observer-independent and repeatable framework as a reference and a knowledge base for a multitude of applied purposes (see, e.g. Moen 1999, Strand 2011, Krøgli et al. 2015, Erikstad et al. 2015).

### EcoSyst – a new framework for systematisation of nature diversity

To meet the need for more detailed, systematic, observer-independent and area-covering information about nature’s diversity required by the Nature Diversity Act, the Norwegian Biodiversity Information Centre (NBIC) started development of the project ‘Nature in Norway’ in 2005 (NiN; Halvorsen et al. 2016). NiN has later been developed into a universal theoretical framework for systematisation Nature’s diversity, EcoSyst, which simultaneously addresses biotic and abiotic variation across different levels of organisation from microhabitats to landscapes (Halvorsen et al. 2020).

Since the term ‘landscape’ is understood and applied differently within the disciplines that have landscape as a subject of interest (such as geography, geology, geomorphology, ecology, history, archaeology and landscape architecture; Jones and Stenseke 2011), any attempt to systemtatise variation at the landscape level requires a clear definition of the term ‘landscape’. In the EcoSyst framework, landscape is recognised as a separate level of ecological diversity, simultaneously addressing biotic and abiotic variation in heterogeneous areas of kilometres-wide extent (see Noss 1990). The EcoSyst framework addresses the material, physical properties of the landscape, defined as ‘as a more or less uniform area characterised by its content of observable, natural and human-induced landscape elements, i.e. natural or human-induced objects or characteristics, including spatial units assigned to types at an ecodiversity level lower than the landscape level, which can be identified and observed on a spatial scale relevant for the landscape level of ecodiversity’ (Halvorsen et al. 2020). Furthermore, ‘landscape element’ is defined as ‘a natural or human-induced object or characteristic, including spatial units assigned to types at an ecodiversity level lower than the landscape level, which can be identified and observed on a spatial scale relevant for the landscape level of ecodiversity’. ‘Landscape types’ are defined as reoccurring units ‘more or less uniform areas characterised by their content of observable, natural and human-induced landscape elements’ (Halvorsen et al. 2020).

As a first step towards a new Norwegian landscape type map based upon EcoSyst principles (see Appendix S1), a pilot NiN landscape typology was developed for Nordland county (Erikstad et al. 2015). Based on the experience gained from the Nordland pilot, the study area was expanded to encompass the entire country (Simensen, Halvorsen & Erikstad 2020). The overall methodological approach was similar in both projects (see Fig. 1). First, detailed analyses of landscape variation in a *sample* of observation units (landscapes) were conducted, and the results used to derive a type system (Fig. 1, steps 1–2). With some adaptions, this type system was used as the platform on which a rule-based, semi-automated procedure for mapping of the entire study area was built (steps 3–6). The empirical basis for the landscape type system applied here is multivariate analyses of 85 landscape variables collected in a stratified sample of 100 test areas (25×25 km) that cover 56 400 km^2^ (about one-sixth of mainland Norway, see Simensen, Halvorsen & Erikstad 2020). In addition to serving as the empirical basis for a tentative type system, the analyses provide general knowledge about the distribution of landscape elements throughout the Norway (Simensen, Halvorsen & Erikstad 2020). Within ‘major landscape types’, primarily defined by coarse-scale landform variation, a unique set of ‘complex landscape gradients’ (CLGs) were identified. CLGs are defined as ‘abstract, continuous variable that expresses more or less gradual, co-ordinated change in a set of more or less strongly correlated landscape variables’ (Halvorsen et al. 2020). Examples of CLGs are ‘inner-outer coast’, ‘vegetation cover’ and ‘land-use intensity’. For the first time, a novel procedure to quantify the similarity between different landscapes (‘ecodiversity distance’) was applied (see Halvorsen et al. 2020). The tentative, abstract (i.e. non-spatial) landscape typology was obtained by combining segments along sets of CLGs specific to each major type.

**Fig. 1.**
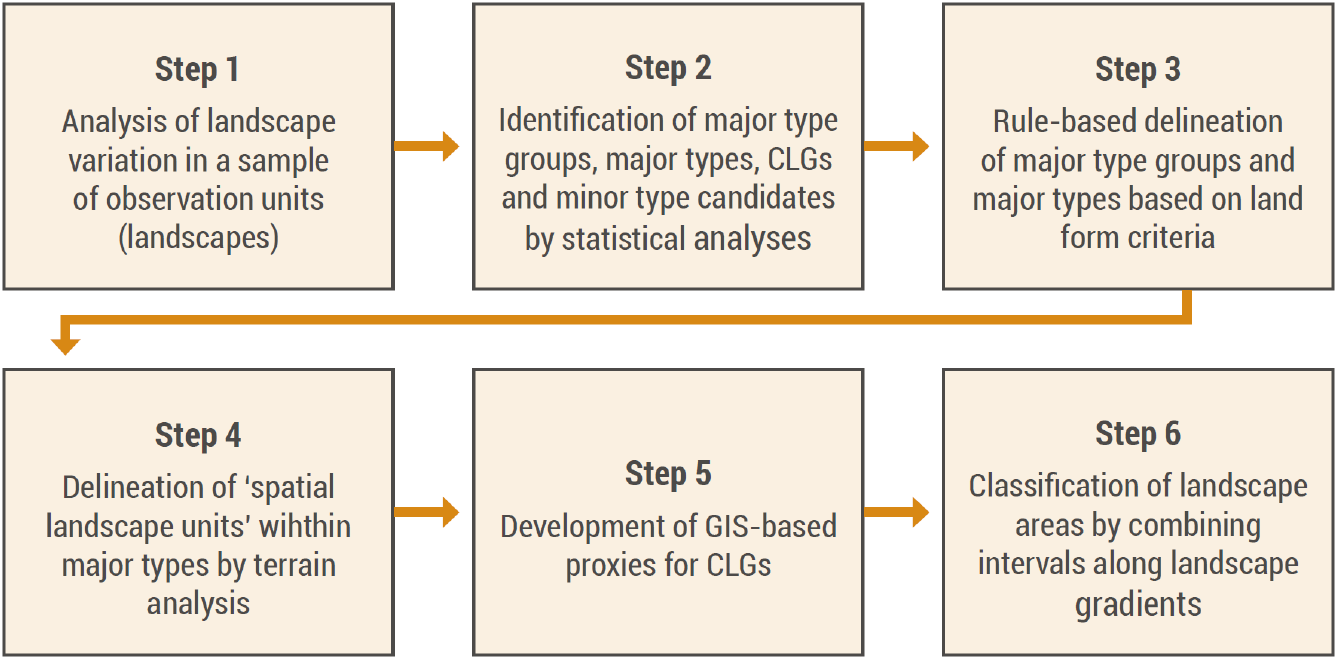
The six-step methodological approach used in the development of a landscape type system for Norway. After an initial phase comprising analysis of landscape variation in a sample of observation units (steps 1–2), the analytic results were translated to a type system that was adapted to applied landscape type mapping by rules for geographical delineation of spatial landscape units and assignment to landscape types (steps 3–6). Whereas steps 1–2 are described in detail in Simensen et al. (2020), this article documents the method for applied landscape type mapping (steps 3–5). CLG = complex landscape gradient.

Few patterns in ecology make sense unless viewed in an explicit geographic context (Lomolino et al. 2017). This papers answers to the challenge of translating the abstract EcoSyst (NiN) landscape typology of Simensen et al. (2020) into the first version of a complete, area-covering, evidence-based landscape-type map for Norway. We accomplish this aim by applying simple map algebra operations on publicly available geographical data sets with full areal coverage for Norway. Furthermore, we present the results of the mapping, including the geographical distribution and descriptive statistics (abundance and areal coverage) for each of the identified landscape types. Finally, we discuss potential errors, uncertainties and limitations of the landscape-type maps, and address the potential value of this new tool for research, management and planning purposes.

## Materials and methods

### Study area

The study area spanned latitudes from 57°57’N to 71°11’N and longitudes from 4°29’E to 31°10’E and comprised the entire mainland of Norway, including the coastal zone and marine areas. The range of variation in natural conditions found in Norway includes most of the variation found in the circumboreal zone (Bryn et al. 2018), including both terrestrial, marine, limnic and snow and ice ecosystems (Halvorsen et al. 2016). All seven bioclimatic temperature-related vegetation zones commonly recognised in northern Europe, from boreo-nemoral to high alpine, occur in Norway (Bakkestuen et al. 2008). Norway has a high mineral and bedrock diversity, and a high diversity of landforms (Ramberg et al. 2008). The diversity of Norwegian landscapes is enhanced by historical land-use such as domestic grazing, outfield fodder collection, heath burning, reindeer husbandry, forestry, and industrial, urban and recreational development (Almås et al. 2004; Hansen & Olsen 2004; Jacobsen & Follum 2008). Note that the 2019 version of the administrative division of Norway into counties and municipalities is used for geographical terms in this study.

### Source data

All the source data used to derive the landscape-type maps were obtained from publicly available geographical data sets with full areal coverage throughout Norway (Table 1). The basis data consisted of: 1) continuous variables (e.g. digital elevation model; DEM); 2) categorical landscape and land-cover data (e.g. AR 50 land-cover types); and 3) point and line data (e.g. buildings and infrastructure). All spatial data were converted to raster format with resolution 100×100 m, or adapted to this grain size by resampling or rasterisation from vector formats. We obtained the DEM by combining a terrestrial DEM interpolated from 20 m height contour lines (N50 topographic maps) with a marine DEM (bathymetric data, 50 m resolution). For 275 freshwater lakes, bathymetry data were avialable from the Norwegian Water Resources and Energy Directorate while for the remaining freshwater lakes >2 km^2^, we interpolated the bathymetry by inverse distance weighting based on DEM values for the terrestrial surroundings (see appendix S6). Finally, terrestrial, marine and freshwater DEMs were combined to a seamless DEM and adapted to 100 m resolution by spatial aggregation (Rød 2015).

**Table 1.**
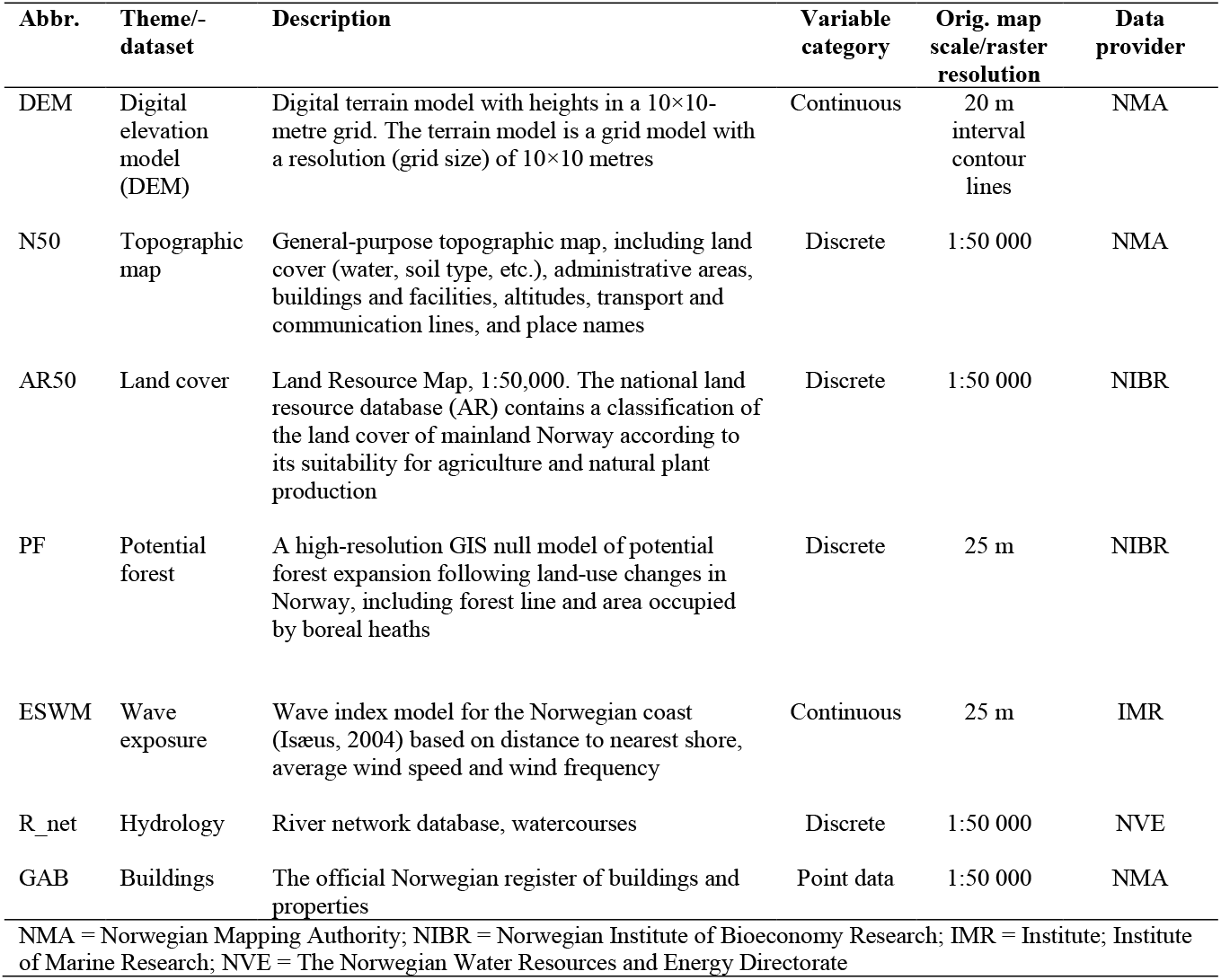
Baseline input map layer information. All layers were converted to raster format with resolution 100×100m.

| Abbr. | Theme/-dataset | Description | Variable category | Orig. map scale/raster resolution | Data provider |
| --- | --- | --- | --- | --- | --- |
| DEM | Digital elevation model (DEM) | Digital terrain model with heights in a 10×10-metre grid. The terrain model is a grid model with a resolution (grid size) of 10×10 metres | Continuous | 20 m interval contour lines | NMA |
| N50 | Topographic map | General-purpose topographic map, including land cover (water, soil type, etc.), administrative areas, buildings and facilities, altitudes, transport and communication lines, and place names | Discrete | 1:50 000 | NMA |
| AR50 | Land cover | Land Resource Map, 1:50,000. The national land resource database (AR) contains a classification of the land cover of mainland Norway according to its suitability for agriculture and natural plant production | Discrete | 1:50 000 | NIBR |
| PF | Potential forest | A high-resolution GIS null model of potential forest expansion following land-use changes in Norway, including forest line and area occupied by boreal heaths | Discrete | 25 m | NIBR |
| ESWM | Wave exposure | Wave index model for the Norwegian coast (Isæus, 2004) based on distance to nearest shore, average wind speed and wind frequency | Continuous | 25 m | IMR |
| R_net | Hydrology | River network database, watercourses | Discrete | 1:50 000 | NVE |
| GAB | Buildings | The official Norwegian register of buildings and properties | Point data | 1:50 000 | NMA |
| NMA = Norwegian Mapping Authority; NIBR = Norwegian Institute of Bioeconomy Research; IMR = Institute; Institute of Marine Research; NVE = The Norwegian Water Resources and Energy Directorate |  |  |  |  |  |

### Methods

Based on the source data, we derived the variables that were used to identify complex landscape gradients by multivariate statistical analyses and, subsequently, to develop the tentative landscape type hierarchy described in Simensen, Halvorsen & Erikstad (2020). The process by which the complex landscape gradients (CLGs) and landscape types were ‘translated’ into spatially explicit variables (i.e. maps) made use of three different approaches:

1. direct spatial projection of variables used in the statistical analyses of landscape variation;
2. application of well-documented GIS-algorithms that replicated (spatially) the results from the statistical analyses (e.g. geomorphometric analyses; see Hengl & MacMillan 2009); and 3) development of new geocomputation methods to derive spatial proxies for the identified complex landscape gradients. Back-transformation of values from the statistical analyses allowed for fine-tuning of criteria for separation between major types based on identified important terrain-characterising variables such as relative relief and the proportion of flat terrain within a larger area.

Derived variables used for modelling were either obtained by reclassification and filtering of categorical land-cover data or by continuous neighbourhood calculations, also referred to as ‘focal statistics’ (Lovelace et al. 2019), or as a ‘moving window’ (Cushman et al. 2010). Focal statistics considers a central cell and its neighbours within a specified distance (window). The window moves over the landscape one cell at a time, calculating the selected metric (e.g. sum) within the window (neighbourhood) and returning that value to the focal cell (see Appendix S2). Our goal was to address landscape variation corresponding to the spatiotemporal domain defined by Dikau (1989) as ‘meso-scale’ (i.e., abiotic and biotic patterns occurring at spatial scales of approximately 10^6^–10^10^ m^2^ in response to processes operating at temporal scales of 10^1^–10^4^ years). We therefore used a neighbourhood circle with a radius of 3000 m around the processing cell to derive coarse-scale geomorphometric variables, as recommended by Pike et al (2009). We derived fine-scale geomorphometric variables and continuous land-cover variables by focal statistics using a radius of 500 m (see Pike et al. 2009; see also Table 2 and Appendix S3–S5).

**Table 2.** Complex landscape gradients (CLGs) with short descriptions. Abbreviations and explanations: Code = CLG term in Norwegian (see Erikstad et al. 2019); KA = coastal hills and mountains; KF = coastal fjords; KS = coastal plains; IA = inland hills and mountains; ID = inland valleys; IS = inland plains; Data = original source of data used for geocomputation (see Table 1); Geocomp. = geocomputation methods; MA = map algebra; focal = focal raster calculations; Vrm = vector ruggedness measure; Tpi = topographic position index; RR = relative relief; Log-tr. = log-transformation; reclass = reclassification; Appendix = reference to detailed description of geocomputation method in Appendix (Supplementary material).

| Code | Complex landscape gradient | Description | Functional variable category | Number of segments | Major-type group | Major type | Data source | Geo-comp. | Detailed descriptions |
| --- | --- | --- | --- | --- | --- | --- | --- | --- | --- |
| KLK-RE-IA | Relief in hills and mountains | Terrain-form variation within (inland) hill and mountain landscapes; from depressions in hills and mountains (RE-IA·1), via undulating inland hills and mountains (RE-IA·2), moderately rugged hills and mountains (RE-IA·3) and rugged hills and mountains (RE-IA·4) to steep and rugged hills and mountains (RE-IA·5). | Geo-ecological | 5 | Inland | IA | DEM | MA; focal; Vrm; Tpi; RR; reclass | Appendix S5.2 |
| KLK-RE-ID-KF | Relief in valleys and fjords | Terrain-form variation within (inland) valley and (coastal) fjord landscapes; as expressed by the depth/width ratio of the valley/fjord relative to its surroundings; from wide fjord/valley (RE-KF·1) via open fjord/valley (RE-KF·2) and narrow fjord/valley (RE-KF·3) to deeply cut fjord/valley (RE-KF·4). | Geo-ecological | 4 | Inland, coast | ID<br>KF | DEM | MA; focal; reclass | Appendix S5.2 |
| KLK-RE-KS | Relief in coastal plains | Terrain-form variation within coastal plains from flat (RE-KS·1) via relatively flat (RE-KS·2) coastal plains to rugged coastal plains, often with remnant peaks (RE-KS·3). | Geo-ecological | 3 | Coast | KS | DEM | MA; focal; RR; reclass | Appendix S5.3 |
| KLK-IYK | Inner-outer coast | Variation in coastal landscapes from areas with ‘inland properties’ on the inner side of larger islands, hardly exposed to the harsh conditions of the open sea (protected inner coast; IYK·1), to the outer coast, directly exposed to the actions of wind, waves and ocean currents (strongly wave-exposed outer coast; IYK·5). The intermediate segments are: moderately protected coast (IYK·2); moderately wave-exposed coast (IYK·3); and wave-exposed outer coast (IYK·4). | Geo-ecological | 4 | Coast | KS | ESWM, N50<br>R_net | MA; focal; Log-tr.; reclass | Appendix S5.4 |
| KLK-KA | Distance to coast | Variation in inland landscape properties from near-coastal inland plains (situated < 5 km from the coastline) to inland plains in the interior of the land mass. | Geo-ecological | 2 | Inland | IS | AR50 | MA; reclass | Appendix S5.5 |

| **Code** | **Complex landscape gradient** | **Description** | **Functional variable category** | **Number of segments** | **Major-type group** | **Major type** | **Data source** | **Geo-comp.** | **Detailed descriptions** |
| --- | --- | --- | --- | --- | --- | --- | --- | --- | --- |
| KLG–IP | Abundance of lakes | Variation within valleys from landscapes without lakes or with small lakes only (IP·1; all lakes < 2 km<sup>2</sup>), via valleys with medium-sized lakes (IP2·1; lakes 2–8 km<sup>2</sup>), to valleys with large lakes, typically ‘inland fjords’ (IP2·1; > 8 km<sup>2</sup>). | Geo-ecological | 3 | Inland | ID | N50 | MA; reclass | Appendix S5.6 |
| KLG–VP | Abundance of wetlands | Variation within inland landscapes in the areal cover of wetlands (mires and springs) and the abundance of small lakes and tarns (which are often associated with wetlands): from low to medium abundance (VP·1) to high abundance (VP·2). | Geo-ecological | 2 | Inland, coast | IS, KF, KS | N50 | MA; focal; reclass | Appendix S5.7 |
| KLG–BP | Glacier presence | Variation in the presence of glacier(s); BP·1: glacier absent; BP·2: glacier present. | Geo-ecological | 2 | Inland | IA, IS, ID | AR50 | MA; reclass | Appendix S5.8 |
| KLG–VE | Vegetation cover | Variation in vegetation cover from forested or potentially forested areas below the climatic forest line (VE·1) to barren mountains, without or with sparse vegetation cover (VE·4). The intermediate segments are: boreal heath-dominated areas below the climatic forest line, kept open due to historical land-use by logging and grazing (VE·2) and open mountain heaths (VE·3). | Bio- ecological | 4 | Inland, coast | IA, IS, ID, KF | AR50, PF | MA; reclass | Appendix S5.9 |
| KLG–AI | Land-use intensity | Variation in impact by human infrastructure (agricultural land-use excepted) from low (AI·1) and intermediate (AI·2) to settlement (village or small town; AI·3) and city (AI·4), expressed by an index (range: 0–13.2) that integrates the abundances of buildings, roads and other visible signs of human infrastructure. | Land-use-related | 4 | Inland, coast | IA, IS, ID, KF, KS | GAB N50 | Focal; reclass | Appendix S5.10 |
| KLG–JP | Agricultural land-use intensity | Variation in expressed agricultural land-use intensity, from low (JP·1) to high (JP·2). | Land-use-related | 2 | Inland, coast | IA, IS, ID, KF, KS | N50 | Focal; reclass | Appendix S5.11 |

### Delineation of major landscape types

Supported by the statistical analyses, we assigned spatial landscape units to one of the three major-type groups by their relation to the coastline. ‘Coastal landscapes’ were identified as landscapes in the interface between the land and marine environments; all landscape units that contained a segment of the continuous coastline of a land area were assigned to the coastal landscapes major-type group. We assigned the spatial landscape units to major type group by zonal operations identifying presence of marine, coastal and terrestrial pixels (Rød 2015).

The statistical analyses (Simensen, Halvorsen & Erikstad 2020) gave support for recognition of three meso-scale landforms in coastal and inland Norway: ‘plains’, ‘hills and mountains’ and ‘fjord and valleys’. We derived these meso-scale landform types by morphometric calculations based upon the DEM. We first delineated valleys and fjords by focal DEM calculations, using 3 km neighbourhoods. Our fjord/valley-model (described in detail in Appendix S4) identified elongated depressions in the terrain by application of the algorithms ‘terrain position index’ (TPI; Gallant & Wilson 2000); the ‘Top Hat approach’ including ‘valley index’ and ‘valley depth‘ (Rodriguez et al. 2002), and ‘peak- and ridge detection’ (Jasiewicz & Stepinski 2013; see appendix S3 and S4 for further details). Valleys and fjords were then split into two major types by their location relative to the coastline by converting identified valleys to vector data and selecting features by relationships to the coastline and subsequently applying a cost-distance calculation (see Appendix S4). Valleys completely submerged under marine water were then assigned to the major type ‘marine valleys’.

‘Coastal plains’ were defined as landscapes with coastline (not already defined as valleys and fjords), situated within ±50 m above/below sea level. The latter condition was operationalised by demanding that >0.5 of all cells within 1 km neighbourhoods were situated in this interval. Delimitation of ‘coastal plains’ on the inland side was made by applying a cost-distance grid. We applied the same procedure to identify ‘inland plains’, defined as terrestrial landscapes (without coastline) and marine plains (entirely submerged by sea water), by demanding height differences (relative relief) < 50 m within a 1 km neighbourhood. Due to highly variable quality of available soil type maps, we decided not to delineate ‘fine-sediment plains’ as a separate major type in this first version of the landscape type map, although this was tentatively identified as a major type on its own by the statistical analyses (see discussion). After delineating fjords, valleys and inland and coastal plains, all remaining areas were defined as ‘hills and mountains’. We then merged the major types to one map and joined areas below the minimum polygon size (4 km^2^) with adjacent polygons according to the procedures outlined in Appendix S4. Finally, polygons of hills and mountain were assigned to major-type group and major type based upon their location relative to the coastline (see above).

### Calculation of positions along complex landscape gradients

We developed spatially explicit proxies (i.e. maps) for the 11 CLGs identified in the statistical analyses (Table 2). Three types of proxies were developed: a) indices based on focal statistics; b) indices based on the presence or absence of landscape elements; and c) composite indices (e.g. obtained as the sum of two or more indices, quantifying and weighing the frequencies of landscape elements and properties). We derived proxies for the three geo-ecological CLGs related to ‘relief’ within coastal plains, coastal fjords, inland valleys and inland hills and mountains, respectively, by morphometric terrain surface calculations based on the DEM (see Appendices S5.1–3). We calculated positions along the CLG ‘inner-outer coast’ as a weighted sum of three elements: 1) amount/abundance of rivers 2) island size; and 3) wave exposure (see Appendix S5.4). The CLG ‘distance to coast’ separated ‘near-coastal’ inland plains from other inland plains and positions were obtained as the Euclidean distance from the nearest coastline-pixel. We used 5 km to the coastline as the threshold for separating ‘near-coastal’ from other inland plains (see Appendix S5.5). ‘Abundance of lakes’ was calculated by classification of lake size into three classes: small lakes <2 km^2^; medium-sized lakes 2–8 km^2^ and large lakes > 8 km^2^ (see Appendix S5.6). Calculations of positions along the CLG ‘abundance of wetlands’ was derived from combining focal calculations of wetlands and point data for small lakes (see Appendix S5.7). The CLG ‘glacier presence’ was obtained directly from reclassification of land-cover data (AR50; Appendix S5.8). We derived the bio-ecological CLG ‘vegetation cover’ by reclassification of land-cover classes from AR50 (obtained from NIBIO). The reclassified model was then combined with the GIS-model for ‘potential forest’ (Bryn et al. 2013; se Appendix S5.9 for reclassification scheme). Positions along the CLG ‘Land-use intensity’ were calculated by a weighted summation of focal calculations of a) buildings; and b) technical infrastructure (see Appendix S5.10 and Erikstad et al. 2013). Positions along the CLG ‘agricultural land-use intensity’ was obtained by focal calculations of arable land derived from AR50 (see Appendix S5.11).

The spatial models for the complex landscape gradients followed the results of the statistical analyses closely. Still, a few deviations and adaptions to functional had to be introduced to enhance mapping functionality. In order to account for ‘extreme’ landscape variation along CLGs that was absent or represented with very few observation units in the sample of observation units used for the statistical analyses, we subjectively introduced four new segments along three CLGs to account for landscape elements that dominate the landscape where they are present: AI·3 (village or small town); city (AI·4); high agricultural land use intensity (JP·2); and glacier presence (BP·2).

### Delineation of spatial landscape units and assignment to minor landscape types

A method for segmentation of the target area into concrete spatial landscape units is integrated into the landscape-type mapping process as outlined in EcoSyst (see Halvorsen et al. 2020). Within each major landscape type, we delineated spatial landscape units by a rule-based division of the landscape into discrete spatial units for subsequent classification into landscape types. We used ridge lines, inflexion points and other breaking points in the terrain curvature to delineate spatial landscape units that are maximally homogeneous with respect to terrain properties and landform characteristics. For this purpose, we adapted and applied methods for delineation of drainage basins (Horton 1932; Gruber & Peckham 2009), adjusted to the scale of our analysis (spatial landscape units from 4–20 km^2^). Within areas with mainly convex landforms (e.g. hills and mountains), we applied the same procedure on an inverted DEM. This was motivated by our intention to identify areas that share terrain surface properties rather than delimit strictly hydrological entities (e.g. a mountain top; se details in Appendix S6).

We assigned segments along relevant CLGs to spatial landscape units by summarising values for each CLG within each spatial landscape unit. For this purpose, we used zonal statistics with majority-calculations, or by registering the abundance of specific landscape elements within each spatial landscape unit. When, e.g. the majority (i.e. > 50%) of the pixels in a polygon (landscape unit) belonged to segment RE·5 within the CLC ‘relief in hills and mountains’, (i.e. steep and rugged hills and mountains), the spatial landscape unit were coded accordingly. We finally obtained the minor landscape types by map algebra, by adding relevant segments along all CLGs for each major landscape type (see Appendix S7). Every unique combination of segments along CLGs identified as important for the major type in question was, by definition, considered as one landscape type (Fig. 2; see Appendix S7 and S8 for details).

**Fig. 2.**
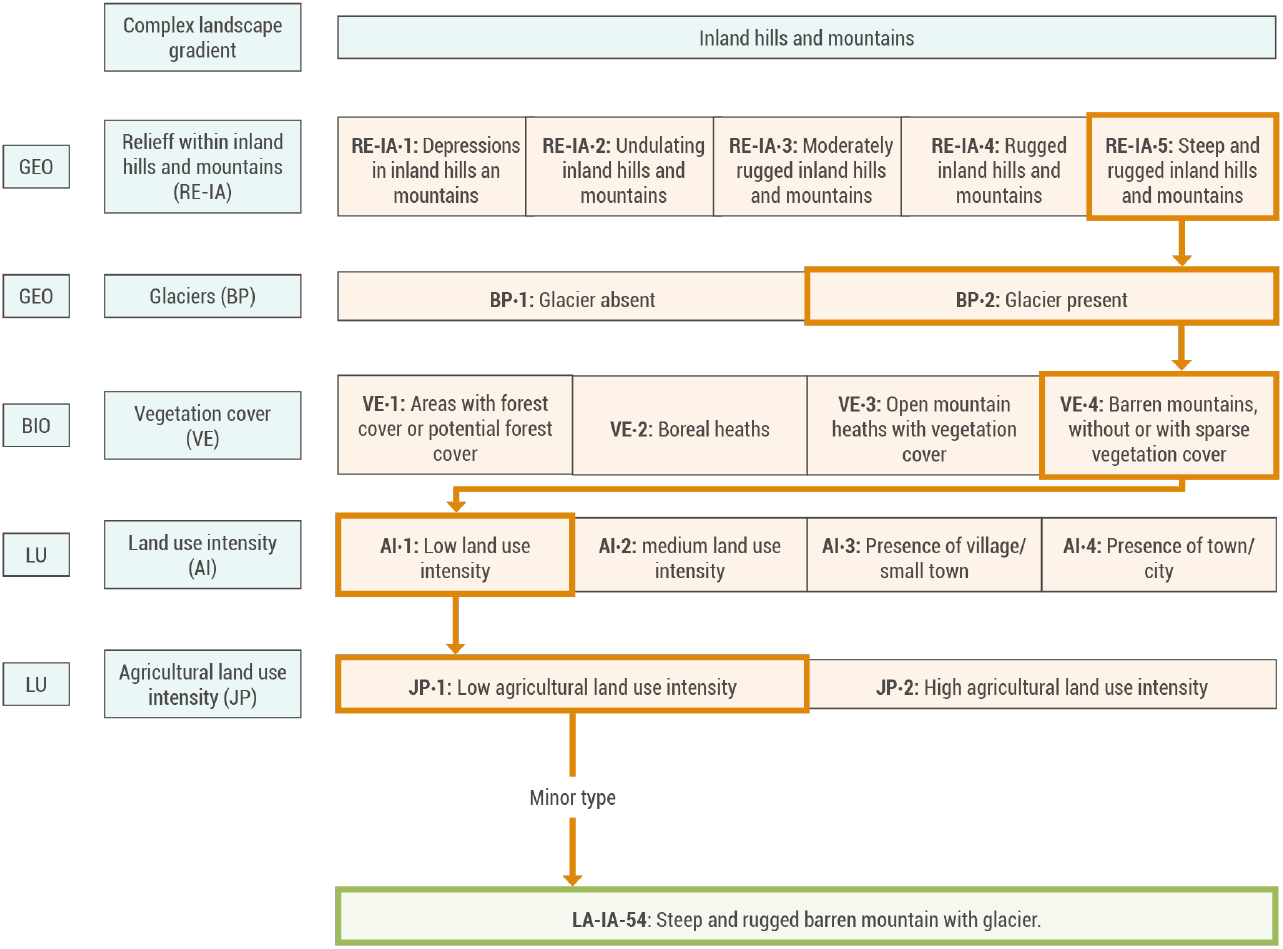
Assignment of spatial landscape units to minor landscape types within a major type. Each spatial landscape unit is assigned to minor type according to the combination of segments along each major-type specific important complex landscape gradient (iCLGs) that characterises the landscape unit. Type assignment is exemplified by the major type ‘inland hills and mountains’ and the minor landscape type ‘steep and rugged barren mountains with glacier’ (CLG-segment combination RE·5 – BP·2 – VE·4 – AI·1 – JP·l; type code IA524ll), shown by red boxes. Theoretically, major-type specific intervals along the five iCLGs for this major type can be combined in 5 · 2 · 4 · 4 · 2 = 320 ways. Most of these combinations are not realised, as exemplified by the unrealistic combination ‘steep mountains with glacier, forest cover, city and high agricultural land use’. In the NiN 2.2 landscape-type map for Norwegian coastal and inland landscapes, 54 of the potential 320 minor types within the major type ‘inland hills and mountains’ are realised. Abbreviations: GEO = geo-ecological CLG; BIO = bioecological CLG; LU = human land-use related CLG. Corresponding type-assignment schemes for all major types are provided in Appendix S7

The raster grids with type codes for all polygons for each major type were transformed from raster data to polygons, and merged to a complete landscape type map with complete areal coverage for the mapping area. The landscape types were given specific codes (e.g. IA-1, IA-2, etc.) and descriptive names based on the presence of expected content of landscape elements and CLG position (e.g. ‘steep and rugged mountains with glacier’; see Table 3 and Fig. S9). We generated textual descriptions of each minor landscape type by concatenating text strings with textual descriptions of each segment in CLGs included in each type. We described commonly occurring landscape elements (i.e. ecosystem types and landforms) for each minor types based on three sources: 1) the statistical analyses in Simensen, Halvorsen & Erikstad (2020); 2) data from distribution modelling of ecosystem types (Horvath et al. 2019; Simensen, Horvath et al. 2020); and 3) expert assessments by the authors.

**Table 3.** Total area of the ten most common minor landscape types within each major type, (marine landscapes not included). Code = major-type (bold text) and minor-type code; Major and minor landscape types = descriptive names for major (bold text) and minor landscape types; spatial landscape units (n) = number of spatial landscape units within each major and minor type. See also Appendix Table S9 for complete tables, including all minor types.

| Code | Major and minor landscape types | Spatial landscape unit (n) | Total area, km<sup>2</sup> | Area in % of major type | Area in % of total area |
| --- | --- | --- | --- | --- | --- |
| **KS** | **Coastal plains** | **4476** | **36,249** | **100.0** | **10.0** |
| KS-44 | Very wave-exposed outer undulating coastal plain | 844 | 6489 | 17.9 | 1.8 |
| KS-57 | Extremely wave-exposed outer undulating coastal plain | 720 | 5811 | 16.0 | 1.6 |
| KS-9 | Sheltered inner flat coastal plain | 674 | 5140 | 14.2 | 1.4 |
| KS-27 | Moderately wave-exposed undulating coastal plain | 634 | 4870 | 13.4 | 1.3 |
| KS-38 | Very wave-exposed outer flat coastal plain | 218 | 1962 | 5.4 | 0.5 |
| KS-52 | Extremely wave-exposed outer flat coastal plain | 198 | 1775 | 4.9 | 0.5 |
| KS-11 | Sheltered inner undulating coastal plain with village/small town | 176 | 1419 | 3.9 | 0.4 |
| KS-21 | Moderately wave-exposed flat coastal plain | 124 | 1064 | 2.9 | 0.3 |
| KS-1 | Sheltered inner coastal plain | 69 | 641 | 1.8 | 0.2 |
| KS-29 | Moderately wave-exposed undulating coastal plain with village/small town | 76 | 596 | 1.6 | 0.2 |
| KS | Other coastal plains (53 minor types)* | 743 | 6482 | 17.9 | 1.8 |
| **KF** | **Coastal fjords** | **3713** | **33,478** | **100.0** | **9.0** |
| KF-9 | Open fjord with settlements/infrastructure | 1060 | 9659 | 28.9 | 2.6 |
| KF-8 | Relatively open fjord | 1003 | 9134 | 27.3 | 2.5 |
| KF-17 | Narrow fjord | 587 | 4933 | 14.7 | 1.3 |
| KF-18 | Narrow fjord with settlements/infrastructure | 267 | 2354 | 7.0 | 0.6 |
| KF-2 | Open fjord with settlements/infrastructure | 219 | 2066 | 6.2 | 0.6 |
| KF-1 | Open fjord | 175 | 1649 | 4.9 | 0.4 |
| KF-11 | Relatively open fjord with village/small town | 117 | 1183 | 3.5 | 0.3 |
| KF-24 | Deeply cut fjord | 127 | 988 | 2.9 | 0.3 |
| KF-4 | Open fjord with village/small town | 48 | 453 | 1.4 | 0.1 |
| KF-20 | Narrow fjord with village/small town | 27 | 301 | 0.9 | 0.1 |
| KF | Other coastal fjords (16 minor types)* | 83 | 758 | 2.3 | 0.2 |
| KA | **Coastal hills and mountains** | 74 | 284 | 100.0 | 0.1 |
| KA-1 | Coastal hills- and mountains* | 74 | 284 | 100.0 | 0.1 |
| **ID** | **Inland valleys** | **11,029** | **94,607** | **100** | **25.8** |
| ID-32 | Open valley below the forest line | 2277 | 18,202 | 19.2 | 5.0 |
| ID-34 | Open valley below the forest line with settlements/infrastructure | 804 | 7394 | 7.8 | 2.0 |
| ID-1 | Wide valley below the forest line | 913 | 7311 | 7.7 | 2.0 |
| ID-65 | Narrow valley below the forest line | 718 | 5801 | 6.1 | 1.6 |
| ID-43 | Open barren mountain valley | 639 | 5318 | 5.6 | 1.4 |
| ID-38 | Open valley with boreal heath below the forest line | 516 | 4213 | 4.5 | 1.1 |
| ID-3 | Wide valley below the forest line with settlements/infrastructure | 420 | 3843 | 4.1 | 1.0 |
| ID-41 | Open valley with heath above the forest line | 441 | 3511 | 3.7 | 1.0 |
| ID-73 | Narrow barren mountain valley | 420 | 3250 | 3.4 | 0.9 |
| ID-45 | Open valley below the forest line with medium sized lakes | 273 | 2491 | 2.6 | 0.7 |
| ID | Other inland valleys (94 minor types)* | 3608 | 33273 | 35.2 | 9.1 |
| **IA** | **Inland hills and mountains** | **21,058** | **170,523** | **100.0** | **46.5** |
| IA-27 | Moderately rugged hills below the forest line | 2907 | 22,494 | 13.2 | 6.1 |
| IA-14 | Undulating hills below the forest line | 2585 | 20,367 | 11.9 | 5.6 |
| IA-1 | Depressions in hilly landscapes below the forest line | 2352 | 17,388 | 10.2 | 4.7 |
| IA-38 | Moderately rugged barren mountains | 2121 | 17,384 | 10.2 | 4.7 |
| IA-36 | Moderately rugged open heath mountains | 1351 | 10,867 | 6.4 | 3.0 |
| IA-33 | Moderately rugged hills and mountains with boreal heath | 1302 | 10,151 | 6.0 | 2.8 |
| IA-53 | Steep and rugged barren mountains | 1038 | 8366 | 4.9 | 2.3 |
| IA-23 | Undulating open heath mountains | 926 | 7464 | 4.4 | 2.0 |
| IA-46 | Rugged barren mountains | 945 | 6938 | 4.1 | 1.9 |
| IA-25 | Undulating barren mountains | 696 | 5902 | 3.5 | 1.6 |
| IA | Other inland hills and mountains (44 minor types)* | 4835 | 43202 | 25.3 | 11.8 |
| **IS** | **Inland plains** | **3413** | **31,834** | **100.0** | **8.8** |
| IS-13 | Inland undulating plain below the forest line with wetlands | 1005 | 9763 | 30.7 | 2.7 |
| IS-1 | Inland undulating plain below the forest line | 673 | 6023 | 18.9 | 1.6 |
| IS-10 | Inland undulating heath mountain plain | 338 | 2975 | 9.3 | 0.8 |
| IS-8 | Inland undulating plain with boreal heath below the forest line | 264 | 2403 | 7.5 | 0.7 |
| IS-4 | Inland undulating plain below the forest line with settlements/infrastructure and agriculture | 215 | 2088 | 6.6 | 0.6 |
| IS-17 | Inland undulating plain with boreal heath below the forest line with wetlands | 148 | 1521 | 4.8 | 0.4 |
| IS-3 | Inland undulating plain below the forest line with settlements/infrastructure | 133 | 1422 | 4.5 | 0.4 |
| IS-11 | Inland undulating barren heath mountain plain | 156 | 1246 | 3.9 | 0.3 |
| IS-18 | Inland undulating heath mountain plain with wetlands | 81 | 675 | 2.1 | 0.2 |
| IS-22 | Coast-near undulating plain below the forest line with settlements/infrastructure and agriculture | 74 | 559 | 1.8 | 0.2 |
| IS | Other coastal plains (26 minor types)* | 326 | 3,159 | 9.9 | 0.9 |

\* See Appendix Table S9.

### Quantification of landscape diversity

We quantified landscape diversity for the entire mapping area by calculating the number of landscape types (richness), the number of spatial landscape units of each type (abundance) and the proportion covered by each type within the total mapping area. We described major spatial patterns of distribution of major and minor landscape types by using terminology (names) of well-known geographical areas, administrative units and bioclimatic gradients (see Bakkestuen et al. 2008).

### Validation

We compared the theoretical number of types estimated from the sample of observation units (Simensen, Halvorsen & Erikstad 2020) with the final results (realised combinations) obetained by landscape-type mapping using the procedures outlined in previous sections. We assessed the consistency and overall quality of the final maps by visual inspection and comparison with high-resolution imagery (orthophotos) and topographic maps (N50), specifically looking for major errors in the delineation and errors in the assignment to segments along each CLG and assignment to major and minor types.

### Software

We used Arc Map, ArcGisPro (ESRI 2018) and SAGA Gis v.2.3 (Conrad, Bechtel et al. 2015) for general geocomputation operations and R version 3.5.2 for statistical analyses (R Core Team 2018).

## Results

The main results of our study include: 1) a hierarchical landscape type system of realised landscape types in Norway (Fig. 3); 2) complete, area-covering and detailed (1:50 000) geographical maps of CLGs and landscape types (Fig. 4 and 5); 3) estimates of abundance and areal coverage for each major type and the most common minor landscape types (Table 3). Definitions of types and summary statistics for all minor types are provided in Appendix S9. Type descriptions and distribution maps for each major and minor type are provided (in Norwegian) by [NBIC](https://www.artsdatabanken.no/nin/landskap/typeinndeling) (2019; see example in Fig. 6).

**Fig. 3.**
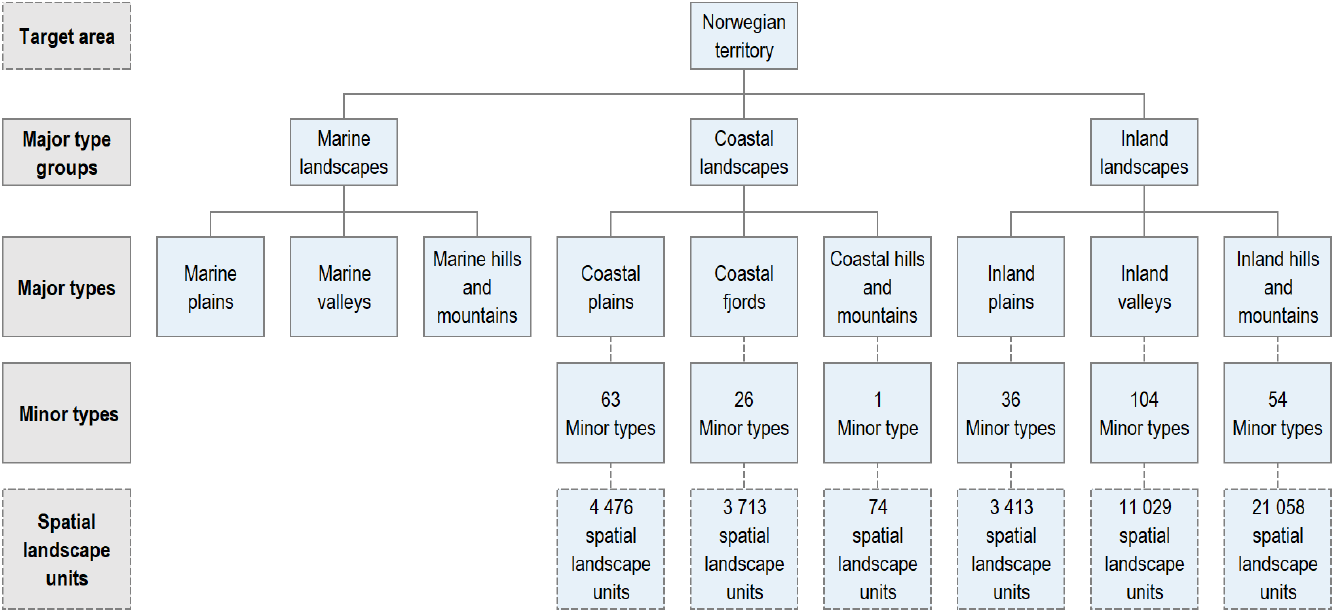
NiN type hierarchy for the landscape level of ecodiversity, with three hierarchical levels: major-type groups, major types and minor types. The minor type level is not yet developed for marine landscapes. The lowest level in the figure consists of spatial landscape units, unique areas within minor types. The numbers refer to the number of units in each category in the first version of the area-covering Norwegian landscape type map.

**Fig. 4.**
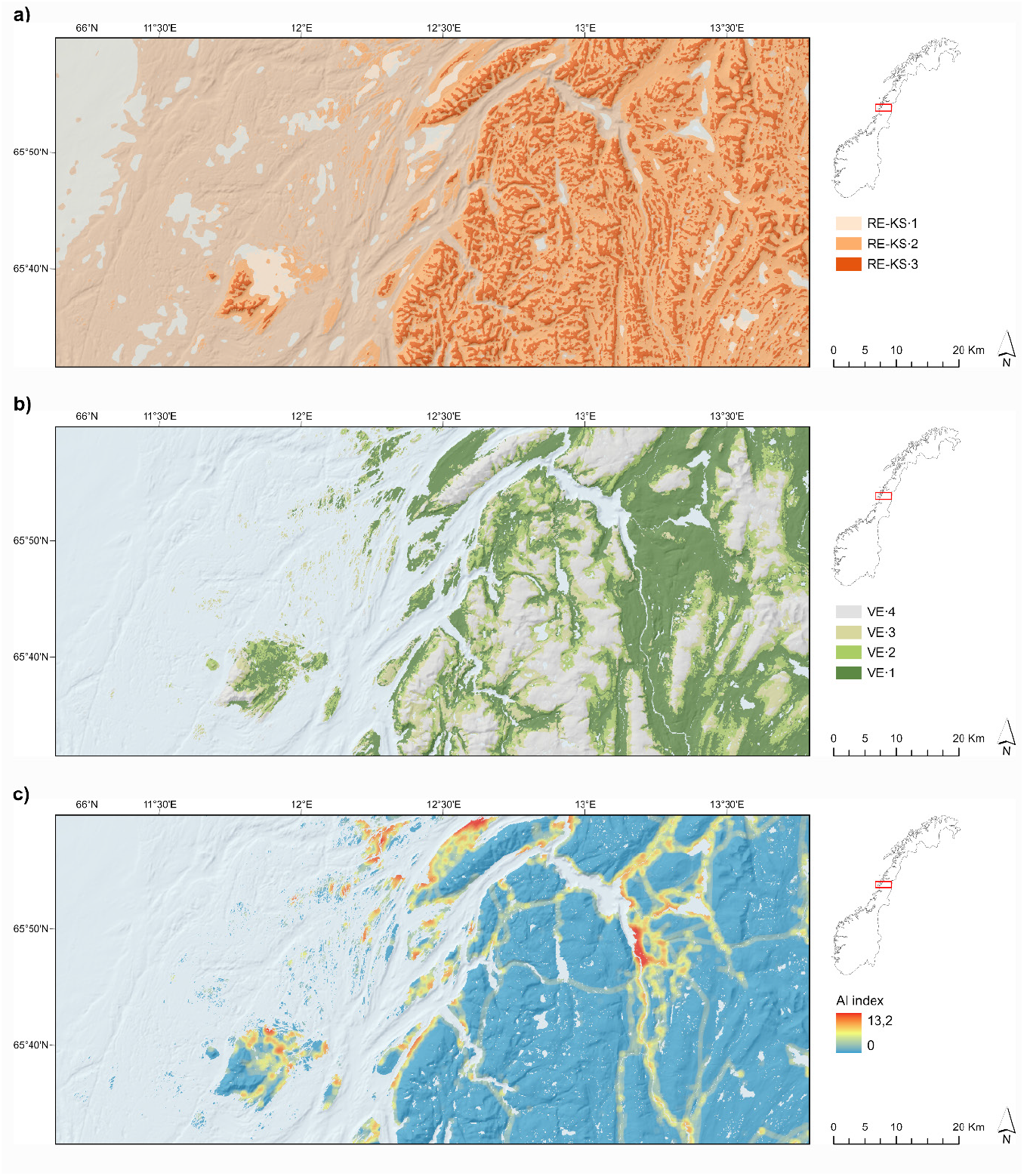
**a-c:** Examples of complex landscape gradients. **a)** The geo-ecological complex landscape gradient ‘relief in coastal plains’ (RE-KS), which expresses terrain-form variation within the coastal plains major landscape type from flat terrain to steep and rugged terrain: flat coastal plains (RE-KS·1); undulating coastal plains (RE-KS·2); rugged coastal plains (RE-KS·3). The intermediate segments are moderately protected coast (IYK·2), moderately wave-exposed coast (IYK·3); wave-exposed outer coast (IYK·4). **b)** The bio-ecological complex landscape gradient ‘vegetation cover’ (VEG), which expresses variation from forested or potentially forested areas below the climatic forest line (VEG·1) to barren mountains, without or with sparse vegetation cover (VEG·4). The intermediate segments are: boreal heath-dominated areas below the climatic forest line, kept open after logging (VEG·3); and open mountain heaths (VEG·2). **c)** The land-use related complex landscape gradient ‘land-use intensity’ (AI) from low (AI·1) and intermediate land-use intensity (AI·2) to settlement (AI·3) and city (AI·4), expressed by a continuous index (range: 0–13.2) that integrates the abundances of buildings, roads and other visible signs of human infrastructure (agricultural land-use excepted).

**Fig. 5.**
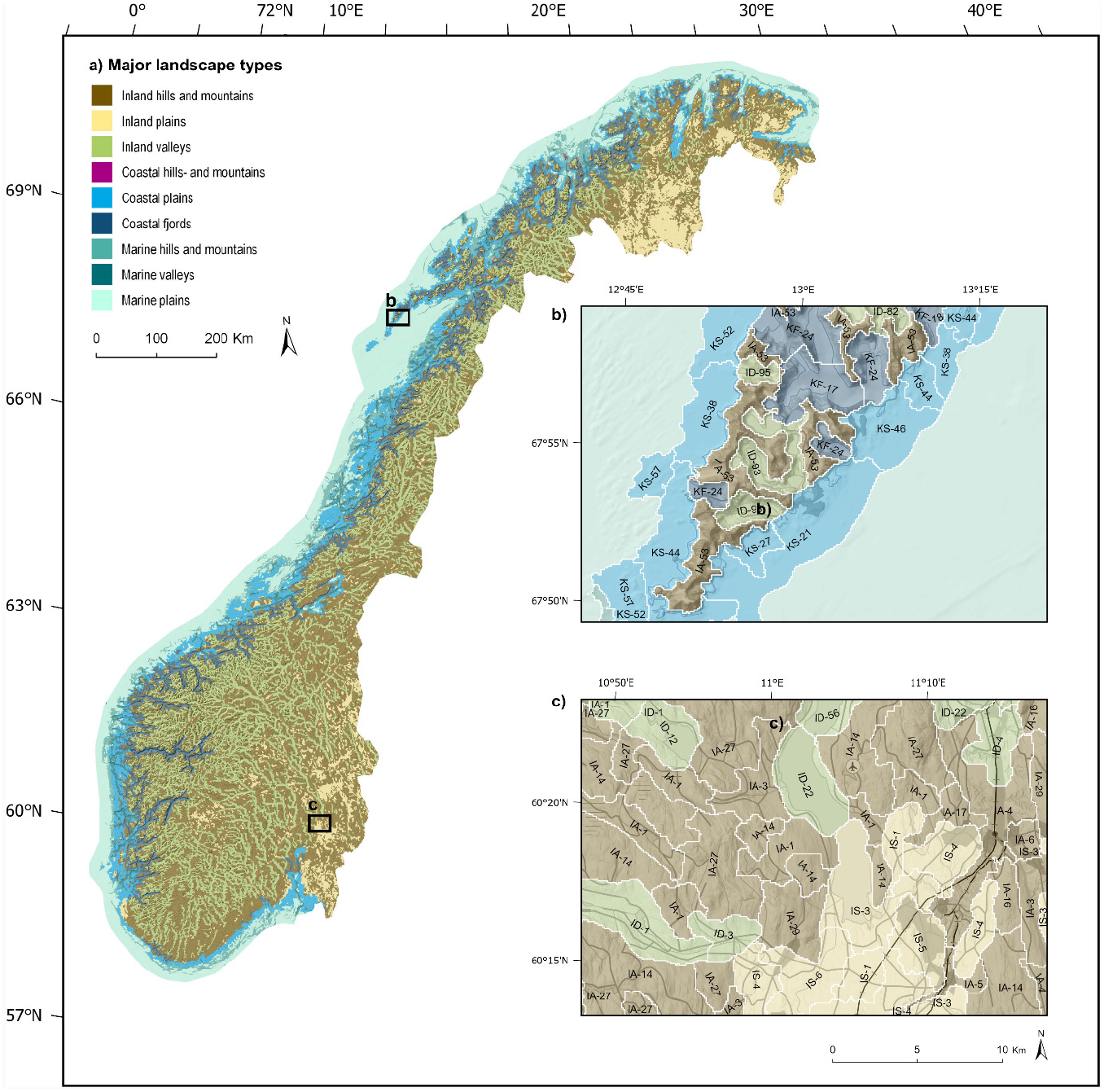
Selected examples from the NiN landscape type maps. a) Major landscape types differentiated based on broad-scale landforms and terrain variation. b–c): Examples showing: major landscape types (indicated by colour); minor landscape types (indicated by codes) and spatial landscape units (delineated by white borderlines). Minor-type codes (e.g. IA-39) are in accordance with Appendix S9. b) Coastal plains, coastal fjords, inland valleys and inland hills and mountains, Moskenes, Lofoten Islands. c) Inland plains, inland valleys and inland hills and mountains, Romerike.

**Fig. 6.**
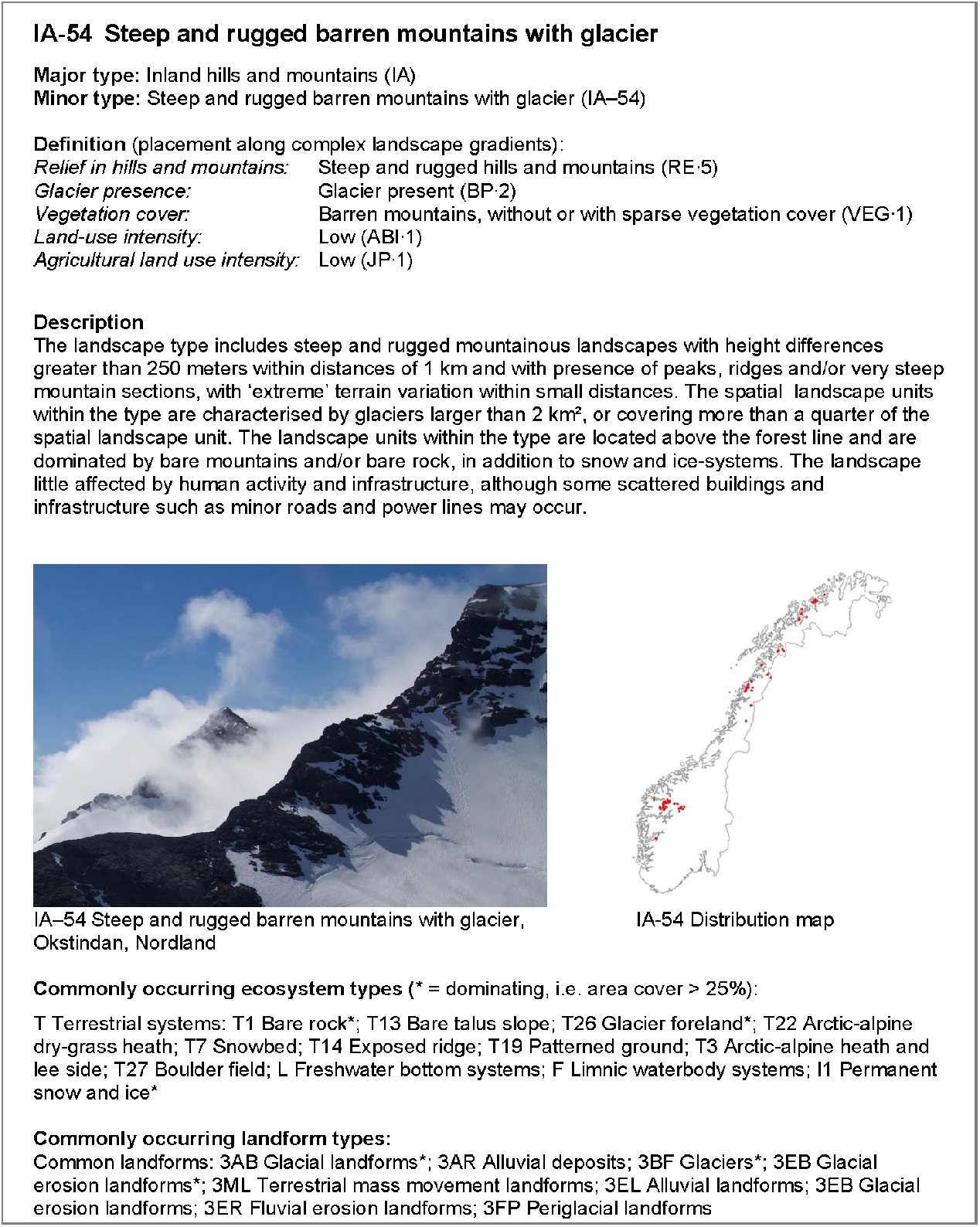
Example from the [landscape database](https://www.artsdatabanken.no/nin/LA/TI/I/A/54), for each type containing information about: major type; minor type, defining combination of segments along complex landscape gradients, a textual description of the minor type; example picture; and description of landscape elements typically occurring within the minor type, including ecosystem types and landforms.

The highest level of generalisation in the type hierarchy (Fig. 3) contained three major-type groups. Of the total target area (470,040) km^2^, inland landscapes covered 297,156 km^2^, marine landscapes covered 102,811 km^2^, and coastal landscapes covered 70,073 km^2^.

Level 2 in the type hierarchy consisted of nine major landscape types. Within coastal and inland landscapes, we identified 284 minor landscape types, defined by unique combinations of segments along the 11 CLGs, i.e. with ca. 8% dissimilarity in landscape element composition between two adjacent minor types. At the most detailed level, we identified 45,640 spatial landscape units (each 2-20 km^2^, with median area 7.4 km^2^, interquartile range 5.4–9.8 km^2^).

Fig. 7, and Fig. S9.1–S9.6 show that most of the minor landscape types were represented by < 200 spatial landscape units, while only 11 minor types contained more than 1000 spatial landscape units. All major types showed well-defined spatial patterns of distribution related to recognisable and well-known combinations of landscape features. Table 3 and Fig. 5 identify inland hills and mountains (170,606 km^2^, covering 46.5% of the area of coastal and inland landscapes) as the most common and widely distributed major landscape type throughout Norway. Minor-type variation within this major type was related to variation along the CLGs relief, vegetation cover and (agricultural) land-use intensity, resulting in 54 minor types (see Fig. 2). The most common and widely distributed minor landscape type in Norway was ‘IA-27 moderately rugged hills below the forest line’, with 2907 spatial landscape units, covering 6.1% of the total inland and coastal mapping area. The maps show a clear and well-known geographical pattern with decreasing relief in inland hills- and mountains from West to East Norway. The spatial landscape units with high relative relief (IA-RE·5) had a distinct optimum along the western coast, while units with low relief (IA-RE·1) had their optimum along the Swedish border. At a local scale, there was a significant amount of variation in this broader spatial pattern. Another evident broad-scale pattern with significant local variation is visible in the spatial distribution of the CLG vegetation cover in inland hills and mountains. At a coarser scale, the variation in vegetation cover from forest-covered areas to barren mountains (VE·1–VE·4) largely followed variation along the two major regional climatic gradients in Norway from oceanic to continental and from southern/low altitudes to northern/high altitudes. Less common minor types within inland hills and mountains occurred because of infrequent or unique combinations of specific degrees of (agricultural) land-use intensity and variation within the geo- and bio-ecological gradients. Presence of glaciers also constitute minor landscape types with relatively few (<100) spatial landscape units.

**Fig. 7.**
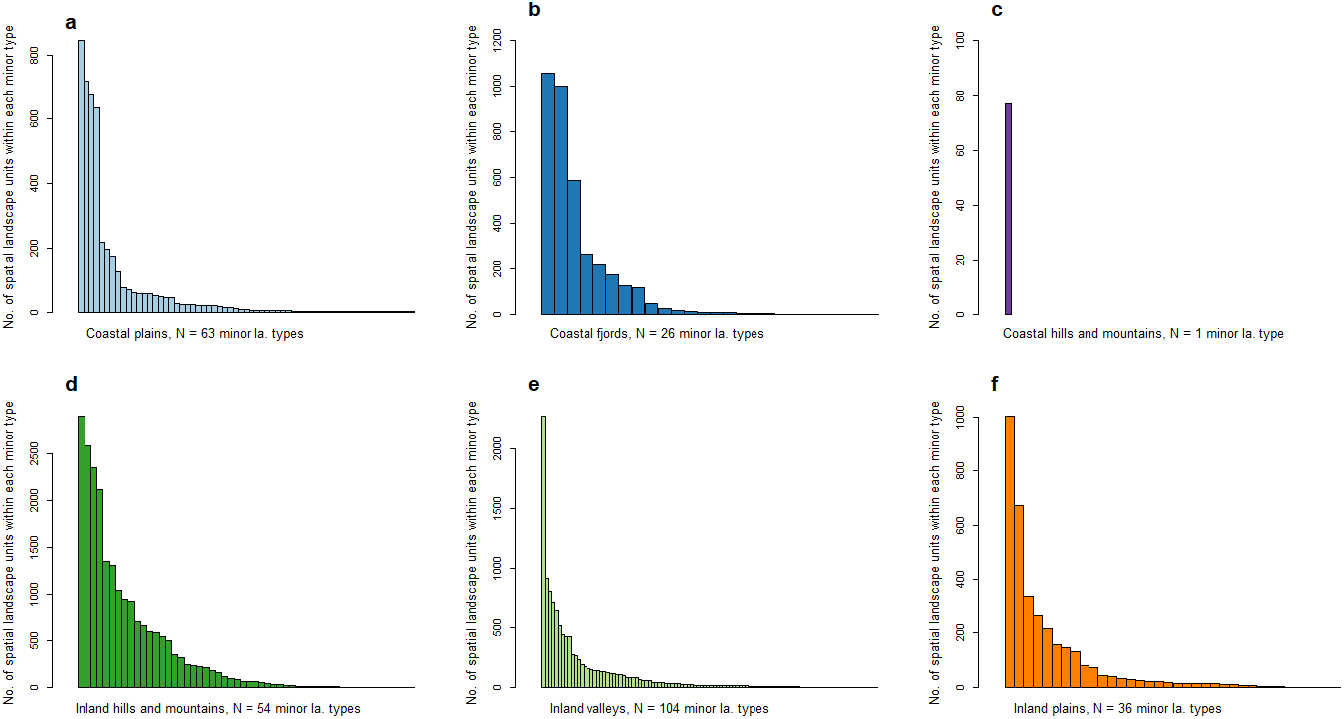
Barplots showing the number of spatial landscape units representing each of the 54 minor landscape types within: **a)** coastal plains; **b)**; coastal fjords; **c)** coastal hills and mountains; d); inland hills and mountains; **e)** inland valleys; and **f**) inland plains. The minor types are sorted by decreasing number of spatial landscape units (i.e. with one bar representing one minor landscape type, with common types to the left and rare types to the right.). The long ‘tails’ to the right show that many landscape types are ‘rare’ with unique combinations of landscape elements and properties. See also Appendix S9.1–S9.6.

Inland valleys (94,676 km^2^, 25.8% of the total inland and coastal mapping area) are widely distributed, and the major type comprises high landscape diversity, with 104 minor types, of which only 25 have more than 100 spatial landscape units. The most common minor type is ‘ID-32 open valley below the forest line’ (18,202 km^2^, 2 277 spatial landscape units). The variation in relief in valleys follow the same west-east-pattern as relief in inland hills and mountains (see examples of valley forms in Appendix S5.2). In addition to relief, vegetation cover and variation in (agricultural) land-use intensity, unique combinations of landscape properties within valleys include minor types defined by the presence of glacier and types defined by variation in hydrological properties related to lake size.

Inland plains (31,834 km^2^, 8.8% of the total inland and coastal mapping area) were frequent within several disjunct areas, otherwise rare. The most common minor type within inland plains was ‘IS-13 Inland undulating plain below the forest line with wetlands’, which covered large areas in Finnmark, Hedmark and Oppland (1005 spatial landscape units; 9763 km^2^). Near-coastal inland plains (KA·1) occurred in the fringes of the coastal plains along the coast, in the lowlands near the Oslofjord (from Østfold to Telemark), in Jæren (Rogaland), on larger islands and in lowland areas near the coast from Trøndelag to Nordland and near the coast in eastern Finnmark. Inland plains with large areas of wetlands and lakes (VP·2) were confined to the southern, middle and northern boreal zone; occurring in two distinct areas: 1) on large coastal islands in Trøndelag and Nordland (e.g. Smøla, Hitra and Frøya, Vega and Andøya); and 2) on inland plains in Finnmark and East Norway. Further variation in minor types within inland plains followed the variation in regional climate and major soil types and was reflected in variation along the CLGs vegetation cover and (agricultural) land-use intensity. Mountain plains (VE·3, VE·4, including premountain plains with boreal heath near the forest line VE·2) occurred in large continuous areas in Finnmarksvidda, the mountain plateau Hardangervidda (Hordaland, Buskerud and Telemark) and as mountain-, premountain- and forest plains (VE·1) in eastern Norway. Inland plains with high agricultural land-use intensity (JP·2) were common in lowland plains in Rogaland (Jæren) and the lowland plains in Southeastern Norway (Vestfold, Østfold, Romerike [Akershus], Toten [Oppland],Hedemarken [Hedmark], and Glåmdalen south of Elverum [Hedmark]), also associated with higher land-use intensity in general, including minor types with cities and villages (AI·3; AI·4). Distinct and unique minor types included glacier on inland plains (i.e. plateau glaciers (BP·2), e.g. Hardangerjøkulen [Hordaland], Svartisen [Nordland] and Folgefonna [Hordaland]).

Coastal plains (36249 km^2^, 10.0% of the total inland and coastal mapping area) occurred along the entire coast from Østfold to Finnmark and were particularly well developed in a more than 50 km broad zone in Helgeland, Nordland. The coastal plains encompassed a high number of minor landscape types (63) relative to the total area it covered, with variation from inner to outer coast as the most visible geographical pattern. The most common minor type was ‘KS-44 Very wave-exposed outer undulating coastal plains’. Steep and rugged coastal plains with residual mountains (‘restfjell’) had scattered occurrences along the entire coast. Coastal plains with large areas of wetlands and lakes (VP·2) were confined to large coastal island in Trøndelag and Nordland in the southern, middle and northern boreal climatic zone. Coastal plains was the major type with the highest number of spatial landscape units for urban areas. Twenty-three of the 36 spatial landscape units including larger city (64%) were located on coastal plains, while 426 of the 1126 spatial landscape units comprising village, towns and cities (37%) were located in the same major type.

Coastal fjords covered 33,478 km^2^ (9.0% of the total inland and coastal mapping area). Relatively open fjords (RE·2) was the most commonly encountered segment within the relief gradient (2231 spatial landscape units). The most common minor type was ‘KF-9 Relatively open fjord with settlements/infrastructure’, covering 9659 km^2^; 2.6% of the area within inland and coastal landscapes. Coastal fjords was the only major landscape type in which the most common minor type was characterised by a land-use intensity above low (AI·1). Six of the most common minor types in fjords had medium (AI·2) or higher land-use intensity. AI·2 typically represented a road along the fjordside with rural settlements and some human land-use. Five of the 36 spatial landscape units including city (14%) were located in fjords.

Coastal hills and mountains (36,249 km^2^, 0.1% of the total inland and coastal mapping area) were confined to coastal areas between inland hills and mountains and marine areas, not fulfilling the criteria for assignment neither to fjords nor to coastal plains. Coastal hills and mountains were most common in north-western Finnmark, else occurred scattered along the coast. Due to the low number of spatial landscape units (77) and weak statistical basis for identification of important CLGs, coastal hills and mountains were not divided into minor types.

The most common major marine landscape type in the target area was marine plains (72,948 km^2^, 71.0% of the marine mapping area). This major type encompassed deep marine plains outside the continental shelf, as well as a few marine plains within wide fjord- and coastal plain-complexes (e.g. Trondheimsfjorden, Porsangerfjorden, Tanafjorden).

Marine hills and mountains (19,493 km^2^, 19.0% of the marine mapping area) covered the continental shelf outside the coastal plains. Marine valleys (10,370 km^2^, 10.1% of the marine mapping area) contained outer parts of fjord systems, entirely submerged by water or deep valleys and gorges that cut through the continental shelf.

The number of realised minor types in the final landscape type map was 284, in contrast to 302 minor types included in the tentative type system derived from statistical analyses (see Appendix S10). We found no misclassification errors in the assignment to types by visual control, but we detected numerous minor delineation irregularities at the most detailed level in the maps (spatial landscape units). We detected minor inconsistencies in assignment to segment 2 along the CLG vegetation cover. Minor delineation irregularities were not quantified or corrected but noted for later revision by use of an improved DEM.

## Discussion

The new Norwegian landscape-type database presented here, provides, to our knowledge, the most detailed maps that simultaneously address biotic and abiotic variation at the landscape level of ecological diversity in Norway. It is also the first Norwegian map that shows marine, coastal and terrestrial landscapes in one consistent landscape-type map. Notably, the type systems obtained through the EcoSyst framework are based on systematically structured empirical evidence (see Halvorsen et al. 2020), rather than based on an expert-based, *a priori* selection of properties and components of the landscape. In contrast to the multivariate statistical analyses underpinning the landscape type system; however, the landscape-type maps presented here are spatially explicit and easy to interpret by the lay citizen. Accompanied by short textual descriptions of types, gradients and landscape elements, including pictures representative for each type (see Fig. 6), the database constitutes a publicly available knowledge base suitable for several applied purposes (see below). As such, it is also one of the first ‘ecological base-maps’ with national coverage, in a series of maps of ecosystems, landscapes and environmental variables (cf. Ministry of Climate and Environment of Norway 2015).

### Comparison with earlier landscape mapping and characterisation efforts

Our maps show well-known patterns in landscape variation, and add spatial details to earlier descriptions and coarse-scale maps of landforms and landscape gradients (see, e.g. Reusch 1905; Rudberg 1960; Moen 1999; Sulebak 2007). Yet, the unique combination of landscape elements and properties presented here have never previously been systematically combined into one type system. Moreover, NiN landscape types has a much finer spatial grain than most earlier mapping efforts with full areal coverage for Norway, such as the National Landscape Reference System (NRL; Puschmann 2005). The mean area of the spatial landscape units at the most detailed level with full areal coverage in the two systems are 8.2 km^2^ (NiN; spatial landscape units) and ~ 700 km^2^ (NRL; sub-regions), respectively. Accordingly, our study lends strong support to Strand (2011) who argue that quantification of landscape variation by use of quantitative variables and proxies for subsequent application in GIS-based mapping opens for consistent landscape-type mapping with a level of detail and at spatial extents unachievable by field-based landscape type mapping. Since NRL addresses the individual character of singular regions and NiN describes geographical phenomena in general (landscape types and gradients), the two systems complement each other and can be used in parallel whenever and wherever appropriate.

NiN landscape-type maps also differ from parametric approaches developed by use of standard sample units of constant size (i.e. by clustering 1×1 or 5×5 km grid cells, see e.g. Krøgli et al. 2015). Use of grid-based units removes the risk of delineation errors and irregularities, but fail to acknowledge important topographic features and structures, and fundamentally different landforms will inevitably be included in the same spatial unit (Bunce et al. 1996). The first phase of the pilot study in Nordland (Erikstad et al. 2015) clearly indicated that use of standardised sample units of 5×5 km is inappropriate for studying patterns in landscape element composition in regions characterised by high landform diversity and ‘steep’ gradients such as Norway.

A detailed discussion of similarities and differences between NiN landscape types and other classifications of landform types and landscape properties is provided by Simensen, Halvorsen & Erikstad (2020).

### Applications in research, planning and management

We propose a number of potential applications of the presented landscape map and the data sets on which it is based or that can be derived from it. Some of these are dealt with below.

First, the landscape-type map is a database which constitutes a resource for *basic landscape research*. There are major knowledge gaps in our understanding and ability to predict how different forms of geodiversity influence biodiversity patterns across spatial and temporal scales (Zarnetske et al. 2019). The data presented here may thus form an appropriate framework for addressing numerous research questions within landscape ecology and physical geography and may improve our knowledge of the linkages between biodiversity and geodiversity at the landscape level (see, e.g. Alahuhta et al. 2019). Landscape-type maps with full areal coverage will also open for systematic studies of the relationships between the various levels of ecological diversity (ecosystems and landscapes; Halvorsen et al. 2020). Recent studies indicate that inclusion of geodiversity and landscape variables as predictors significantly improve distribution models of ecosystem types (Simensen, Halvorsen & Erikstad 2020) as well as species (Bailey et al. 2017). Landscape-type maps also open for studies of the intimate relationship between man and nature, a classic topic in geography (Antrop & van Eetvelde 2017). Our results reveal interesting patterns of human interrelations and interactions with geomorphology (‘anthropogeomorphology’; Goudie & Viles 2010), that may be explored and quantified in greater detail, i.e. by quantifying the ‘human niche’ in the environment (cf. Xu et al. 2019). Maps showing current observable patterns in landscape composition, are also a good starting point and a reference for studies of the historical processes (the driving forces) that have caused these patterns, (see, e.g., Bürgi 2004Turner & Gardner 2015; Plieninger et al. 2016;) and landscape change trajectories (cf. Käyhkö & Skånes 2005).

Second, NiN landscape-type maps provide a framework for *monitoring of landscape changes* (Erikstad, Blumentrath, Bakkestuen, & Halvorsen, 2014). Kienast et al. (2015) and Walz et al. (2015) propose comprehensive sets of indicators for measurement of landscape changes with nation-wide application in Switzerland and Germany, respectively. The identification of CLGs opens for further studies of the relative magnitude of the drivers behind landscape change, and the response to these drivers as expressed in landscape element composition. Such knowledge may have potential importance also for future predictions of landscape change (Plieninger et al. 2016).

Third, as pointed out by several authors (e.g. van Eetvelde and Antrop, 2009; Yang et al. 2020), landscape-type maps addressing the material, observable landscape are well suited as a knowledge base for subsequent holistic *‘landscape character assessment*’ (LCA). LCA is defined as ‘a distinct, recognisable and consistent pattern of elements that make one landscape different from another, rather than better or worse’ (Swanwick 2002), or put in another way, ‘what gives an area of landscape its personality’ (Fairclough et al. 2018). Landscape-type maps may also be used in combination with methods rooted in cultural geography (see Cosgrove 2008) to address, e.g. human perception of the landscape (Tveit et al. 2006; Hunziker et al. 2008) or historical landscape development (Fairclough and Herring 2016). Combining complementary methods directed explicitly towards user needs may thus compensate for limitations and trade-offs of single methods and bridge knowledge from the natural sciences with methods from the humanities (Yang 2020).

Fourth, landscape-type maps may be well suited as a supporting tool for *conservation planning* (Beier et al. 2015). For this purpose, the introduction of explicit value criteria is necessary, since NiN landscape types, like other entities defined within the EcoSyst framework, aim at being value neutral (Halvorsen et al. 2020). The most common value criteria applied in environmental value assessments according to Erikstad et al. (2008) are rarity (existing only in small numbers and therefore valuable or interesting), representativeness (a typical example of something), diversity (a range of many people or things that are different from each other) and environmental relation/function (e.g. important landscape ecological functions such as dispersion corridors, etc). Given clear goals, such as the preservation of the total diversity of landscape types with little anthropogenic impact, landscape type maps will be a valuable resource and a good starting point for efforts towards such goals. Due to errors and uncertainties, a direct translation between e.g. rarity and ‘high conservation value’ without further analyses is nevertheless an oversimplification that must be avoided.

Finally, landscape-type maps are intended to be a useful knowledge-base for general *spatial planning and environmental impact assessment* at the local to regional administrative levels. Erikstad et al. (2020) have shown that the landscape-type maps can be successfully applied in assessing cumulative effects, which is vital for environmental impact assessments (EIA) and strategic environmental assessments (SEA). Furthermore, the spatial grain of the landscape-type maps make them well suited as a spatial framework for the development of landscape policies (specific measures aimed at the protection, management and planning of landscapes); and for setting ‘landscape quality objectives’ (i.e. agreed-upon goals for the development a particular landscape, see Council of Europe 2000). Unbiased, systematically structured information about the composition, structure and function of landscapes is also the first step towards land-use policies to assure sustainability and resilience for our ‘everyday landscapes’ (Kremen & Merenlender 2018). In this context, landscape-type maps may also provide a spatial framework suitable as a tool in informatic citizen participation and policy-making (Conrad, Cassar et al. 2011), or as a tool for negotiating landscape values (Solecka 2018).

### Errors, uncertainties, limitations and possible improvements of the mapping procedure

The type system and the maps presented here are models, that is abstract representations of real landscape variation based on simplifying assumptions. Like all other spatial models, they are affected by errors and uncertainties in the input variables in the analysis (Rød 2015). Despite intense efforts to make landscape analysis as observer-independent as possible, our semi-automated method still relies on several subjective choices made during the analytic procedure (see Yang 2020). Although these choices are thoroughly documented, and hence transparent and repeatable, further development of the mapping procedures should aim to reduce the need for manual interpretations throughout the process. Development of streamlined and standardised scripts, including geocomputation for landscape analysis and mapping within the EcoSyst framework, may speed up the documentation process, increase repeatability and facilitate applicability in other geographical settings.

An advantage of the EcoSyst framework is that each minor type (theoretically) represent a fixed amount of landscape variation, in our case, 8% compositional turnover between two adjacent types). How fine-grained the ideal type system should be is discussed in Simensen, Halvorsen & Erikstad (2020). As indicated by Fig. 7 and Fig. S9.1–9.6, most of the minor landscape types contain < 200 spatial landscape units, while only 11 minor types are represented by more than 1000 spatial landscape units. Many of the infrequent (‘rare’) landscape types comprise variation along ‘extreme segments’ introduced in the mapping procedure, mostly related to high agricultural land-use intensity or presence of a larger city. The subjective introduction of these ‘extreme segments’ (see methods section) may thus have led to an inflation of the number of minor types. On the other hand, geographical inspection confirms that these types represent ‘real’ (i.e. observable) landscape variation, e.g. landscapes dominated by distinct features such as large cities, glaciers, etc. in unique combinations with other landscape elements. Whether they should be singled out as separate types or incorporated in related types, should be discussed based upon analysis of a larger data set.

Visual inspection of the final maps also indicated inconsistencies and irregularities in the delineation of the spatial landscape units, the most detailed level in the hierarchy (delineation errors, cf. Eriksen et al. 2019). We suggest two possible reasons for delineation errors: 1) varying quality of the digital elevation model applied in the analyses (e.g. lack of bathymetry data); and 2) imperfect algorithms for delineation of spatial landscape units, especially within fjords, valleys and plains. Development of more robust delineation methods, applied on new elevation data derived from LIDAR technology (Norwegian Mapping Authority 2020) will therefore be of high priority in a possible next version of the landscape type map.

We are confident that future landscape analyses will benefit from inclusion of variables that were not available at the time the analysis presented here, was performed. Inclusion of representative data from the level below the landscape level in the NiN-system [i.e. presence of, and the fractional area occupied by, major NiN ecosystem types (cf. Bryn et al. 2018)], will represent a significant improvement. Furthermore, inclusion of landscape elements and landscape properties related to historical land use (Fairclough & Herring 2016) will broaden the scope of the type system. Better data from marine mapping programs (MAREANO; Buhl-Mortensen et al. 2015) will open for development of marine landscape type maps also at the minor-type level (see, e.g. Harris et al. 2014).

#### Conclusion

Research questions in geography and landscape ecology require interdisciplinary answers from several scales (Estes et al. 2018). In the study presented here, we have provided a synopsis of the current knowledge of observable landscape diversity in Norway obtained from various area-covering sources of information at the landscape level. We have demonstrated that, with some adaptions, the general theoretical principles proposed by Halvorsen et al. (2016; 2020) can be operationalised as a semi-automated, spatially explicit procedure for mapping at a relatively detailed level across large regions encompassing a considerable amount of landscape variation. Despite the limitations discussed above, we maintain that the landscape map presented here and the data it contains constitute a tool with may be useful as support for a wide range of applications. Hence, the data presented here may contribute to knowledge-based policies for conservation, planning and management of the unique landscape diversity of Norway.

## Data availability

A presentation of the landscape type system including descriptions of types and gradients are available in Norwegian at Norwegian Biodiversity Information Centre: (https://www.artsdatabanken.no/nin/landskap). The new landscape type maps are available as a web map service https://arcg.is/0SiTvP Geographical data can be downloaded as a complete dataset from Geonorge: https://kartkatalog.geonorge.no/metadata/naturtyper-i-norge-landskap/77512fbd-cfc5-497a-8c41-ebaf5f736ded. Additional code and supplementary data are available from GitHub: https://github.com/Artsdatabanken/landskap-ubehandlet; and data in miscellaneous formats is available at https://data.artsdatabanken.no/Natur_i_Norge/Landskap

