## Supplementary material for "Diversity and distribution of landscape types in Norway"

**Appendix – Supplementary material**

S1. Principles for constructing the NiN type hierarchy at the landscape level

S2  Neighbourhood calculations

S3. Terrain variables (morphometry) derived from digital elevation model

S4. Identifying valleys and fjords

S5. Development of GIS-based proxies for complex landscape gradients

S6. Delineation of spatial landscape units

S7. Assignment of spatial landscape units to minor landscape types

S8. GIS-procedure; assignment of spatial landscape units to minor landscape types

S9. Quantifying landscape diversity

S10. Validation

**S1 Principles, procedures and criteria used in NiN version 2.2.0 for constructing the type hierarchy for the landscape level.**

**Basic principles and procedures**

1. The NiN (and EcoSyst) type hierarchy for the landscape level is constructed by a process that integrates identification of abstract types with operationalisation of these units as concrete spatial landscape units.
2. A maximum of three major-type groups are recognised – 'coastal landscapes' which include the coastline and the adjacent strips of sea and land; 'marine landscapes' on the seaward side and 'inland landscapes' on the landward side.
3. A small number $n$ of basic geomorphologic forms (meso-scale landforms), e.g. 'plains', 'hills and mountains' and 'fjord and valleys' are recognised, each defined by a set of explicit geomorphological criteria.
4. Landscape major-type candidates are obtained as the up to $3n$ realised combinations of 3 major type groups and $n$ meso-scale landforms. These candidates are the templates for division into landscape major types.

**Landscape major types have to satisfy the following five main criteria:**

5. Landscape major types shall be characterised and defined by distinct, meso-scale geomorphological features such as plains, hills, fjords, valleys and mountains, and be mappable by landform classification procedures based on surface geometry. Accordingly, landscape major types are nested within landscape major-type candidates.
6. A spatial unit representing a major-type candidate may be divided into two or more major types if they all satisfy the criterion that one and the same group of complex landscape gradients (CLGs) is relevant for describing the variation in landscape element composition throughout each major type. Two candidate major types thus have to possess at least one CLG in their group of important CLGs (iCLGs) not shared by the other major type.
7. Each major type shall comprise more than one landscape distance unit (LDU) of variation in landscape-element composition.

**Landscape minor types have to satisfy the following main criteria:**

8. Minor types are defined as ideal combinations of major-type specific segments along the CLGs that are important within the major-type in question. CLGs are sorted by functional category in the order (1) geo-ecological CLGs, (2) bio-ecological CLGs, and (3) CLGs related to human land-use. Within each of these categories, CLGs are ordered by importance, i.e. gradient length.
9. The areas that shall be assigned to minor type are spatial landscape units, i.e. areas obtained by segmentation of major-type polygons by a rule-based procedure by which landform, terrain properties and surface geometry are taken into account. As a basic rule, the spatial landscape units shall have a minimum size of 4 km².

*References S1*

Halvorsen, R., Skarpaas, O., Bryn, A., Bratli, H., Erikstad, L., Simensen, T. & Lieungh, E. 2020. Towards a systematics of ecodiversity: The EcoSyst framework. *Global Ecology and Biogeography*.

### S2 Neighbourhood calculations

For indexes calculated as focal statistics, we applied a neighbourhood operation that computed an output raster where the value for each output cell was a function of the values of all the input cells that were present in a specified neighbourhood around that location. Conceptually, on execution, the algorithm visits each cell in the raster and calculates the specified statistic with the identified neighbourhood. The cell for which the statistic is being calculated is referred to as the processing cell. The value of the processing cell, as well as all the cell values in the identified neighbourhood, is included in the neighbourhood statistics calculation. If not specifically mentioned, focal statistics were calculated as frequencies of raster grid cells registered in a standard raster of 81 grid cells, each 100 × 100 m, in a circular neighbourhood with radius 500 m (geoprocessing tool: 'Focal statistics'; neighbourhood: circle; radius = 5; units type = cell). If, for example, 10 grid cells possessed a specific key property, a value of the key variable of 10 or, measured as a frequency, 10/81 = 0.123, was recorded.

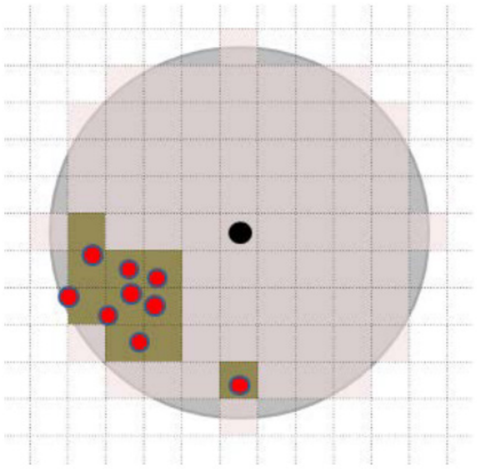

**Fig. S2.** The principle of key variable calculation as a focal statistics (frequency) with 81 standardized routes of 100 × 100 m. The 81 cells (marked with pink shadow) are wholly or partly within a circle (adjacent circle, marked with grey shadow) with a radius of 500 m around one focus point for which the key variable is calculated (marked with a black dot located in the middle of a route). In the example, the property the key variable is based on (e.g. instance of buildings) indicated by red dots. In this example, the key variable has the value 10, or alternatively 0.123 if given as frequency.

### S3 Morphometric variables applied in the calculations

Table S3. Terrain variables derived from DEM.

| Abbr. | Derived variable name | Description | Focal radius | Reference |
|---|---|---|---|---|
| TPI | Topographic position index | TPI (topographic position index) measures the difference between a cell elevation value and the average elevation of the neighbourhood around that cell. | 3000 m<br>500 m | (Gallant & Wilson 2000; Jenness 2006) |
| Tpi_ha | Extremely convex terrain | As above, Tpi values > 10. | 500 m | (Gallant & Wilson 2000; Jenness 2006) |
| VI, VD | Valley index; valley depth | Module Valley and Ridge Detection (Top Hat Approach); calculating fuzzy valley and ridge class memberships using the 'Top Hat' approach. | 3000 m<br>500 m | (Rodriguez et al. 2002) |
| | | Identification of peaks and ridges. | | (Jasiewicz & Stepinski 2013) |
| Steep_a | AA steep terrain/slope | Amount of steep terrain. | 500 m | (Gallant & Wilson 2000) |
| Rug | Terrain ruggedness VRM3, mean | Vector Ruggedness Measure (VRM) measures terrain ruggedness as the variation in three-dimensional orientation of grid cells within a neighborhood. Vector analysis is used to calculate the dispersion of vectors normal (orthogonal) to grid cells within the specified neighborhood. This method effectively captures variability in slope and aspect into a single measure. Ruggedness values in the output raster can range from 0 (no terrain variation) to 1 (complete terrain variation). Typical values for natural terrains range between 0 and about 0.4. | 500 m | (Hobson 1972; Sappington et al. 2007) |
| RR1_m | Altitudinal range | Distance between highest and lowest altitude | 500 m | (Gallant & Wilson 2000) |

*References S3*


Gallant, J.C. & Wilson, J.P. 2000. *Terrain analysis: principles and applications*. New York: Wiley.

Hobson, R. D. 1972. Surface roughness in topography: quantitative approach. Pages 221–245 in R. J. Chorley (ed.) *Spatial analysis in geomorphology*. New York: Harper and Row.

Jasiewicz, J. & Stepinski, T.F. 2013. Geomorphons — a pattern recognition approach to classification and mapping of landforms. *Geomorphology*, 182, 147-156.

Jenness, J. 2006. *Topographic position index (tpi)*. Arizona: Jenness Enterprises.

Rodriguez, F., Maire, E., Courjault-Radé, P. & Darrozes, J. 2002. The Black Top Hat function applied to a DEM: A tool to estimate recent incision in a mountainous watershed (Estibère Watershed, Central Pyrenees). *Geophysical Research Letters*, 29(6), 9-1-9-4.

Sappington, J.M., Longshore, K.M. & Thomson, D.B. 2007. Quantifiying Landscape Ruggedness for Animal Habitat Anaysis: A case Study Using Bighorn Sheep in the Mojave Desert. *Journal of Wildlife Management*, 71, 1419 -1426.

### S4. Identifying major types

*Defining valleys and fjords*

In the pilot project identification of the valleys and fjords were based on a simple index (Terrain Position Index; TPI; Gallant & Wilson 2000) calculated with a neighbourhood of 6000 metres (radius = 3000 m). This index gives negative values if the terrain is concave (pixel value less than the mean value in the neighbourhood), positive if convex. It does not, however, distinguish if the landform is a true valley or if the signal is linked to the lower part of an escarpment. We therefore changed the basic calculation using the algorithms Valley index and valley depth from the choice «Valley and Ridge Detection (Top Hat Approach)", a part of «Terrain Classification» under «Terrain analysis» in Saga GIS. The calculation was done with a neighbourhood distance of 6 km. The rest of the calculations were done in ArcMap using model builder. The first basic task was to define all depressions that have a character that define them as valleys or fjords. In the initial stage of the work this was done using the index TPI. Additionally, we used the algorithms Valley and Ridge Detection (Top Hat Approach) in the software package SAGA GIS as an alternative. We used two indices from this algorithm group calculated for the same neighbourhood as the TPI (3 km radius), "Valley Index" (VI) and "Valley Depth" (VD). The result of the SAGA GIS calculation was imported into Arc Map (10.5.1) for further analysis. The final valley and fjord result was calculated in 6 basic steps (A-F) as follows:

a) The first calculations identified areas that both have VI>0.5 and VD>100m and added up to 5 pixles (500m in cost-distance from VI>0.5 were all areas with VI<0.5 had a cost value of 100) if VD>100m.
b) The second calculation adjusted the fjord area upwards in positions to include terrestrial parts of the fjord system in cases were exceptionally deep water (typically deep fjords) affected the delineation. The adjustment was limited to: 1) areas within 1 km from the coastline; 2) areas with a negative TPI (measured over a radius of 3km and only on a terrestrial DEM (TPI_6t); 3) not in deep sea below 50m depth and not in areas with a strong signal indicating coastal plains (see below). The adjustment was limited to 750 cost distance were the areas that did not meet these requirements had the cost value of 100 and the areas within the requirements have the cost value 1.
c) The third calculation extracted the features "Peaks" (2) and "Ridges" (3) from a Geomorphon map for Norway based on the 100 m DEM. The selected features were isolated for areas within the fjord-valley model B and selected in two versions, one showing objects larger than 100 000 $m^2$ and one objects longer than 2 km.
d) The fourth step identified the ridges within the valley system that had a length over 2 km and excluded these areas from the Fjord-valley model.
e) The fifth step included selecting small and medium sized fjord/valley-polygons down to a length of 2km and within a distance of 1 km from the coast and merging them to one file.
f) The sixth and final step implicated selecting small/medium-sized valley/fjord-polygons from e that were not completely marine and merge them with large valley/fjord-polygons and medium valley/fjord-polygons with a length of more than 6000 m. This procedure is a manual procedure adjusting for different (mostly) edge effect for a small number of polygons especially in the border zone between fjords and valleys and within the coastal plains. In this process, the number of polygons was reduced from 829 to 794.

*Identifying coastal plains*

Coastal plains' were defined as landscapes with coastline (not already defined as valleys and fjords) where height differences were less than ±50 m above/below sea level (i.e. >0.5 of all cells within this interval in 1 km neighbourhoods, and limited on the inland side by a cost-distance grid). These areas were identified by an index measuring pixels in the altitude range of -50 m to + 50 m in a measuring circle with radius 500m and an index value > 0.5.

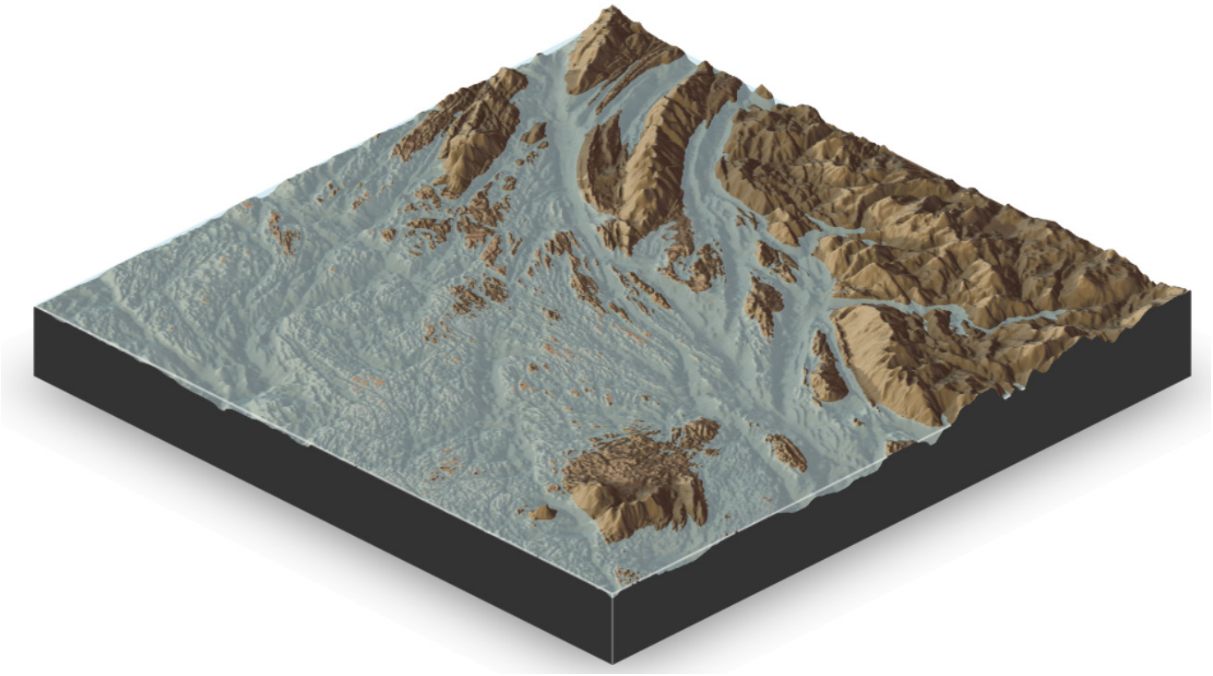

Fig. S4: coastal plains, Vega, Nordland.

*Identifying inland plains*

Inland plains' were defined as inland landscapes (without coastline) where the height differences were less than 50 m (i.e. >0.5 of all cells within this interval in 1 km neighbourhoods). These areas were identified by an index measuring pixels in the altitude range of 0 m to + 50 m in a measuring circle with radius 500m and an index value > 0.5. See also Fig. S5.2 f.

*Identifying hills- and mountains*

All remaining areas were defined as 'hills and mountains'. The major types were combined to one map, small areas (<4km²) were removed (joined with neighbours) according to the procedures in Appendix SN. Finally, hills- and mountain polygons were assigned to major-type group and major type based on their location relative to the coast-line (see above).

### S5. Development of GIS-based proxies for complex landscape gradients

#### *S5.1 Relief in inland hills and mountains (a–e) and inland plains (f)*

*a)*

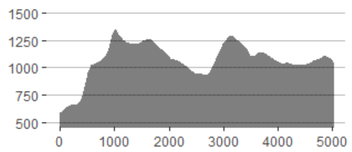


*b)*

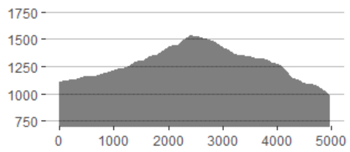


*c)*

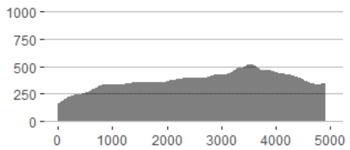


*d)*

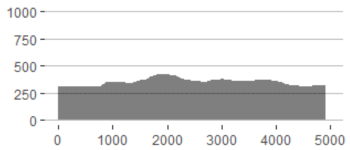


*e)*

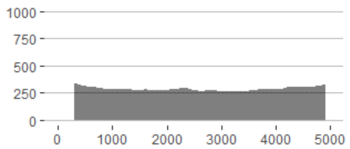


*f)*

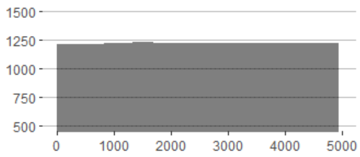


**Fig. S5.1.** Relief in hills and mountains from top: a) RE·5: Stetind, Nordland; b: RE·4: Hummelfjell, Os, Hedmark; c) RE·3: Kobberhaugene, Oslo, d) RE·2: Karsajok, Finnmark and e) RE·1: Hunndalen, Gjøvik, Oppland. Vertical exaggeration in terrain sections = 2×.

*S5.2 Relief in valleys and fjords*

Relief in valleys and fjords expresses the depth and width of the valley (or the fjord) relative to its surroundings; from wide, open valleys and fjords to narrow, deep and steep-sided valleys and fjords (see Sulebak 2007). To measure valley depth and valley width, we derived centrelines following the bottom of the valley, and constructed cross-section lines perpendicular to the centrelines. Along the centrelines and the cross-sections, we derived measurements of relief in valleys and fjords by focal calculations and raster calculations.

Assignment of spatial landscape units to segments along the relief-gradient in valleys and fjords, was done by calculating mean values for each polygon. The following threshold values were applied for stepwise division of the complex gradient: 1) open fjord/valley/valley with gentle slopes (< 0.13); 2) relatively open fjord/valley ( 0.13-0.3); 3) relatively narrow and steep-sided fjord/valley (0.3-0.5), 4) narrow and steep-sided fjord/valley (> 0.5).

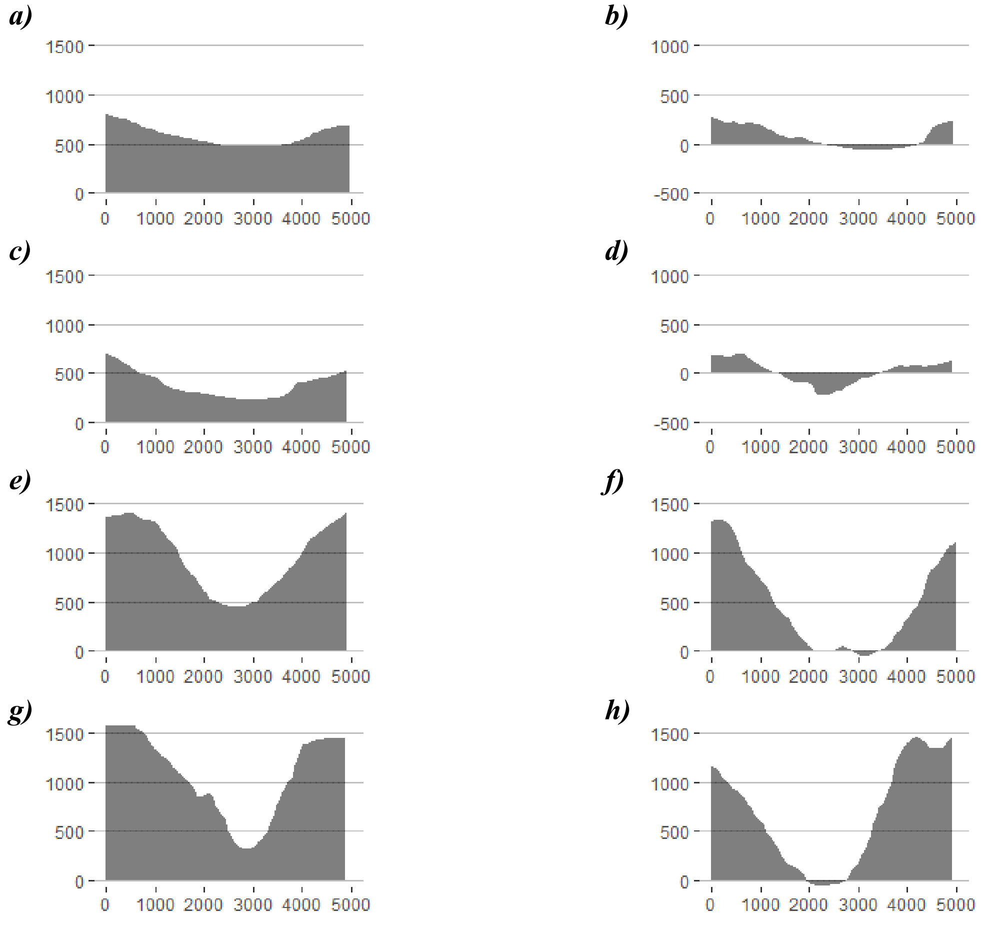


**Fig. S5.2.** examples of terrain form variation in landscape areas within valleys on the left and fjords on the right. From upper left to the lower right: a) RE·1: wide open valley, Telneset, Tynset, Hedmark; b) RE·1: wide open fjord, Repparfjorden, Finnmark; c) RE·2: open valley, Øyer, Oppland; d) RE·3": open fjord, Hvisten, Akershus; e) 3: RE·3: steep and narrow valley; Bøverdalen, Lom, Oppland, f) RE·4: steep and narrow fjord, Odda, Hordaland, g) RE·5: deep-cut narrow valley; Underdal, Sogn og Fjordane; h) RE·5: deep-cut, narrow fjord, Geiranger, Møre og Romsdal. Vertical exaggeration in terrain sections = 2×.

*S5.1 Relief in coastal plains*

Within coastal plains, terrain ruggedness (Rug3_m) and presence of extremely convex terrain (Tpi1h_a) were identified as the two variables explaining most of the terrain-form variation (Simensen et al. 2020). Vector Ruggedness Measure (VRM) measures terrain ruggedness as the variation in three-dimensional orientation of grid cells within a neighbourhood and expresses landscape variation along a gradient from flat towards rugged terrain, effectively capturing variability in slope and aspect in a single measure. Tpi1h_a (a measure of TPI1-values > 10) identifies extremely steep terrain, typically residual mountains, steep hillocks and mounds (e.g. the landform ‘rauk’).

A proxy for the complex landscape gradient was obtained by combining the two terrain-variables in a weighted raster calculation. First, the natural logarithm of the absolute values of the Rug3 raster was calculated (geoprocessing tool: raster calculator). Secondly, we applied a neighbourhood calculation with 1 km neighbourhoods (geoprocessing tool: ‘Focal statistics’; neighbourhood: circle; radius = 5; units type = cell). The values were transformed to a range from 0-1. A threshold-value of 0.62 between step one (low relief - flat coastal plains) and step two (medium-steep relief) was chosen by comparing values from the ordination analyses with spatial application of the new proxy. Step three (steep and rugged coastal plains with residual mountains, hillocks and mounds) were identified separately, by adding the values of areas within the coastal plains more than 150 m above sea level with Tpi 1 > 10. Classification of landscape areas was done by assigning all landscape areas by majority of relief classes within all pixels in the within each landscape area.

*a)*

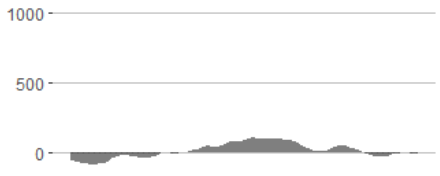


*b)*

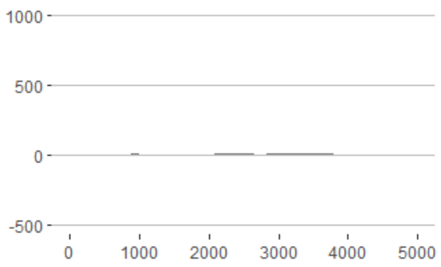


*c)*

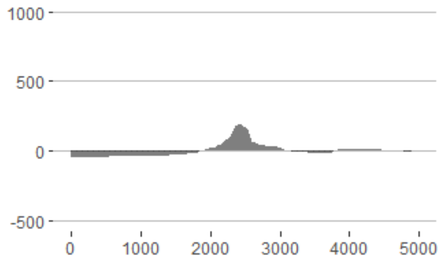


**Fig. S5.3.** examples of terrain form variation from landscape areas within coastal plains; a) Røst, Nordland: relief = 1 (low), b) Brønnøysund, Nordland; relief = 2 (medium), and c) Træna, Nordland; relief = 3 (high). Vertical exaggeration in terrain sections = 2×.

*S5.4 Inner-outer coast*

The complex landscape gradient 'inner-outer coast' was identified in the ordination analyses as the main gradient within the major type 'irregular coastal plains'. The GIS-based index for this gradient was obtained as the sum of three indices, quantified and weighted in order to project the results from the statistical analyses to entire Norwegian coast. The CLG 'Inner-outer coast' was calculated a weighted sum of three elements: 1) amount/abundance of rivers 2) island size; and 3) wave exposure.

*Amount of rivers* (R_net_a ) represents the inland element of the index, since the size of drainage basins and the amount of rivers increase towards the inland from the outer coast. R_net_a was calculated by focal statistics, within a 6 km neighbourhood (geoprocessing tool: = 'focal statistis'; neigbourhood = circle, radius = 30, units type = cell), and divided by number of land pixels. The maximum value was 0.45. Hence, we reversed the variable at the value 0.5 and multiplied by 2 (geoprocessing tool: = 'raster calculator'); by use of the expression:

$$R\_net\_ny2 \quad = 2 \cdot (0.5 - R\_net\_ny), \tag{1}$$

where '*R_net_ny2*' and '*R_net_ny*' represented the raster layers in the calculation.

Inverse island size, '*Oyst_i*', represented the middle part of the gradient (in a geographical sense) between inland and outer coast). Larger islands have 'inland properties' in landscape element composistion, whereas small islands, islets and skerries have typical coastal properties. The results of the landscape analyses (Simensen et al. 2020) indicated that this gradient should be divided in to 5 major-type-adjusted segments and that the distribution of observation units along the axis (i.e. the complex gradient) is even. We created a pixel-based proxy for this element with values between 0-1. The values were continuous and log-linear for each segment according to table S5.4.

**Table S5.4** Island size values corresponding to the threshold values for division into segments along the complex landscape gradient inner–outer coast.

| Segment | Oyst_i values | Back-transformation (island size in km$^2$) | Value after transformation |
|---|---|---|---|
| 4–5 | 0.80 | 1 | 0.8 |
| 3–4 | 0.72 | 16 | 0.6 |
| 2–3 | 0.58 | 200 | 0.4 |
| 1–2 | 0.30 | Border: island - mainland | 0; value for largest island is set to 0.2 |

The minimum value of an island was set to 1 m$^2$, corresponding to the value 1 in this part of the index. The variable was created by a logarithm transformation of the island size in m, and subsequently by the creation of a continuous variable, log-linear within each segment. The resulting values were:

Mainland: Oyst_ny = 0
(1-2): largest island (Hinnøy, 2201212711 m$^2$: 0.2; ln (2 201 212 711) = 21.512
(2-3) Island 200 km$^2$: 0.4; ln (200,000,000) = 19.114
(3-4) Island 16 km$^2$: 0.6; ln (16,000,000) = 16.588
(4-5) Island 1 km$^2$: 0.8; ln (1 000 000) = 13.816
Maximum value: Island 1 m$^2$: 1; ln (1) = 0

From these values, we applied the following transformations:

Island larger than 200 km²: Oyst_ny = 0.4 – 0.2* (ln A – 19.114)/2.398
Island 16–200 km²: Oyst_ny = 0.6 – 0.2* (ln A – 16.588)/2.526
Island 1–16 km2: Oyst_ny = 0.8 – 0.2* (ln A – 13.816)/2.772
Island less than 1 km²: Oyst_ny = 1 – 0.2* (ln A)/13.816

We calculated Island size (*Oyst_ny*) for all 113 334 islands in Norway within the national N50 database, and assigned island size values (*Oyst_ny* ) to coastal pixels by zonal statistics. All pixels on the mainland were given the value 0.

The 'wave exposure' element of the inner-outer-coast index, representing the outer part of the gradient was derived from the wave index model at the Norwegian coast (Isæus 2004). In the Isæus-index, wave exposure was modelled, with a spatial resolution of 25 m, as an index using data on fetch (distance to nearest shore, island or coast), averaged wind speed and wind frequency (estimated as the amount of time that the wind came from one of 16 direction). According to Isæus, data on wind speed and wind direction were delivered by the Norwegian Meteorological Institute and averaged over a 10-year period (i.e. 1995-2004). The model was run using the program WaveImpact based on the method 'Simplified Wave Model' (SWM) developed and described by (Isæus 2004). The model has been run by NIVA for the whole Norwegian coast and has been used e.g. as part of the habitat modelling of the National program for mapping biodiversity – coast (Bekkby et al. 2013). Since the ESWM is logarithmic, we used ln(ESWM). The maximum value for ESWM in 100 × 100 m pixels for the whole of Norway is 287,6250; ln (max) = 14.872, ln (min) = 4.779. The new derived variable for the wave exposure element 'Ekspv_ny' was calculated with values from 0-1 by the formulae:

$$Ekspv\_ny = \frac{ln\ (ESWM)\ - ln\ (ESWM\ min)}{ln\ (ESWM\ max) - \ ln\ (ESWM\ min)} \qquad (2)$$

We derived the key variable as the maximum value of the index within neighborhoods of 1 km radius (geoprocessing tool: = 'focal statistics'; neighbourhood: circle; radius = 5; units type = cell).

We derived the final proxy for the landscape gradient 'inner-outer coast' by combining the three elements in the complex landcape gradient. This was done with an arithmetic raster calculation (geoprocessing tool: 'raster calculator') on the three new derived proxies by the formula:

$$IYK\ = 0.049\ (Ekspv\_ny) +\ 0.036\ (Oyst\_ny) -\ 0.087\ (R\_net\_ny) \qquad (3)$$

where '*Ekspve_ny*' was the new derived proxy for wave-exposure: '*Oyst_ny*' was the new derived proxy for inverse island size, and '*R_net_ny*' was the new derived proxy for amount/abundance of rivers. The resulting values were applied to all pixels including coast-line, while the values for areas without coast-line was removed (geoprocessing tool: 'raster calculator'). A spatial application of the gradient is illustrated in Fig. S5.4.

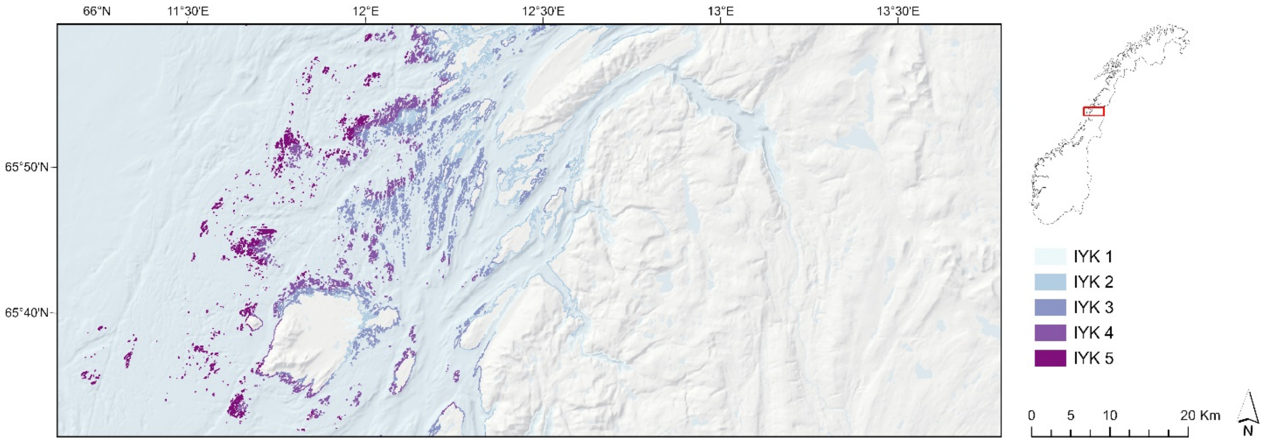


**Fig. S5.4.** The complex landscape gradient 'inner-outer coast', shown for an area within Nordland county.

#### *S5.5 Distance to coast*

The complex landscape gradient 'distance to coast' was identified within the major type 'inland plains'. The gradient separates landscape areas within inland plains near the coast (but without coastline) from other inland plains. The proxy for the complex landscape gradient was obtained by initial reclassification of land cover data (AR50) into two classes; 1) ocean areas (value = 1) and 2) terrestrial areas (value = 0) ; followed by a calculation of Euclidian distance to nearest ocean pixel within each terrestrial pixel. (geoprocessing tools: 'Reclassify'; 'euclidean distance' [input feature source data: 'ocean pixels']. The resulting raster included distance to coast for all terrestrial land pixels. In order to reproduce the results from ordination analyses, a cut-of value of 5 km was chosen as the threshold value for 'coast-near landscapes.

**Table S5.5.** Classification scheme for the complex landscape gradient 'distance to coast'.

| Class | Distance to coast | Definition |
|---|---|---|
| 1 | < 5 km | Irregular inland plains in coastal areas (but without coastline) |
| 2 | > 5 km | Inland plains more than 5 km from the coastline |

#### *S5.6 Abundance of lakes*

'Abundance of lakes' is a complex landscape gradient applied within the major type 'inland valleys'. Size of freshwater lakes were calculated, and lakes were classified within three segments, see Table. S5.6.

**Table S5.6.** Threshold values for discrete stepwise variation within the complex landscape gradient 'freshwater lakes'.

| Freshwater lake (FL) | Definition | Examples |
|---|---|---|
| FL a | Valley without lakes or with lakes < 2 km² | Østensjøvannet, Oslo (0.33 km²), Årungen, Akreshus (1.17 km²) |
| FL b | Valley with medium sized lake 2 > km² | Lustadvatnet, Trøndelag (7. 16 km²), Maridalsvannet, Oslo (4.2 km²) |
| FL c | Valleys with large lakes/inland fjords 8 > km² | Dokkfløyvatnet, Oppland (9.4 km², Vågåvatn, Oppland (15 km²), Storsjøen, Hedmark (45) km² |

Landscape areas were subsequently designated to the steps along the gradient, based on presence in at least ten pixels (> 0.1 km²), within a landscape area or alternatively absence (< 0.1 km²).

#### *S5.7 Abundance of wetlands*

'Abundance of wetlands' (i.e. proportion of wetland and small lakes) was identified as a complex landscape gradient within the major types 'fjords', 'coastal plains' and 'inland plains'. A 'wetland index' were created to represent the complex gradient, by combining data from to data sets: frequency of mire (*Mire_a*) and the frequency of freshwater lakes (point data; *Inns_s*). Values for both variables were obtained as frequencies of presence within land pixels (focal statistics), registered in a standard raster of 81 grid cells, each 100 × 100 m, in a circular neighbourhood with radius 500 m. The final index was obtained by adding the two layers by the formulae:

$$VP\_index = freq\ (Mire\_a) + freq\ (Inns\_a) \tag{4}$$

Finally, the index was transformed into a binary index; pixels with a higher value than 0.2 were designated to the category 'High proportion of wetland and small lakes'. Assignment of spatial landscape units to segments along the CLG, was done by assigning all landscape areas with a majority of land pixels > 0.2 to the class 'high proportion of wetland and small lakes'.

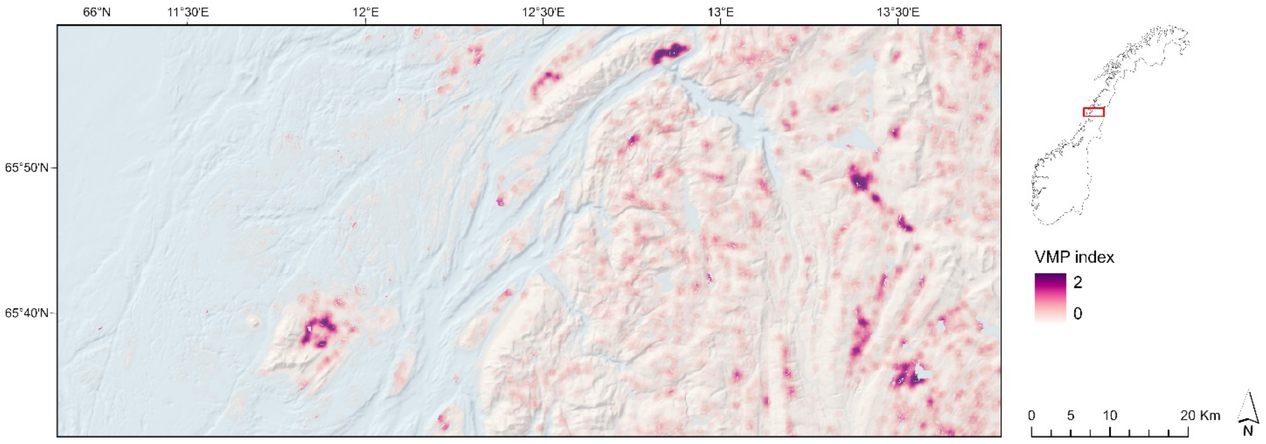


**Fig. S5.7.** The complex landscape gradient 'wetlands', shown for an area within Nordland county.

#### *S5.8 Glacier presence*

Glaciers cover 0.7% of the total land area of mainland in Norway, and due to recent estimates (Andreassen et al. 2012), there are 2534 glaciers and 2692 'glacier units' in Norway. Were glaciers being present they dominate the landscape properties and the composition of other landscape elements, e.g. resulting in absence of landscape elements normally present in similar areas. Glaciers were not identified as a complex gradient within any major types during the ordination analyses. Our interpretation was that this is due to lack of sampling units including glaciers. Because of their

dominance in the landscape where present, we subjectively included glaciers as a complex landscape gradient.
A simple ‘glacier index’ was created simply by a reclassification of AR 50 land-cover data from to a binary map with presence and absence of glaciers. Assignment of spatial landscape units to segments along this CLG was done by assigning all landscape areas with more than 2 km² and/or more 25% of the pixels within the polygon to the class ‘presence of glacier, while all other areas were designated to the class ‘absence of glacier’.

#### *S5.9 Vegetation cover*

‘Vegetation cover’, i.e. ‘potential vegetation/forest cover’, was identified as a complex landscape gradient within the major types ‘inland plains’, ‘valleys’ and ‘hills and mountains’. The GIS-proxy for the gradient was obtained by a reclassification of land cover data from NIBR, combined with a model of the forest line in Norway, and areas with boreal heaths (Bryn et al. 2013). We applied different reclassification rules above and below the forest line, according to Table S5.9. Classifications was done by standards raster operations (geoprocessing tools: ‘reclassify’ and ‘raster calculator’).

**Table S5.9.** Reclassification scheme for the complex landscape gradient 'vegetation cover' derived from a reclassification of AR50, and a model of boreal heaths and forest line.

| Original values | | Reclassified values | | | | | |
|---|---|---|---|---|---|---|---|
| **Original land cover type, AR50** | | **Reclassified land cover type for areas above/outside forest line*** | | **Reclassified areas within boreal heathlands*** | | **Reclassified land cover type for areas below forest line*** | |
| Value | Description** | New value | Description | New value | Description | New value | Description |
| 10 | Built up areas | 2 | Potential forest cover above forest line | 3 | Boreal heaths | 4 | Potential forest cover |
| 24 | Fully cultivated land | 2 | Vegetation covered areas above forest line | 3 | Boreal heaths | 4 | Potential forest cover |
| 25 | Surface cultivated land | 2 | Vegetation covered areas above forest line | 3 | Boreal heaths | 4 | Potential forest cover |
| 20 | Unspecified agriculture | 2 | Vegetation covered areas above forest line | 3 | Boreal heaths | 4 | Potential forest cover |
| 31 | Coniferous forest | NA | Not applicable (NA) | NA | – | 4 | Potential forest cover |
| 32 | Broad leaved forest | NA | – | NA | – | 4 | Potential forest cover |
| 33 | Mixed forest | NA | – | NA | – | 4 | Potential forest cover |
| 30 | Unspecified forest | NA | – | NA | – | 4 | Potential forest cover |
| 51 | Not vegetated | 1 | Barren | 3 | Boreal heaths | 1 | Barren |
| 52 | Sparse vegetation | 1 | Barren | 3 | Boreal heaths | 1 | Barren |
| 53 | Lichen | 2 | Vegetation covered areas above forest line | 3 | Boreal heaths | 2 | Vegetation covered areas above forest line |
| 54 | Intermediate vegetation above forest line | 2 | Vegetation covered areas above forest line | 3 | Boreal heaths | 2 | Vegetation covered areas above forest line |
| 55 | Vigorous vegetation above forest line | 2 | Vegetation covered areas above forest line | 3 | Boreal heaths | 2 | Vegetation covered areas above forest line |
| 50 | Unspecified open areas above forest line | 2 | Vegetation covered areas above forest line | 3 | Boreal heaths | 2 | Vegetation covered areas above forest line |
| 60 | Peat bogs | 2 | Vegetation covered areas above forest line | 3 | Boreal heaths | 4 | Potential forest cover |
| 70 | Snow/Ice | 1 | Barren | NA | – | NA | – |
| 81 | Freshwater | NA | – | NA | – | NA | – |
| 82 | Ocean water | NA | – | NA | – | NA | – |
| 99 | Not mapped | NA | – | NA | – | NA | – |

*(Bryn et al. 2013), **See (Aune-Lundberg and Strand 2010) for definitions

#### *S5.10 Land use-intensity*

*Land use intensity*

The complex landscape gradient ‘land use intensity’ was identified in the ordination analyses as a main gradient within all major types, except within coastal plains. The GIS-based proxy for this gradient was obtained as the sum of three indices, quantified and weighted in order to project the results from the statistical analyses. The index consisted of two components: 1) a building component (ByI), and 2) a component (KfI) that indicates the occurrence of artificial surfaces such as built up and constructed areas. Agricultural land use such as cultivation is not included, as this was identified as an independent gradient. Extensive human land use related to e.g. grazing, reindeer husbandry, forestry and other indirect ecological disturbances, are not included in the gradient, due to lack of data. Both components (buildings and infrastructure/built up areas) are calculated as frequencies of raster grid cells registered in a standard raster of 81 grid cells, each 100 × 100 m, in a circular neighbourhood with radius 500 m. Data included in the index is given in Table S5.11.

**Table S5.10.** Components included in the infrastructure index

| Component | Presence of landscape elements within 100m grid cells | Data source |
|---|---|---|
| **Building component (ByI)** | Buildings (of any kind) | GAB |
| | Linear elements from N50 'constructions' (power lines) | N50 |
| | Linear elements from N50 'transportation' (roads and railways are included, but minor trails, tractor trails and paths are not included) | N50 |
| **Built up areas and infrastructure (KfI)** | Built up areas | N50 |
| | Urban fabric | N50 |
| | Industrial areas | N50 |
| | Airports | N50 |
| | Mine, dump and construction sites | N50 |
| | Graveyards | N50 |
| | Sport and leisure facilities (including golf courses, ski jump facilities, ski resorts, etc.) | N50 |

ByI and KfI are combined to AI by the following formula:

$$Ifl = 2 \cdot \log_2(4 + ByI) + \log_2(4 + KfI) - 3 \cdot \log_2 4 = 2 \cdot \log_2(4 + ByI) + \log_2(4 + KfI) - 6 \quad (6)$$

Note that ByI and KfI in the formula, as expressed above, are numbered as the number of cells with buildings or constructed areas (of a maximum possible number of 81). However, in practical use, ByI and KfI must be corrected for the presence of clean seas or insurers. If ByI and KfI are expressed as frequency (share of total number of routes not completely covered by sea or water), $ByI^*$ and $KfI^*$, respectively, the formula becomes:

$$Ifl = 2 \cdot \log_2(4 + 81\, ByI^*) + \log_2(4 + 81\, KfI^*) - 6 \quad (7)$$

In principle, we could have used the frequency of cells directly as a 'building index'. The downside would be that an increase of 1 cell from 60 to 61 cells with buildings would have the same effect as an increase from 1 to 2, which do not provide a relevant measurement of how buildings affects the landscape. Therefore, we used a logarithmic scale with baseline 2, which basically means that each doubling of the cell number results in an increase in the index value of 1 unit. In order for a logarithmic transformation to work in practice, a 'correction factor' must be inserted (a number added to the number of cells with buildings). The reason for this is that $\log_2 x \rightarrow -\infty$ when $x \rightarrow 0$ so that '$\log_2 0$' makes no sense. The size of the 'correction factor' $k$ affects the relative weight applied to changes in the lower and upper part of the scale. A correction factor $k = 1$ is commonly applied, so the lowest value on the transformed scale becomes 0. With $k = 1$, the difference in index value with an increase from 0 to 8 'building routes' is 3.16 units, which is about the same increase as we get when the number of 'building cells' increases from 9 to 81. This correction does not seem reasonable; the imbalance has been shifted to the other side - differences in the lower part of the scale are added too much weight. The balance between the lower and the upper part of the scale is controlled by $k$. We chose $k = 4$, which increases the index value with one unit when the number of 'building cells' increases from 0 to 4, from 4 to 12, from 12 to 28 and from 28 to 60. The minimum value of the building component of the index can be obtained when there are no cells with buildings, $\log_2 4 = 2$.

To get an index starting at 0, we must subtract this minimum value. Since the building component of the index has weight 2 and the area component, which also uses 2-logarithms with $k = 4$, has weight 1, we have to subtract $-6\ (3 \cdot \log_2 4)$ for the index's initial value to be 0. The variation in the index relative to each single component is shown in Table 1.

The index's maximum value (presence of buildings and constructed land in all cells) is 13.23. The infrastructure index is 2-logarithmic in each component. In principle, each doubling of the frequency of buildings and constructed landmarks increases the value of ByI and KfI with a constant number of units. However, the two components of the Infrastructure Index are not considered to be equally important for the land use intensity in the landscape. The presence of buildings is considered to have a stronger impact on the landscape than the presence of constructed land-cover types. This is why ByI contributes $\frac{2}{3}$ and KfI $\frac{1}{3}$ to IfI. The consequence of this choice is that the maximum value for IfI that can be obtained on the basis of KfI alone (constructed areas/infrastructure in all cells but no buildings, e.g. a major quarry) is 4.41, while the maximum value that can be obtained on the basis of occurrence of buildings alone is 8.82. Constructed areas and infrastructure, however, still contributes to the geographic representation of the proxy because a built-up area will normally have a high index value for many cells in a neighbourhood, while a building usually affect the values for only one or a maximum 4 cells (large buildings).

The presence of larger city was added to the complex landscape gradient by identifying presence (sum) of more than 13 pixels with presence of larger city (N50; category 'city' measured in 3x3 neighbourhoods).

#### *S5.11 Agricultural land use intensity*

An agricultural index was used to identify landscapes totally dominated by agriculture. The agricultural land use index was obtained as frequency of presence of the land cover type 'fully cultivated land' (focal statistics), registered in a standard raster of 81 grid cells, each 100 × 100 m, in a circular neighbourhood with radius 500 m. The index was transformed into a binary index; pixels with a higher value than 0.25 were designated to the category 'high agricultural land use intensity'. We assigned all spatial landscape units with a majority of land pixels > 0.25 to the class 'high agricultural land use intensity'.

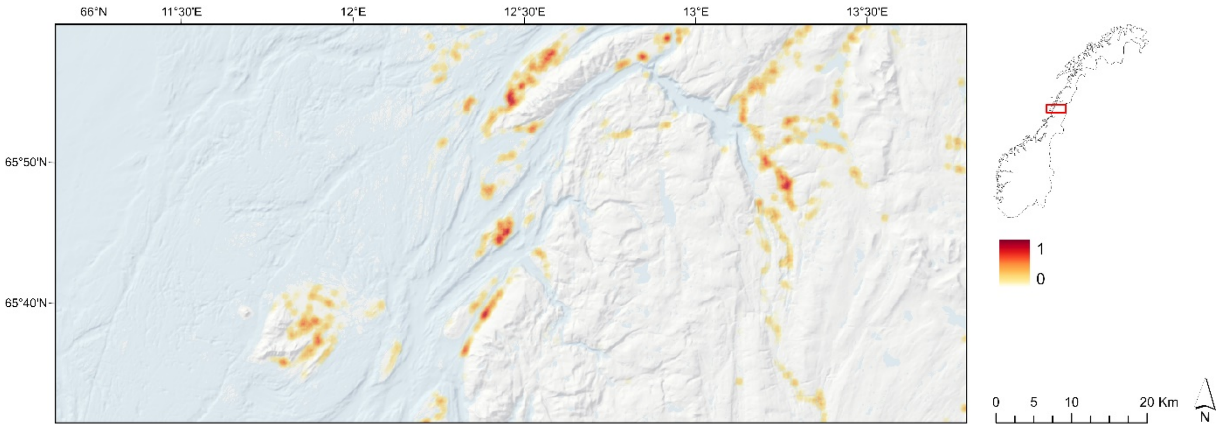


**Fig. S5.11.** The complex landscape gradient 'agricultural land use intensity', shown for an area within Nordland county. The colour scale

*References S5:*


Andreassen, L.M., Winsvold, S.H., (eds.) 2012. *Inventory of Norwegian glaciers*: Oslo: Norwegian Water Resources and Energy Directorate.

Aune-Lundberg, L. & Strand, G.-H. 2010. *CORINE land cover classes. Examination of the content of CLC classes in Norway*. Report 5/210 from Norwegian Forest and Landscape Institute.

Bekkby, T., Moy, F. E., Olsen, H., Rinde, E., Bodvin, T., Bøe, R., Steen, H., Grefsrud, E.S., Espeland, S.H., Pedersen, A., Jørgensen, N. M. 2013. The Norwegian Program for Mapping of Marine Habitats – Providing Knowledge and Maps for ICZMP. In E. Moksness; E. Dahl & J. Støttrup (eds.) *Global Challenges in Integrated Coastal Zone Management* (Vol. Vol II, pp. 21-30). Oxford: John Wiley & Sons.

Bryn, A. 2008. Recent forest limit changes in south-east Norway: Effects of climate change or regrowth after abandoned utilisation? *Norsk Geografisk Tidsskrift - Norwegian Journal of Geography*, 62(4), 251-270.

Isæus, M. 2004. *Factors structuring Fucus communities at open and complex coastlines in the Baltic Sea*. PhD thesis. Stockholm: Stockholms universitet.

### S6 Delineation of spatial landscape units

Within each major type, we delineated spatial units (landscape areas) by a rule-based division of the landscape into discrete spatial units for subsequent classification into landscape types. Since the aim of the delineation process was to identify features with shared terrain properties and landform characteristics at the given scale (4-20 km$^2$), we adapted and applied a method developed for delineation of watersheds and drainage basins. A watershed or a drainage basin is the upslope area that contributes flow – generally water – to a common outlet as concentrated drainage. Watersheds are scale-dependent and are separated by ridgelines and other breaking points in the terrain curvature (drainage divides). Applied at the appropriate scale, watershed analysis is a reliable tool in order to delineate units with shared landform characteristics, in areas with mainly concave landforms (e.g. valleys and fjords). Within areas with mainly convex landforms (e.g. hills and mountains), we applied the same procedure on an inverted DEM.

Due to lack of bathymetry for freshwater lakes (and some additional inconsistencies in the original DEM for subsea areas), a new DEM was produced for the purpose of delineating landscape areas. We added bathymetry data for 275 freshwater lakes from The Norwegian Water Resources and Energy Directorate (NVE). For all other remaining lakes larger than 2km$^2$, we isolated the existing elevation data for central areas within lakes, more than 200 meters from the shoreline (geoprocessing tool: 'buffer'; buffer distance: 200m). We assigned new elevation values to the areas in the middle of freshwater lakes by subtracting 50 meters from the elevation of the surface water level in the original DEM (geoprocessing-tool: 'raster calculator'). Finally, we interpolated elevation data for the remaining areas within the lakes and less than 200 m from the shoreline. This was done by converting the centre of each 100 m raster cell to points (geoprocessing-tool 'raster to point'), followed by interpolation of intermediate areas between the shoreline and the central lake area (geoprocessing tool: inverse distance weighted (IDW); default settings). New interpolated bathymetry data for freshwater lakes were finally added to the original DEM by replacing DEM values for all lakes larger than 2 km$^2$.

The drainage basins were created by locating the pour points at the edges of the analysis window (where water would pour out of the raster), as well as sinks, then identifying the contributing area above each pour point. This results in a raster of drainage basins/watersheds. The drainage basins were delineated by use of the geoprocessing tools 'basin' and 'watershed' according to table S6. The 'basin' tool was used as the standard procedure, while the 'watershed' tool was applied in areas with inconsistencies in the DEM due to lack of bathymetry data, allowing for detailed control over the parameters in the calculation of the catchment areas for the basins.

**Table S6.** Delineation procedures for spatial landscape units

| Major landscape type | Material | Delineation procedure |
|---|---|---|
| Coastal plains | Inverted DEM | Basin |
| Coastal hills and mountains | DEM | Basin |
| Coastal hills and mountains (concave areas) | DEM (inverted DEM in concave areas) | Watershed |
| Fjords | DEM | Watershed |
| Inland valleys | DEM | Watershed |
| Inland hills and mountains | Inverted DEM | Basin |
| Inland irregular plains | Inverted DEM | Basin |

Both basins and watersheds were derived from a flow direction raster which identifies the flow direction from every pixel, in order to find all sets of connected cells that belonged to the same drainage basin (geoprocessing tool: ‘flow direction’; input surface raster = DEM; ‘force all cells to flow outward’= enabled; ‘flow direction type’ = D8). The geoprocessing tool: ‘basin’ was applied with the flow direction raster as the input value and default settings.

In order to apply the ‘watershed’ tool, gaps in the DEM were first identified (geoprocessing tool: ‘sink’, settings = default), followed by the creation of a ‘depression-less’ DEM (geoprocessing tool: ‘fill’, applying the z-limit-values of gaps first identified by the sink tool). To provide the source locations for determination of contributing area, a ‘flow accumulation raster’ was created (geoprocessing tool: ‘flow accumulation’; input = flow direction raster; other settings = default). When the threshold values for a flow accumulation raster is used to define a watershed, the pour points for the watershed will be the junctions of a stream network derived from flow accumulation. A classified flow accumulation raster (the minimum number of cells that constitute a stream) was thus created with the settings ‘classify’, and applying ‘500’ and ‘2’ as the threshold values for two classes (geoprocessing tool: ‘raster calculator’; expression: ‘[SetNull ("FlowAcc_Flow2" < 500, 1)]’). The geoprocessing tool ‘stream link’ was then applied to assign unique values to river section between junctions, and creating a flow (threshold) accumulation raster.  Finally, the geoprocessing tool ‘watershed’ was applied, with the ‘flow-direction raster’ and the ‘threshold accumulation raster’ as the input parameters.

Watersheds (within the major types valleys and fjords) and basins (within all other major types) was clipped by removing areas outside each major type. Within each major type, the watersheds/basins were successively aggregated into landscape areas by merging adjacent polygons smaller than the minimum size of a landscape area (4km$^2$; geoprocessing tool: ‘eliminate’; input layer = the relevant watershed/basin layer clipped by major type; eliminating polygons by border = enabled).

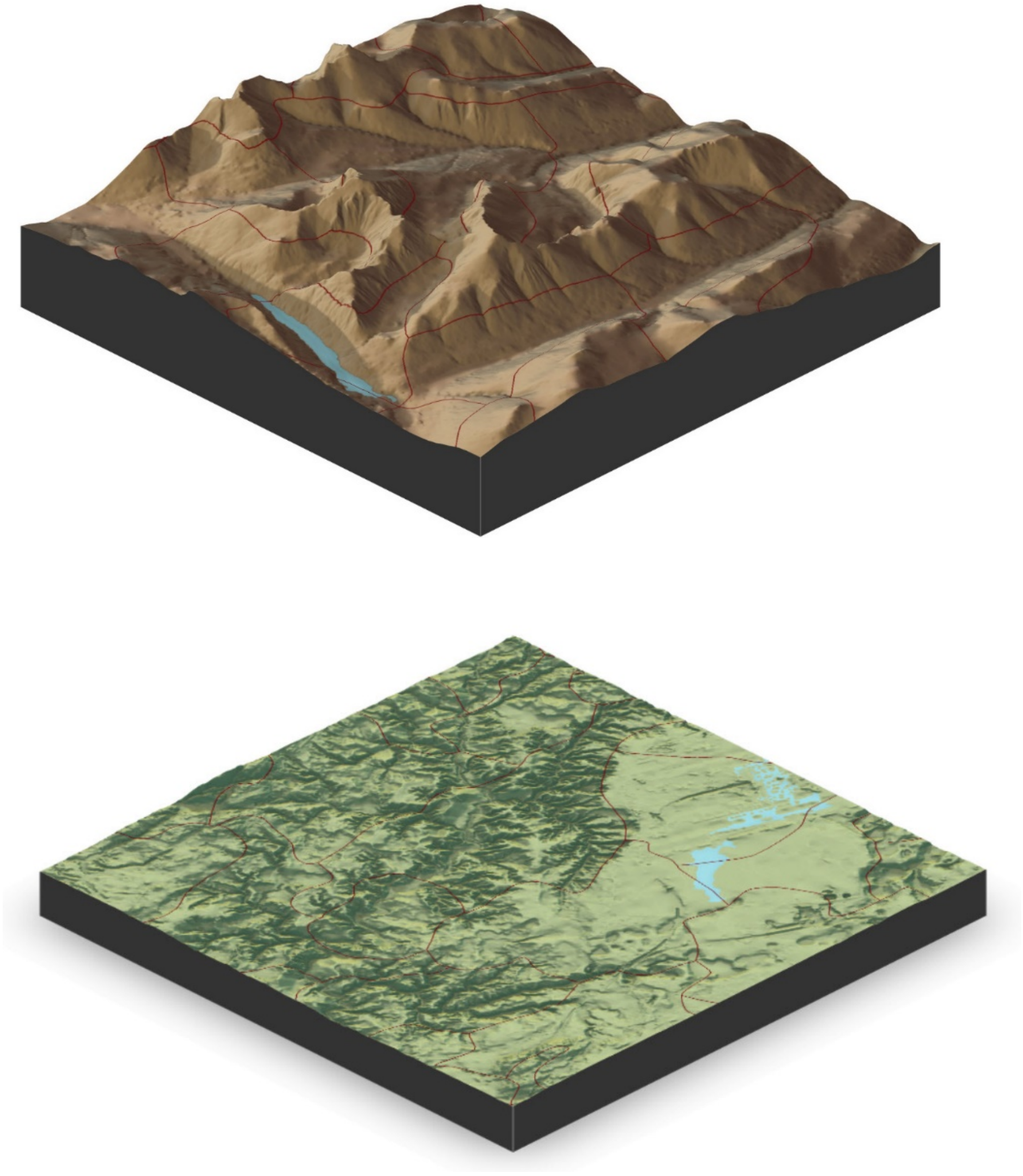

**Fig. S6.** a) 3D-models illustrating delineation of landscape units with 2× vertical exaggeration. Borders between spatial landscape units in red colour. a) example from a landscape dominated by the major types ‘inland valleys’ and ‘inland hills and mountains’ (Rondane). b) Example from inland plains (Gardermoen).

### S7 assignment of spatial units to minor landscape types

The spatial landscape units were assigned to minor types by combining steps along each CLG identified within each major type. By definition, each unique combination of gradient intervals is a minor type. The principle is illustrated in Fig. 2 in the main text. Fig. S7.1–S7.5 show type-assignment schemes for each major type. We assigned codes by use of raster values for each spatial landscape unit according to the classification schemes in Table S8.1–S8.5 below. Separate raster grids were calculated for each CLG, summarising the values for each CLG within each delineated landscape unit. Calculations were performed with the GP-tool 'Zonal statistics', and with the calculation method according to Table S8.1–S8.5 (e.g. sum, median, majority, etc.). The values were reclassified to discrete segments by applying threshold-values in accordance with the identified ecodiversity distance units in landscapes. For CLG with relatively distinct steps, landscape units were classified by presence or absence (e.g. glacier, presence of a city, etc.). 'Presence' was defined by threshold-values set by an interpretation of the analyses in Simensen et al. (2020). As the landscape units vary considerably in size, two different sets of criteria were applied; one size-criterion and one by proportion. E.g., for a landscape unit to be assigned to the segment 'presence of glacier' along the CLG GL (glacier), the polygon needed to have more than 2 km² and/or more 25% of the pixels within the polygon with presence of glacier.

In GIS, the first gradient within each major type was given values with the number of digits equal to n number of CLGs within the major type, while the second gradient was given values with n-1 digits, etc. When, e.g. the majority (i.e. > 50%) of the pixels in a polygon (landscape unit) belonged to the step RE5 within the CLC relief in hills and mountains, (i.e. steep and rugged hills and mountains), this polygon was given the value 50 000. Finally, all the CLG-rasters were added, in order to obtain CLG-codes for each delineated landscape unit (polygon), according to the principle illustrated in Box S7. The raster grids with type-codes for all polygons of each major type were transformed from raster to polygons and joined to a complete landscape type map with complete area-coverage for the mapping area. The type-codes were finally given serial numbers within each major type (e.g. IA·1, IA·2, etc.) and descriptive names based on the presence of landscape elements according to their placements along the steps within a CLG (e.g. 'steep and rugged mountains with glacier', see appendix S9).

**Box S7**. Example illustrating the process of obtaining a raster with type codes for each polygon by a simple raster calculation by adding rasters with CLG-values (integers) for the values segments along each CLG:

| | | | |
|---|---|---|---|
| IA Raster 1 | RE | | 50000 |
| IA Raster 2 | GL | + | 2000 |
| IA Raster 3 | VE | + | 400 |
| IA Raster 4 | LU | + | 10 |
| IA Raster 5 | AU | + | 1 |
| GT-Code | | | = 52411 |

| CLG Cat. | Complex landscape gradient | Coastal plains | | | |
|---|---|---|---|---|---|
| G | Inner – outer coast (IYK) | **IYK·1**: Protected inner coastal plains | **IYK·2**: Moderately wave-exposed coastal plains | **IYK·3**: Very wave-exposed outer coastal plains | **IYK·4**: Extremely wave-exposed outer coastal |
| G | Relief within coastal plains (RE-KS) | **KS-RE·1**: Low relative relief | **KS-RE·2**: Medium relative relief | **KS-RE·3**: High relative relief (including rauk/residual mountains) | |
| VP | Abundance of wetlands (VP) | **VP·1**: Low-medium proportion/abundance of wetlands and small lakes | **VP·2**: High proportion/abundance of wetlands and small lakes | | |
| L | Relief within coastal plains (RE-KS) | **AI·1-KS**: Low-medium land use intensity | **AI·2**: presence of willage/small town | **AI·3**: presence of town/city | |
| L | Agricultural land use intensity (JP) | **JP·1**: Low agricultural land use intensity | **JP·2**: High agricultural land use intensity | | |

**Fig. S7.1.** Assignment of spatial landscape units to minor landscape type within major type 'coastal plains'. Abbreviations: CLG = complex landscape gradient; Cat. = category; G = geo-ecological CLG; B = bioecological CLG; L = human land-use-related CLG.

| CLG Cat. | Complex landscape gradient | Coastal fjords | | | |
|---|---|---|---|---|---|
| G | Relief within fjords (RE-KF) | **RE-KF·1**: Open fjord with gentle slopes | **RE-KF·2**: Relatively open fjord | **RE-KF·3**: Relatively narrow and steep-sided | **RE-KF·4**: Narrow and steep-sided fjord |
| G | Abundance of wetlands (VP) | **VP·1**: Low-medium proportion/abundance of wetlands and small lakes | **VP·2**: High proportion/abundance of wetlands and small lakes | | |
| L | Land use intensity (AI) | **AI·1**: Low land use intensity | **AI·2**: medium land use intensity | **AI·3**: presence of willage/small town | **AI·4**: presence of town/city |
| L | Agricultural land use intensity (JP) | **JP·1**: Low agricultural land use intensity | **JP·2**: High agricultural land use intensity | | |

**Fig. S7.2.** Assignment of spatial landscape units to minor landscape type within major type 'coastal fjords'. Abbreviations: CLG = complex landscape gradient; Cat. = category; G = geo-ecological CLG; B = bioecological CLG; L = human land-use-related CLG.

| CLG Cat. | Complex landscape gradient | Inland valleys | | | |
|---|---|---|---|---|---|
| G | Relief within valleys (RE-ID) | **RE-ID·1**: Open valley with gentle slopes | **RE-ID·2**: Relatively open valley | **RE-ID·3**: Relatively narrow and steep-sided valley | **RE-ID·4**: Narrow and steep-sided valley |
| G | Abundance of lakes (IP) | **IP·1**: Valley without larger lakes | **IP·2**: Valley with medium sized lake | **IP·3**: Valleys with large lakes/inland fjords | |
| G | Glaciers (BP) | **BP·1**: Glacier absent | **BP·2**: Glacier present | | |
| B | Vegetation cover (VE) | **VE·1**: areas with forest cover or potential forest | **VE·2**: Boreal heaths | **VE·3**: open mountain heaths with vegetation | **VE·4**: barren mountains, without or with sparse |
| L | Land use intensity (AI) | **AI·1**: Low land use intensity | **AI·2**: medium land use intensity | AI·3: presence of willage/small town | **AI·4**: presence of town/city |
| L | Agricultural land use intensity (JP) | **JP·1**: Low agricultural land use intensity | **JP·2**: High agricultural land use intensity | | |

**Fig. S7.3.** Assignment of spatial landscape units to minor landscape type within major type ‘inland valleys’. Abbreviations: CLG = complex landscape gradient; Cat. = category; G = geo-ecological CLG; B = bioecological CLG; L = human land-use-related CLG.

| CLG Cat. | Complex landscape gradient | Inland hills and mountains | | | | |
|---|---|---|---|---|---|---|
| G | Relief within inland hills and mountains (RE-IA) | **RE-IA·1**: Depressions in inland hills and mountains | **RE-IA·2**: Undulating inland hills and mountains | **RE-IA·3**: moderately rugged inland hills and mountains | **RE-IA·4**: rugged inland hills and mountains | **RE-IA·5**: Steep and rugged inland hills and mountains |
| G | Glaciers (BP) | **BP·1**: Glacier absent | **BP·2**: Glacier present | | | |
| B | Vegetation cover (VE) | **VE·1**: areas with forest cover or potential forest cover | **VE·2**: Boreal heaths | **VE·3**: open mountain heaths with vegetation cover | **VE·4**: barren mountains, without or with sparse vegetation cover | |
| L | Land use intensity (AI) | **AI·1**: Low land use intensity | **AI·2**: medium land use intensity | **AI·3**: presence of willage/small town | **AI·4**: presence of town/city | |
| L | Agricultural land use intensity (JP) | **JP·1**: Low agricultural land use intensity | **JP·2**: High agricultural land use intensity | | | |

**Fig. S7.4.** Assignment of spatial landscape units to minor landscape type within major type ‘inland hills- and mountains’. Abbreviations: CLG = complex landscape gradient; Cat. = category; G = geo-ecological CLG; B = bioecological CLG; L = human land-use-related CLG.

<table>
<tr><th>CLG Cat.</th><th>Complex landscape gradient</th><th colspan="4">Inland plains</th></tr>
<tr><td>G</td><td>Distance to coast (KA)</td><td colspan="2">KA ·1: Coast-near inland plains</td><td colspan="2">KA ·2: Other inland plains</td></tr>
<tr><td>G</td><td>Abundance of wetlands (VP)</td><td colspan="2">VP ·1: Low-medium proportion/abundance of wetlands and small lakes</td><td colspan="2">VP ·2: Low proportion/abundance of wetlands and small lakes</td></tr>
<tr><td>G</td><td>Glaciers (BP)</td><td colspan="2">BP·1: Glacier absent</td><td colspan="2">BP·2: Glacier present</td></tr>
<tr><td>B</td><td>Vegetation cover (VE)</td><td>VE·1: areas with forest cover or potential forest</td><td>VE·2: Boreal heaths</td><td>VE·3: open mountain heaths with vegetation</td><td>VE·4: barren mountains, without or with sparse</td></tr>
<tr><td>L</td><td>Land use intensity (AI)</td><td>AI·1: Low land use intensity</td><td>AI·2: medium land use intensity</td><td>AI ·3: presence of willage/small town</td><td>AI·4: presence of town/city</td></tr>
<tr><td>L</td><td>Agricultural land use intensity (JP)</td><td colspan="2">JP·1: Low agricultural land use intensity</td><td colspan="2">JP·2: High agricultural land use intensity</td></tr>
</table>

**Fig. S7.5.** Assignment of spatial landscape units to minor landscape type within major type 'inland plains'. Abbreviations: CLG = complex landscape gradient; Cat. = category; G = geo-ecological CLG; B = bioecological CLG; L = human land-use-related CLG.

### S8 GIS-procedures for assignment of spatial landscape units to minor landscape types

**Table S8.1**. GIS-procedures for minor-type assignment, coastal plains.

| Segments along CLGs | Raster calculation code | Definition | Threshold values, original gradient (see S5) | Classification procedure for delineated polygons based on integer raster data | Ignored values in raster calculation (NODATA) |
|---|---|---|---|---|---|
| – | 100 000 | Major type: coastal plains | | | All other major types |
| **Inner–outer coast (IYK)** | | | | | |
| IYK·1 | 10 000 | Protected inner coastal plains | IYK-index < 2.4 | Majority of coastline pixels within each polygon. | All non-coastline pixels |
| IYK·2 | 20 000 | Moderately wave-exposed coastal plains | IYK-index < 2.6 | Majority of coastline pixels within each polygon. | All non-coastline pixels |
| IYK·3 | 30 000 | Very wave-exposed outer coastal plains | IYK-index < 2.815 | Majority of coastline pixels within each polygon. | All non-coastline pixels |
| IYK·4 | 40 000 | Extremely wave-exposed outer coastal plains | IYK-index > 2.815 | Majority of coastline pixels within each polygon. | All non-coastline pixels |
| **Relief (RE-KS)** | | | | | |
| RE-KS·1 | 1 000 | Low relative relief | VRM < 0.62 | Majority of pixels with relative relief = 1 | None |
| RE-KS·2 | 2 000 | Medium relative relief | VRM > 0.62 | Majority of pixels with relative relief = 2 | None |
| RE-KS·3 | 3 000 | High relative relief | TPI>10) within > 0.01 km² | Sum <1 pixel more than 150 meters above sea level and with TPI 1 > 10 | None |
| **Abundance of wetlands (VP)** | | | | | |
| VP·1 | 100 | Low-medium proportion/abundance of wetlands and small lakes | Wetland-index < 0.2 | Majority of land-pixels | Marine areas |
| VP·2 | 200 | High proportion of wetland and small lakes | Wetland-index > 0.2 | Majority of land pixels | Marine areas, and terrestrial areas less than 1 km² |
| **Land-use intensity (AI-KS)** | | | | | |
| AI-KS·1 | 10 | Low-medium land-use intensity | AI < 11,7 | Majority of land pixels within polygon | Lakes, marine areas |
| AI-KS·2 | 20 | Presence of village/small town | AI > 11.7 within > 0.3 km² | Presence (sum) of concentration of more than 30 pixels with AI > 11.7 | All other values |
| AI-KS·3 | 30 | Presence of larger city | N50 City-index > 13 within > 0.01 km² | Presence (sum) of more than 13 pixels with presence of larger city (measured in 3x3 neighbourhoods) | All other values |
| **Agricultural land-use intensity (JP)** | | | | | |
| JP·1 | 1 | Low agricultural land-use intensity | Agricultural index < 0.25 | Majority | Lakes, marine areas |
| JP·2 | 2 | High agricultural land-use intensity | Agricultural index > 0.25 | Majority | Lakes, marine areas |

**Table S8.2**. GIS-procedures for minor-type assignment, coastal fjords.

| Segments along CLGs | Raster calculation code | Definition | Threshold values, original gradient (see S5) | Classification procedure for delineated polygons based on integer raster data | Ignored values in raster calculation (NODATA) |
|---|---|---|---|---|---|
| - | 200 000 | Major type: fjords | | | All other major types |
| **Relief (RE-KF)** | | | | | |
| RE-KF·1 | 1 000 | Narrow and steep-sided fjord | < 0.13 | Mean of continuous values within each polygon | None |
| RE-KF·2 | 2 000 | Relatively narrow and steep-sided fjord | > 0.13 | Mean of continuous values within each polygon | None |
| RE-KF·3 | 3 000 | Relatively open fjord | > 0.3 | Mean of continuous values within each polygon | None |
| RE-KF·4 | 4 000 | **Open fjord with gentle slopes** | > 0.5 | Mean of continuous values within each polygon | None |
| **Abundance of wetlands (VP)** | | | | | |
| VP·1 | 100 | Low proportion of wetlands and small lakes | Wetland-index < 0.2 | Majority of land-pixels | Marine areas |
| VP·2 | 200 | High proportion of wetland and small lakes | Wetland-index > 0.2 | Majority of land pixels | Marine areas, and terrestrial areas less than 1 km² |
| **Land-use intensity (AI)** | | | | | |
| AI·1 | 10 | Low land-use intensity | AI < 6 | Less than 2 km² and less 25% of the pixels within the polygon with LU > 6 | Lakes, marine areas |
| AI·2 | 20 | Medium land-use intensity | AI > 6 | More than 2 km² and/or more 25% of the pixels within the polygon with LU > 6 | Lakes, marine areas |
| AI·3 | 30 | Presence of village/small town | AI > 11.7 within > 0.30 km² | Presence (sum) of concentration of more than 30 pixels | All other values |
| AI·4 | 40 | Presence of larger city | N50 City-index > 13 within > 0.01 km² | Presence (sum) of more than 13 pixels with presence of larger city (measured in 3x3 neighbourhoods) | All other values |
| **Agricultural land-use intensity (JP)** | | | | | |
| JP·1 | 1 | Low or medium agricultural land-use intensity | Agricultural index < 0.25 | Version A: Majority of land pixels within polygon | Lakes, marine areas |
| JP·2 | 2 | High agricultural land-use intensity | Agricultural index > 0.25 | Version A: Majority of land pixels within polygon | Lakes, marine areas |

**Table S8.3**. GIS-procedures for minor-type assignment, inland plains.

| Segments along CLGs | Raster calculation code | Definition | Threshold values, original gradient (see S5) | Classification procedure for delineated polygons based on integer raster data | Ignored values in raster calculation (NODATA) |
|---|---|---|---|---|---|
| | 300 000 | Major type: inland irregular plains | | | All other major types |
| **Distance to coast** | | | | | |
| KA·1 | 100 000 | Coast-near inland plain | < 6 km | Majority of land pixels within 6 km distance to the coast | None |
| KA·2 | 200 000 | Other inland plains | > 6 km | Majority of land pixels with more than 6 km distance to the coast | None |
| **Abundance of wetlands (VP)** | | | | | |
| VP·1 | 10 000 | Low proportion of wetlands and small lakes | Wetland-index < 0.2 | Majority of land-pixels | Lakes, marine areas |
| VP·2 | 20 000 | High proportion of wetland and small lakes | Wetland-index > 0.2 | Majority of land pixels | Lakes, marine areas |
| **Glaciers (BP)** | | | | | |
| BP·1 | 1 000 | Without glacier | | Less than 2 km² and less 25% of the pixels within the polygon with glacier | None |
| BP·2 | 2 000 | Presence of glacier | | More than 2 km² and/or more 25% of the pixels within the polygon with glacier | None |
| **Vegetation (VE)** | | | | | |
| VE·1 | 100 | Productive areas below the forest line | | Majority of class within each polygon | Lakes, marine areas |
| VE·2 | 200 | Boreal heaths | | Majority of class within each polygon | Lakes, marine areas |
| VE·3 | 300 | Mountain heaths | | Majority of class within each polygon | Lakes, marine areas |
| VE·4 | 400 | Barren mountains | | Majority of class within each polygon | Lakes, marine areas |
| **Land-use intensity (AI)** | | | | | |
| AI·1 | 10 | Low land-use intensity | AI < 6 | Majority of land pixels within polygon | Lakes, marine areas |
| AI·2 | 20 | Medium land-use intensity | AI > 6 | Majority of land pixels within polygon | Lakes, marine areas |
| AI·3 | 30 | Presence of village/small town | AI > 11.7 within > 0.30 km² | Presence (sum) of concentration of more than 30 pixels | All other values |
| AI·4 | 40 | Presence of larger city | N50 City-index > 13 within > 0.01 km² | Presence (sum) of more than 13 pixels with presence of larger city (measured in 3x3 neighbourhoods) | All other values |
| **Agricultural land-use intensity (JP)** | | | | | |
| JP·1 | 1 | Low or medium agricultural land-use intensity | Agricultural index < 0.25 | Majority of land pixels within polygon with Agricultural Index > 0.25 | Lakes, marine areas |
| JP·2 | 2 | High agricultural land-use intensity | Agricultural index > 0.25 | Majority of land pixels within polygon with agricultural index> 0.25 | Lakes, marine areas |

**Table S8.4**. GIS-procedures for minor-type assignment, inland valleys.

| Segments along CLGs | Raster calculation code | Definition | Threshold values, original gradient (see S5) | Classification procedure for delineated polygons based on integer raster data | Ignored values in raster calculation (NODATA) |
|---|---|---|---|---|---|
| – | 5 000 000 | Major type: valleys | | | All other major types |
| **Relative relief within valleys and fjords (RRFV)** | | | | | |
| RE-ID·1 | 100 000 | Open valley with gentle slopes | < 0.13 | Mean of continuous values within each polygon | None |
| RE-ID·2 | 200 000 | Relatively open valley | > 0.13 | Mean of continuous values within each polygon | None |
| RE-ID·3 | 300 000 | Relatively narrow and steep-sided valley | > 0.3 | Mean of continuous values within each polygon | None |
| RE-ID·4 | 400 000 | Narrow and steep-sided valley | > 0.5 | Mean of continuous values within each polygon | None |
| **Freshwater lake (IP)** | | | | | |
| IP·1 | 10 000 | Valley without larger lakes | < 2 km² | Absence (< 0,1 km²) | None |
| IP·2 | 20 000 | Valley with medium sized lake | 2 > km² | Presence (> 0,1km²) | None |
| IP·3 | 30 000 | Valleys with large lakes/inland fjords | 8 > km² | Presence (> 0,1 km²) | None |
| **Glacier (GL)** | | | | | |
| BP·1 | 1 000 | Without glacier | < 1 | Less than 2 km² and less 25% of the pixels within the polygon with glacier | None |
| BP·2 | 2 000 | Presence of glacier | > 1 | More than 2 km² and/or more 25% of the pixels within the polygon with glacier | None |
| **Vegetation (VE)** | | | | | |
| VE·1 | 100 | Productive areas below the forest line | | Majority of class within each polygon | |
| VE·2 | 200 | Boreal heaths | | Majority of class within each polygon | |
| VE·3 | 300 | Mountain heaths | | Majority of class within each polygon | |
| VE·4 | 400 | Barren mountains | | Majority of class within each polygon | |
| **Land-use intensity (AI)** | | | | | |
| AI·1 | 10 | Low land-use intensity | AI < 6 | Less than 2 km² and less 25% of the pixels within the polygon with LU > 6 | Lakes, marine areas |
| AI·2 | 20 | Medium land-use intensity | AI > 6 | More than 2 km² and/or more 25% of the pixels within the polygon with LU > 6 | Lakes, marine areas |
| AI·3 | 30 | Presence of village/small town | AI > 11.7 within > 0.30 km² | Presence (sum) of concentration of more than 30 pixels | All other values |
| AI·4 | 40 | Presence of larger city | N50 City-index > 13 within > 0.01 km² | Presence (sum) of more than 13 pixels with presence of larger city (measured in 3x3 neighbourhoods) | All other values |
| **Agricultural land-use intensity (JP)** | | | | | |
| JP·1 | 1 | Low or medium agricultural land-use intensity | Agricultural index < 0.25 | Agricultural Index > 0.25 for majority of land pixels within polygon | |
| JP·2 | 2 | High agricultural land-use intensity | Agricultural index > 0.25 | Agricultural Index > 0.25 for majority of land pixels within polygon | Lakes, marine areas |

**Table S8.5**. GIS-procedures for minor-type assignment, inland hills and mountains.

| Segments along CLGs | Raster calculation code | Definition | Threshold values, original gradient (see S5) | Classification procedure for delineated polygons based on integer raster data | Ignored values in raster calculation (NODATA) |
|---|---|---|---|---|---|
| – | 600 000 | Major type: hills and mountains | | | All other major types |
| **Relief, inland hills and mountains (RE-IA)** | | | | | |
| RE-IA·1 | 10 000 | Concave (valley-shaped) | TPI6<0 | Majority within each polygon | None |
| RE-IA·2 | 20 000 | Low/gentle hills and mountains | RR<100 | Majority within each polygon | None |
| RE-IA·3 | 30 000 | Medium hilly hills and mountains | 250>RR>100 | Majority within each polygon | None |
| RE-IA·4 | 30 000 | Steep hills and mountains | RR>250 | Majority within each polygon | None |
| RE-IA·5 | 40 000 | Steep hills and mountains landscapes with peaks | Mountaintop = tp1>125, hoh>300 and sink in inverse DEM | Presence > 5 pixels with 'mountain-top'. | All other values |
| **Glacier (GL)** | | | | | |
| BP·1 | 1 000 | Without glacier | < 0.25 | Less than 2 km² and less 25% of the pixels within the polygon with glacier | None |
| BP·2 | 2 000 | Presence of glacier | > 0.25 | More than 2 km² and/or more 25% of the pixels within the polygon with glacier | None |
| **Vegetation (VE)** | | | | | |
| VE·1 | 100 | Below the forest line | | Majority of the class within each polygon | |
| VE·2 | 200 | Boreal heaths | | Majority of the class within each polygon | |
| VE·3 | 300 | Mountain heaths | | Majority of the class within each polygon | |
| VE·4 | 400 | Barren mountains | | Majority of the class within each polygon | |
| **Land-use intensity (AI)** | | | | | |
| AI·1 | 10 | Low land-use intensity | AI < 6 | Less than 2 km² and less 25% of the pixels within the polygon with LU > 6 | Lakes, marine areas |
| AI·2 | 20 | Medium land-use intensity | AI > 6 | More than 2 km² and/or more 25% of the pixels within the polygon with LU > 6 | Lakes, marine areas |
| AI·3 | 30 | Presence of village/small town | AI > 11.7 within > 0.30 km² | Presence (sum) of concentration of more than 30 pixels | All other values |
| AI·4 | 40 | Presence of larger city | N50 City-index > 13 within > 0.01 km² | Presence (sum) of more than 13 pixels with presence of larger city (measured in 3x3 neighbourhoods) | All other values |
| **Agricultural land-use intensity (JP)** | | | | | |
| JP·1 | 1 | Low or medium agricultural land-use intensity | Agricultural index < 0.25 | Agricultural Index > 0.25 for majority of land pixels within polygon | |
| JP·2 | 2 | High agricultural land-use intensity | Agricultural index > 0.25 | Agricultural Index > 0.25 for the majority of land pixels within polygon | Lakes, marine areas |

**Table S8.6**. GIS-procedures for minor-type assignment, marine landscapes.

| Raster calculation code | Definition |
|---|---|
| 7 000 000 | Major type submarine coastal plains |
| 8 000 000 | Submarine fjords, < 0.1 km² terrestrial areas (10 pixels) |
| 9 000 000 | Submarine hills and mountains |
| 10 000 000 | Submarine deep-water plains |

### S9 Total area for minor landscape types

**Table S9**. Total area for all minor landscape types. Abbreviations: MiT no. = Minor landscape type number; MiT Code = minor landscape-type code; CLG segment combination = combinations of segments along complex landscape gradients (CLGs) that constitute each minor landscape type (see appendix S8); n = number of spatial landscape units within each minor landscape type.

| MiT no. | MiT Code | Minor type – descriptive name | CLG segment combination | n | Total area km² |
|---|---|---|---|---|---|
| | | **Inland hills and mountains** | | | |
| 1 | IA-1 | Depressions in hilly landscapes below the forest line | RE-IA·1&BP·1&VEG·1&AI·1&JP·1 | 2352 | 17389.2 |
| 2 | IA-2 | Depressions in hilly landscapes below the forest line with agriculture | RE-IA·1&BP·1&VEG·1&AI·1&JP·2 | 10 | 63.7 |
| 3 | IA-3 | Depressions in hilly landscapes below the forest line with settlements/infrastructure | RE-IA·1&BP·1&VEG·1&AI·2&JP·1 | 580 | 4576.8 |
| 4 | IA-4 | Depressions in hilly landscapes below the forest line with settlements/infrastructure and agriculture | RE-IA·1&BP·1&VEG·1&AI·2&JP·2 | 220 | 1498.9 |
| 5 | IA-5 | Depressions in hilly landscapes below the forest line with village/small town | RE-IA·1&BP·1&VEG·1&AI·3&JP·1 | 58 | 515.1 |
| 6 | IA-6 | Depressions in hilly landscapes below the forest line with village/small town and agriculture | RE-IA·1&BP·1&VEG·1&AI·3&JP·2 | 26 | 211.0 |
| 7 | IA-7 | Depressions in hilly landscapes below the forest line with city | RE-IA·1&BP·1&VEG·1&AI·4&JP·1 | 2 | 9.1 |
| 8 | IA-8 | Depressions in hills and mountain landscapes with boreal heath | RE-IA·1&BP·1&VEG·2&AI·1&JP·1 | 546 | 4085.8 |
| 9 | IA-9 | Depressions in hills and mountain landscapes with boreal heath and agriculture | RE-IA·1&BP·1&VEG·2&AI·1&JP·2 | 1 | 12.3 |
| 10 | IA-10 | Depressions in hills and mountain landscapes with boreal heath and settlements/infrastructure | RE-IA·1&BP·1&VEG·2&AI·2&JP·1 | 10 | 60.8 |
| 11 | IA-11 | Depressions in open heath mountain landscapes | RE-IA·1&BP·1&VEG·3&AI·1&JP·1 | 660 | 4938.9 |
| 12 | IA-12 | Depressions in barren mountain landscapes | RE-IA·1&BP·1&VEG·4&AI·1&JP·1 | 494 | 3623.1 |
| 13 | IA-13 | Depressions in barren mountain landscapes with glacier | RE-IA·1&BP·2&VEG·4&AI·1&JP·1 | 9 | 49.9 |
| 14 | IA-14 | Undulating hills below the forest line | RE-IA·2&BP·1&VEG·1&AI·1&JP·1 | 2585 | 20366.2 |
| 15 | IA-15 | Undulating hills below the forest line with agriculture | RE-IA·2&BP·1&VEG·1&AI·1&JP·2 | 4 | 28.2 |
| 16 | IA-16 | Undulating hills below the forest line with settlements/infrastructure | RE-IA·2&BP·1&VEG·1&AI·2&JP·1 | 348 | 3103.7 |
| 17 | IA-17 | Undulating hills below the forest line with settlements/infrastructure and agriculture | RE-IA·2&BP·1&VEG·1&AI·2&JP·2 | 118 | 849.8 |
| 18 | IA-18 | Undulating hills below the forest line with village/small town | RE-IA·2&BP·1&VEG·1&AI·3&JP·1 | 67 | 583.8 |
| 19 | IA-19 | Undulating hills below the forest line with village/small town and agriculture | RE-IA·2&BP·1&VEG·1&AI·3&JP·2 | 17 | 121.6 |
| 20 | IA-20 | Undulating hills below the forest line and city | RE-IA·2&BP·1&VEG·1&AI·4&JP·1 | 1 | 8.8 |
| 21 | IA-21 | Undulating hills and mountains with boreal heath | RE-IA·2&BP·1&VEG·2&AI·1&JP·1 | 593 | 4652.4 |
| 22 | IA-22 | Undulating hills and mountains with boreal heath and settlements/infrastructure | RE-IA·2&BP·1&VEG·2&AI·2&JP·1 | 7 | 46.7 |
| 23 | IA-23 | Undulating open heath mountains | RE-IA·2&BP·1&VEG·3&AI·1&JP·1 | 926 | 7464.7 |
| 24 | IA-24 | Undulating open heath mountains with settlements/infrastructure | RE-IA·2&BP·1&VEG·3&AI·2&JP·1 | 1 | 6.6 |
| 25 | IA-25 | Undulating barren mountains | RE-IA·2&BP·1&VEG·4&AI·1&JP·1 | 696 | 5902.9 |
| 26 | IA-26 | Undulating barren mountains with glacier | RE-IA·2&BP·2&VEG·4&AI·1&JP·1 | 24 | 328.6 |
| 27 | IA-27 | Moderately rugged hills below the forest line | RE-IA·3&BP·1&VEG·1&AI·1&JP·1 | 2907 | 22492.3 |
| 28 | IA-28 | Moderately rugged hills below the forest line with agriculture | RE-IA·3&BP·1&VEG·1&AI·1&JP·2 | 2 | 11.8 |
| 29 | IA-29 | Moderately rugged hills below the forest line with settlements/infrastructure | RE-IA·3&BP·1&VEG·1&AI·2&JP·1 | 237 | 2013.7 |

| MiT no. | MiT Code | Minor type – descriptive name | CLG segment combination | n | Total area km² |
|---|---|---|---|---|---|
| 30 | IA-30 | Moderately rugged hills below the forest line with settlements/infrastructure and agriculture | RE-IA·3&BP·1&VEG·1&AI·2&JP·2 | 31 | 223.1 |
| 31 | IA-31 | Moderately rugged hills below the forest line with village/small town | RE-IA·3&BP·1&VEG·1&AI·3&JP·1 | 43 | 362.9 |
| 32 | IA-32 | Moderately rugged hills below the forest line with village/small town and agriculture | RE-IA·3&BP·1&VEG·1&AI·3&JP·2 | 3 | 29.9 |
| 33 | IA-33 | Moderately rugged hills and mountains with boreal heath | RE-IA·3&BP·1&VEG·2&AI·1&JP·1 | 1302 | 10154.6 |
| 34 | IA-34 | Moderately rugged hills and mountains with boreal heath and agriculture | RE-IA·3&BP·1&VEG·2&AI·1&JP·2 | 1 | 7.3 |
| 35 | IA-35 | Moderately rugged hills and mountains with boreal heath and settlements/infrastructure | RE-IA·3&BP·1&VEG·2&AI·2&JP·1 | 10 | 90.0 |
| 36 | IA-36 | Moderately rugged open heath mountains | RE-IA·3&BP·1&VEG·3&AI·1&JP·1 | 1351 | 10867.9 |
| 37 | IA-37 | Moderately rugged open heath mountains with settlements/infrastructure | RE-IA·3&BP·1&VEG·3&AI·2&JP·1 | 2 | 28.2 |
| 38 | IA-38 | Moderately rugged barren mountains | RE-IA·3&BP·1&VEG·4&AI·1&JP·1 | 2121 | 17385.4 |
| 39 | IA-39 | Moderately rugged barren mountains with glacier | RE-IA·3&BP·2&VEG·4&AI·1&JP·1 | 67 | 828.1 |
| 40 | IA-40 | Rugged hills with forests | RE-IA·4&BP·1&VEG·1&AI·1&JP·1 | 314 | 2082.0 |
| 41 | IA-41 | Rugged hills below the forest line with settlements/infrastructure | RE-IA·4&BP·1&VEG·1&AI·2&JP·1 | 9 | 62.0 |
| 42 | IA-42 | Rugged hills below the forest line with settlements/infrastructure and agriculture | RE-IA·4&BP·1&VEG·1&AI·2&JP·2 | 1 | 3.7 |
| 43 | IA-43 | Rugged hills below the forest line with village/small town | RE-IA·4&BP·1&VEG·1&AI·3&JP·1 | 1 | 17.4 |
| 44 | IA-44 | Rugged hills and mountains with boreal heath | RE-IA·4&BP·1&VEG·2&AI·1&JP·1 | 248 | 1796.5 |
| 45 | IA-45 | Rugged open heath mountains | RE-IA·4&BP·1&VEG·3&AI·1&JP·1 | 175 | 1291.8 |
| 46 | IA-46 | Rugged barren mountains | RE-IA·4&BP·1&VEG·4&AI·1&JP·1 | 945 | 6936.8 |
| 47 | IA-47 | Rugged barren mountains with glacier | RE-IA·4&BP·2&VEG·4&AI·1&JP·1 | 54 | 438.6 |
| 48 | IA-48 | Steep and rugged hills with forests | RE-IA·5&BP·1&VEG·1&AI·1&JP·1 | 154 | 1194.6 |
| 49 | IA-49 | Steep and rugged hills below the forest line with settlements/infrastructure | RE-IA·5&BP·1&VEG·1&AI·2&JP·1 | 2 | 13.6 |
| 50 | IA-50 | Steep and rugged hills below the forest line with village/small town | RE-IA·5&BP·1&VEG·1&AI·3&JP·1 | 2 | 20.1 |
| 51 | IA-51 | Steep and rugged hills and mountains with boreal heath | RE-IA·5&BP·1&VEG·2&AI·1&JP·1 | 212 | 1597.9 |
| 52 | IA-52 | Steep and rugged open heath mountains | RE-IA·5&BP·1&VEG·3&AI·1&JP·1 | 88 | 623.2 |
| 53 | IA-53 | Steep and rugged barren mountains | RE-IA·5&BP·1&VEG·4&AI·1&JP·1 | 1038 | 8362.1 |
| 54 | IA-54 | Steep and rugged barren mountains with glacier | RE-IA·5&BP·2&VEG·4&AI·1&JP·1 | 97 | 1087.5 |
| | | **Inland valleys** | | | |
| 55 | ID-1 | Wide valley below the forest line | RED·1&IP·1&BP·1&VEG·1&AI·1&JP·1 | 913 | 7312.0 |
| 56 | ID-2 | Wide valley below the forest line with agriculture | RED·1&IP·1&BP·1&VEG·1&AI·1&JP·2 | 1 | 8.6 |
| 57 | ID-3 | Wide valley below the forest line with settlements/infrastructure | RED·1&IP·1&BP·1&VEG·1&AI·2&JP·1 | 420 | 3844.0 |
| 58 | ID-4 | Wide valley below the forest line with settlements/infrastructure and agriculture | RED·1&IP·1&BP·1&VEG·1&AI·2&JP·2 | 122 | 955.5 |
| 59 | ID-5 | Wide valley below the forest line with village/small town | RED·1&IP·1&BP·1&VEG·1&AI·3&JP·1 | 40 | 400.1 |
| 60 | ID-6 | Wide valley below the forest line with village/small town and agriculture | RED·1&IP·1&BP·1&VEG·1&AI·3&JP·2 | 12 | 131.3 |
| 61 | ID-7 | Wide valley with boreal heath below the forest line | RED·1&IP·1&BP·1&VEG·2&AI·1&JP·1 | 144 | 1203.7 |
| 62 | ID-8 | Wide valley with boreal heath below the forest line with settlements/infrastructure | RED·1&IP·1&BP·1&VEG·2&AI·2&JP·1 | 3 | 31.0 |
| 63 | ID-9 | Wide valley with heath above the forest line | RED·1&IP·1&BP·1&VEG·3&AI·1&JP·1 | 84 | 721.7 |
| 64 | ID-10 | Wide barren mountain valley | RED·1&IP·1&BP·1&VEG·4&AI·1&JP·1 | 85 | 660.0 |
| 65 | ID-11 | Wide barren mountain valley with glacier | RED·1&IP·1&BP·2&VEG·4&AI·1&JP·1 | 1 | 4.5 |
| 66 | ID-12 | Wide valley below the forest line with medium sized lakes | RED·1&IP·2&BP·1&VEG·1&AI·1&JP·1 | 170 | 1526.0 |

| MiT no. | MiT Code | Minor type – descriptive name | CLG segment combination | n | Total area km² |
|---|---|---|---|---|---|
| 67 | ID-13 | Wide valley below the forest line with medium sized lakes and agriculture | RED·1&IP·2&BP·1&VEG·1&AI·1&JP·2 | 1 | 6.8 |
| 68 | ID-14 | Wide valley below the forest line with medium sized lakes and settlements/infrastructure | RED·1&IP·2&BP·1&VEG·1&AI·2&JP·1 | 99 | 960.4 |
| 69 | ID-15 | Wide valley below the forest line with medium sized lakes and settlements/infrastructure and agriculture | RED·1&IP·2&BP·1&VEG·1&AI·2&JP·2 | 13 | 97.0 |
| 70 | ID-16 | Wide valley below the forest line with medium sized lakes and village/small town | RED·1&IP·2&BP·1&VEG·1&AI·3&JP·1 | 4 | 37.5 |
| 71 | ID-17 | Wide valley with boreal heath below the forest line with medium sized lakes | RED·1&IP·2&BP·1&VEG·2&AI·1&JP·1 | 34 | 292.1 |
| 72 | ID-18 | Wide valley with boreal heath below the forest line, medium sized lakes and settlements/infrastructure | RED·1&IP·2&BP·1&VEG·2&AI·2&JP·1 | 1 | 18.1 |
| 73 | ID-19 | Wide valley with heath above the forest line with medium sized lakes | RED·1&IP·2&BP·1&VEG·3&AI·1&JP·1 | 31 | 246.9 |
| 74 | ID-20 | Wide barren mountain valley with medium sized lakes | RED·1&IP·2&BP·1&VEG·4&AI·1&JP·1 | 27 | 232.0 |
| 75 | ID-21 | Wide valley below the forest line with inland fjord | RED·1&IP·3&BP·1&VEG·1&AI·1&JP·1 | 148 | 1492.0 |
| 76 | ID-22 | Wide valley below the forest line with inland fjord and settlements/infrastructure | RED·1&IP·3&BP·1&VEG·1&AI·2&JP·1 | 143 | 1472.3 |
| 77 | ID-23 | Wide valley below the forest line with inland fjord with settlements/infrastructure and agriculture | RED·1&IP·3&BP·1&VEG·1&AI·2&JP·2 | 9 | 54.5 |
| 78 | ID-24 | Wide valley below the forest line with inland fjord and village/small town | RED·1&IP·3&BP·1&VEG·1&AI·3&JP·1 | 13 | 182.4 |
| 79 | ID-25 | Wide valley below the forest line with inland fjord and village/small town and agriculture | RED·1&IP·3&BP·1&VEG·1&AI·3&JP·2 | 1 | 16.4 |
| 80 | ID-26 | Wide valley below the forest line with inland fjord and city | RED·1&IP·3&BP·1&VEG·1&AI·4&JP·1 | 2 | 32.8 |
| 81 | ID-27 | Wide valley with boreal heath below the forest line with inland fjord | RED·1&IP·3&BP·1&VEG·2&AI·1&JP·1 | 41 | 422.4 |
| 82 | ID-28 | Wide valley with boreal heath below the forest line with inland fjord and settlements/infrastructure | RED·1&IP·3&BP·1&VEG·2&AI·2&JP·1 | 4 | 61.6 |
| 83 | ID-29 | Wide valley with boreal heath below the forest line with inland fjord and village/small town | RED·1&IP·3&BP·1&VEG·2&AI·3&JP·1 | 1 | 5.8 |
| 84 | ID-30 | Wide valley with heath above the forest line with inland fjord | RED·1&IP·3&BP·1&VEG·3&AI·1&JP·1 | 21 | 217.7 |
| 85 | ID-31 | Wide barren mountain valley with inland fjord | RED·1&IP·3&BP·1&VEG·4&AI·1&JP·1 | 18 | 299.8 |
| 86 | ID-32 | Open valley below the forest line | RED·2&IP·1&BP·1&VEG·1&AI·1&JP·1 | 2277 | 18201.4 |
| 87 | ID-33 | Open valley below the forest line with agriculture | RED·2&IP·1&BP·1&VEG·1&AI·1&JP·2 | 3 | 22.8 |
| 88 | ID-34 | Open valley below the forest line with settlements/infrastructure | RED·2&IP·1&BP·1&VEG·1&AI·2&JP·1 | 804 | 7395.4 |
| 89 | ID-35 | Open valley below the forest line with settlements/infrastructure and agriculture | RED·2&IP·1&BP·1&VEG·1&AI·2&JP·2 | 108 | 847.8 |
| 90 | ID-36 | Open valley below the forest line with village/small town | RED·2&IP·1&BP·1&VEG·1&AI·3&JP·1 | 56 | 563.4 |
| 91 | ID-37 | Open valley below the forest line with village/small town and agriculture | RED·2&IP·1&BP·1&VEG·1&AI·3&JP·2 | 17 | 192.6 |
| 92 | ID-38 | Open valley with boreal heath below the forest line | RED·2&IP·1&BP·1&VEG·2&AI·1&JP·1 | 516 | 4209.2 |
| 93 | ID-39 | Open valley with boreal heath below the forest line with settlements/infrastructure | RED·2&IP·1&BP·1&VEG·2&AI·2&JP·1 | 6 | 66.6 |
| 94 | ID-40 | Open valley with boreal heath below the forest line with village/small town | RED·2&IP·1&BP·1&VEG·2&AI·3&JP·1 | 2 | 8.5 |
| 95 | ID-41 | Open valley with heath above the forest line | RED·2&IP·1&BP·1&VEG·3&AI·1&JP·1 | 441 | 3509.8 |
| 96 | ID-42 | Open valley with heath above the forest line with settlements/infrastructure | RED·2&IP·1&BP·1&VEG·3&AI·2&JP·1 | 4 | 28.8 |
| 97 | ID-43 | Open barren mountain valley | RED·2&IP·1&BP·1&VEG·4&AI·1&JP·1 | 639 | 5319.1 |
| 98 | ID-44 | Open barren mountain valley with glacier | RED·2&IP·1&BP·2&VEG·4&AI·1&JP·1 | 24 | 202.1 |
| 99 | ID-45 | Open valley below the forest line with medium sized lakes | RED·2&IP·2&BP·1&VEG·1&AI·1&JP·1 | 273 | 2490.4 |
| 100 | ID-46 | Open valley below the forest line with medium sized lakes and settlements/infrastructure | RED·2&IP·2&BP·1&VEG·1&AI·2&JP·1 | 109 | 1151.9 |
| 101 | ID-47 | Open valley below the forest line with medium sized lakes, settlements/infrastructure and agriculture | RED·2&IP·2&BP·1&VEG·1&AI·2&JP·2 | 16 | 112.2 |
| 102 | ID-48 | Open valley below the forest line with medium sized lakes and village/small town | RED·2&IP·2&BP·1&VEG·1&AI·3&JP·1 | 8 | 97.5 |
| 103 | ID-49 | Open valley with boreal heath below the forest line with medium sized lakes | RED·2&IP·2&BP·1&VEG·2&AI·1&JP·1 | 83 | 740.7 |

| MiT no. | MiT Code | Minor type – descriptive name | CLG segment combination | n | Total area km² |
|---|---|---|---|---|---|
| 104 | ID-50 | Open valley with boreal heath below the forest line with medium sized lakes and settlements/infrastructure | RED·2&IP·2&BP·1&VEG·2&AI·2&JP·1 | 1 | 6.6 |
| 105 | ID-51 | Open valley with heath above the forest line with medium sized lakes | RED·2&IP·2&BP·1&VEG·3&AI·1&JP·1 | 68 | 607.0 |
| 106 | ID-52 | Open barren mountain valley with medium sized lakes | RED·2&IP·2&BP·1&VEG·4&AI·1&JP·1 | 117 | 1051.9 |
| 107 | ID-53 | Open barren mountain valley with medium sized lakes and settlements/infrastructure | RED·2&IP·2&BP·1&VEG·4&AI·2&JP·1 | 1 | 41.1 |
| 108 | ID-54 | Open barren mountain valley with medium sized lakes and glacier | RED·2&IP·2&BP·2&VEG·4&AI·1&JP·1 | 5 | 39.8 |
| 109 | ID-55 | Open valley below the forest line with inland fjord | RED·2&IP·3&BP·1&VEG·1&AI·1&JP·1 | 194 | 2010.8 |
| 110 | ID-56 | Open valley below the forest line with inland fjord and settlements/infrastructure | RED·2&IP·3&BP·1&VEG·1&AI·2&JP·1 | 129 | 1602.2 |
| 111 | ID-57 | Open valley below the forest line with inland fjord and settlements/infrastructure and agriculture | RED·2&IP·3&BP·1&VEG·1&AI·2&JP·2 | 15 | 146.1 |
| 112 | ID-58 | Open valley below the forest line with inland fjord and village/small town | RED·2&IP·3&BP·1&VEG·1&AI·3&JP·1 | 12 | 255.7 |
| 113 | ID-59 | Open valley with boreal heath below the forest line with inland fjord | RED·2&IP·3&BP·1&VEG·2&AI·1&JP·1 | 39 | 388.0 |
| 114 | ID-60 | Open valley with boreal heath below the forest line with inland fjord and settlements/infrastructure | RED·2&IP·3&BP·1&VEG·2&AI·2&JP·1 | 2 | 18.1 |
| 115 | ID-61 | Open valley with heath above the forest line with inland fjord | RED·2&IP·3&BP·1&VEG·3&AI·1&JP·1 | 29 | 329.9 |
| 116 | ID-62 | Open valley with heath above the forest line with inland fjord and settlements/infrastructure | RED·2&IP·3&BP·1&VEG·3&AI·2&JP·1 | 1 | 12.0 |
| 117 | ID-63 | Open barren mountain valley with inland fjord | RED·2&IP·3&BP·1&VEG·4&AI·1&JP·1 | 35 | 334.0 |
| 118 | ID-64 | Open barren mountain valley with inland fjord and glacier | RED·2&IP·3&BP·2&VEG·4&AI·1&JP·1 | 2 | 14.7 |
| 119 | ID-65 | Narrow valley below the forest line | RED·3&IP·1&BP·1&VEG·1&AI·1&JP·1 | 718 | 5799.0 |
| 120 | ID-66 | Narrow valley below the forest line with agriculture | RED·3&IP·1&BP·1&VEG·1&AI·1&JP·2 | 3 | 13.2 |
| 121 | ID-67 | Narrow valley below the forest line with settlements/infrastructure | RED·3&IP·1&BP·1&VEG·1&AI·2&JP·1 | 128 | 1161.2 |
| 122 | ID-68 | Narrow valley below the forest line with settlements/infrastructure and agriculture | RED·3&IP·1&BP·1&VEG·1&AI·2&JP·2 | 12 | 85.3 |
| 123 | ID-69 | Narrow valley below the forest line with village/small town | RED·3&IP·1&BP·1&VEG·1&AI·3&JP·1 | 3 | 26.4 |
| 124 | ID-70 | Narrow valley with boreal heath below the forest line | RED·3&IP·1&BP·1&VEG·2&AI·1&JP·1 | 269 | 2119.6 |
| 125 | ID-71 | Narrow valley with boreal heath below the forest line with settlements/infrastructure | RED·3&IP·1&BP·1&VEG·2&AI·2&JP·1 | 3 | 14.6 |
| 126 | ID-72 | Narrow valley with heath above the forest line | RED·3&IP·1&BP·1&VEG·3&AI·1&JP·1 | 81 | 617.3 |
| 127 | ID-73 | Narrow barren mountain valley | RED·3&IP·1&BP·1&VEG·4&AI·1&JP·1 | 420 | 3250.9 |
| 128 | ID-74 | Narrow barren mountain valley with settlements/infrastructure | RED·3&IP·1&BP·1&VEG·4&AI·2&JP·1 | 1 | 14.8 |
| 129 | ID-75 | Narrow barren mountain valley with glacier | RED·3&IP·1&BP·2&VEG·4&AI·1&JP·1 | 42 | 358.1 |
| 130 | ID-76 | Narrow valley below the forest line with medium sized lakes | RED·3&IP·2&BP·1&VEG·1&AI·1&JP·1 | 60 | 576.4 |
| 131 | ID-77 | Narrow valley below the forest line with medium sized lakes and settlements/infrastructure | RED·3&IP·2&BP·1&VEG·1&AI·2&JP·1 | 13 | 106.1 |
| 132 | ID-78 | Narrow valley below the forest line with medium sized lakes and village/small town | RED·3&IP·2&BP·1&VEG·1&AI·3&JP·1 | 1 | 6.5 |
| 133 | ID-79 | Narrow valley with boreal heath below the forest line with medium sized lakes | RED·3&IP·2&BP·1&VEG·2&AI·1&JP·1 | 11 | 87.7 |
| 134 | ID-80 | Narrow valley with boreal heath below the forest line with medium sized lakes and settlements/infrastructure | RED·3&IP·2&BP·1&VEG·2&AI·2&JP·1 | 1 | 14.2 |
| 135 | ID-81 | Narrow valley with heath above the forest line with medium sized lakes | RED·3&IP·2&BP·1&VEG·3&AI·1&JP·1 | 9 | 71.8 |
| 136 | ID-82 | Narrow barren mountain valley with medium sized lakes | RED·3&IP·2&BP·1&VEG·4&AI·1&JP·1 | 29 | 245.6 |
| 137 | ID-83 | Narrow barren mountain valley with medium sized lakes and glacier | RED·3&IP·2&BP·2&VEG·4&AI·1&JP·1 | 2 | 15.7 |
| 138 | ID-84 | Narrow valley below the forest line with inland fjord | RED·3&IP·3&BP·1&VEG·1&AI·1&JP·1 | 53 | 513.5 |
| 139 | ID-85 | Narrow valley below the forest line with inland fjord and settlements/infrastructure | RED·3&IP·3&BP·1&VEG·1&AI·2&JP·1 | 13 | 173.5 |

| MiT no. | MiT Code | Minor type – descriptive name | CLG segment combination | n | Total area km² |
|---|---|---|---|---|---|
| 140 | ID-86 | Narrow valley with inland fjord and settlements/infrastructure and agriculture | RED·3&IP·3&BP·1&VEG·1&AI·2&JP·2 | 1 | 6.2 |
| 141 | ID-87 | Narrow valley with boreal heath below the forest line with inland fjord | RED·3&IP·3&BP·1&VEG·2&AI·1&JP·1 | 5 | 47.0 |
| 142 | ID-88 | Narrow valley with heath above the forest line with inland fjord | RED·3&IP·3&BP·1&VEG·3&AI·1&JP·1 | 1 | 11.7 |
| 143 | ID-89 | Narrow barren mountain valley with inland fjord | RED·3&IP·3&BP·1&VEG·4&AI·1&JP·1 | 11 | 98.5 |
| 144 | ID-90 | Deeply cut valley below the forest line | RED·4&IP·1&BP·1&VEG·1&AI·1&JP·1 | 234 | 1654.2 |
| 145 | ID-91 | Deeply cut valley below the forest line with settlements/infrastructure | RED·4&IP·1&BP·1&VEG·1&AI·2&JP·1 | 13 | 129.2 |
| 146 | ID-92 | Deeply cut valley below the forest line with village/small town | RED·4&IP·1&BP·1&VEG·1&AI·3&JP·1 | 1 | 10.2 |
| 147 | ID-93 | Deeply cut valley with boreal heath below the forest line | RED·4&IP·1&BP·1&VEG·2&AI·1&JP·1 | 112 | 711.6 |
| 148 | ID-94 | Deeply cut valley with boreal heath below the forest line with settlements/infrastructure | RED·4&IP·1&BP·1&VEG·2&AI·2&JP·1 | 1 | 2.5 |
| 149 | ID-95 | Deeply cut valley with heath above the forest line | RED·4&IP·1&BP·1&VEG·3&AI·1&JP·1 | 23 | 138.1 |
| 150 | ID-96 | Deeply cut barren mountain valley | RED·4&IP·1&BP·1&VEG·4&AI·1&JP·1 | 153 | 1091.5 |
| 151 | ID-97 | Deeply cut barren mountain valley with glacier | RED·4&IP·1&BP·2&VEG·4&AI·1&JP·1 | 13 | 126.8 |
| 152 | ID-98 | Deeply cut valley below the forest line with medium sized lakes | RED·4&IP·2&BP·1&VEG·1&AI·1&JP·1 | 11 | 96.9 |
| 153 | ID-99 | Deeply cut valley below the forest line with medium sized lakes and settlements/infrastructure | RED·4&IP·2&BP·1&VEG·1&AI·2&JP·1 | 1 | 6.9 |
| 154 | ID-100 | Deeply cut barren mountain valley with medium sized lakes | RED·4&IP·2&BP·1&VEG·4&AI·1&JP·1 | 3 | 31.8 |
| 155 | ID-101 | Deeply cut valley below the forest line with inland fjord | RED·4&IP·3&BP·1&VEG·1&AI·1&JP·1 | 11 | 140.6 |
| 156 | ID-102 | Deeply cut valley below the forest line with inland fjord and settlements/infrastructure | RED·4&IP·3&BP·1&VEG·1&AI·2&JP·1 | 1 | 13.0 |
| 157 | ID-103 | Deeply cut valley with boreal heath below the forest line with inland fjord | RED·4&IP·3&BP·1&VEG·2&AI·1&JP·1 | 4 | 35.4 |
| 158 | ID-104 | Deeply cut barren mountain valley with inland fjord | RED·4&IP·3&BP·1&VEG·4&AI·1&JP·1 | 1 | 12.5 |
| | | **Inland plains** | | | |
| 159 | IS-1 | Inland undulating plain below the forest line | DIK·1&BP·1&VP·1&VEG·1&AI·1&JP·1 | 673 | 6017.9 |
| 160 | IS-2 | Inland undulating plain below the forest line with agriculture | DIK·1&BP·1&VP·1&VEG·1&AI·1&JP·2 | 9 | 63.0 |
| 161 | IS-3 | Inland undulating plain below the forest line with settlements/infrastructure | DIK·1&BP·1&VP·1&VEG·1&AI·2&JP·1 | 133 | 1422.1 |
| 162 | IS-4 | Inland undulating plain below the forest line with settlements/infrastructure and agriculture | DIK·1&BP·1&VP·1&VEG·1&AI·2&JP·2 | 215 | 2088.5 |
| 163 | IS-5 | Inland undulating plain below the forest line with village/small town | DIK·1&BP·1&VP·1&VEG·1&AI·3&JP·1 | 24 | 271.5 |
| 164 | IS-6 | Inland undulating plain below the forest line with village/small town and agriculture | DIK·1&BP·1&VP·1&VEG·1&AI·3&JP·2 | 45 | 537.3 |
| 165 | IS-7 | Inland undulating plain below the forest line with city | DIK·1&BP·1&VP·1&VEG·1&AI·4&JP·1 | 1 | 21.7 |
| 166 | IS-8 | Inland undulating plain with boreal heath below the forest line | DIK·1&BP·1&VP·1&VEG·2&AI·1&JP·1 | 264 | 2403.6 |
| 167 | IS-9 | Inland undulating plain with boreal heath below the forest line with settlements/infrastructure | DIK·1&BP·1&VP·1&VEG·2&AI·2&JP·1 | 1 | 5.7 |
| 168 | IS-10 | Inland undulating heath mountain plain | DIK·1&BP·1&VP·1&VEG·3&AI·1&JP·1 | 338 | 2974.7 |
| 169 | IS-11 | Inland undulating barren heath mountain plain | DIK·1&BP·1&VP·1&VEG·4&AI·1&JP·1 | 156 | 1246.0 |
| 170 | IS-12 | Inland undulating barren mountain plain with glacier | DIK·1&BP·2&VP·1&VEG·4&AI·1&JP·1 | 13 | 93.1 |
| 171 | IS-13 | Inland undulating plain below the forest line with wetlands | DIK·1&BP·1&VP·2&VEG·1&AI·1&JP·1 | 1005 | 9759.5 |
| 172 | IS-14 | Inland undulating plain below the forest line with wetlands and settlements/infrastructure | DIK·1&BP·1&VP·2&VEG·1&AI·2&JP·1 | 33 | 321.5 |
| 173 | IS-15 | Inland undulating plain below the forest line with wetlands and settlements/infrastructure and agriculture | DIK·1&BP·1&VP·2&VEG·1&AI·2&JP·2 | 1 | 11.4 |
| 174 | IS-16 | Inland undulating plain below the forest line with wetlands and village/small town | DIK·1&BP·1&VP·2&VEG·1&AI·3&JP·1 | 1 | 16.0 |
| 175 | IS-17 | Inland undulating plain with boreal heath below the forest line with wetlands | DIK·1&BP·1&VP·2&VEG·2&AI·1&JP·1 | 148 | 1520.3 |

| MiT no. | MiT Code | Minor type – descriptive name | CLG segment combination | n | Total area km² |
|---|---|---|---|---|---|
| 176 | IS-18 | Inland undulating heath mountain plain with wetlands | DIK·1&BP·1&VP·2&VEG·3&AI·1&JP·1 | 81 | 673.6 |
| 177 | IS-19 | Inland undulating barren mountain plain with wetlands | DIK·1&BP·1&VP·2&VEG·4&AI·1&JP·1 | 13 | 91.2 |
| 178 | IS-20 | Coast-near undulating plain below the forest line | DIK·2&BP·1&VP·1&VEG·1&AI·1&JP·1 | 23 | 214.6 |
| 179 | IS-21 | Coast-near undulating plain below the forest line with settlements/infrastructure | DIK·2&BP·1&VP·1&VEG·1&AI·2&JP·1 | 16 | 139.3 |
| 180 | IS-22 | Coast-near undulating plain below the forest line with settlements/infrastructure and agriculture | DIK·2&BP·1&VP·1&VEG·1&AI·2&JP·2 | 74 | 559.1 |
| 181 | IS-23 | Coast-near undulating plain below the forest line with village/small town | DIK·2&BP·1&VP·1&VEG·1&AI·3&JP·1 | 13 | 93.2 |
| 182 | IS-24 | Coast-near undulating plain below the forest line with village/small town and agriculture | DIK·2&BP·1&VP·1&VEG·1&AI·3&JP·2 | 28 | 238.1 |
| 183 | IS-25 | Coast-near undulating plain below the forest line with city | DIK·2&BP·1&VP·1&VEG·1&AI·4&JP·1 | 2 | 17.5 |
| 184 | IS-26 | Coast-near undulating plain with boreal heath below the forest line | DIK·2&BP·1&VP·1&VEG·2&AI·1&JP·1 | 14 | 143.4 |
| 185 | IS-27 | Coast-near undulating heath mountain plain | DIK·2&BP·1&VP·1&VEG·3&AI·1&JP·1 | 13 | 103.2 |
| 186 | IS-28 | Coast-near undulating barren mountain plain | DIK·2&BP·1&VP·1&VEG·4&AI·1&JP·1 | 23 | 176.5 |
| 187 | IS-29 | Coast-near undulating barren mountain plain with glacier | DIK·2&BP·2&VP·1&VEG·4&AI·1&JP·1 | 1 | 8.4 |
| 188 | IS-30 | Coast-near undulating plain below the forest line with wetlands | DIK·2&BP·1&VP·2&VEG·1&AI·1&JP·1 | 41 | 343.7 |
| 189 | IS-31 | Coast-near undulating plain below the forest line with wetlands and agriculture | DIK·2&BP·1&VP·2&VEG·1&AI·1&JP·2 | 1 | 6.7 |
| 190 | IS-32 | Coast-near undulating plain below the forest line with wetlands and settlements/infrastructure | DIK·2&BP·1&VP·2&VEG·1&AI·2&JP·1 | 6 | 42.2 |
| 191 | IS-33 | Coast-near undulating plain below the forest line with wetlands and settlements/infrastructure and agriculture | DIK·2&BP·1&VP·2&VEG·1&AI·2&JP·2 | 2 | 18.4 |
| 192 | IS-34 | Coast-near undulating plain with boreal heath below the forest line with wetlands | DIK·2&BP·1&VP·2&VEG·2&AI·1&JP·1 | 5 | 51.3 |
| 193 | IS-35 | Coast-near undulating heath mountain plain with wetlands | DIK·2&BP·1&VP·2&VEG·3&AI·1&JP·1 | 12 | 116.0 |
| 194 | IS-36 | Coast-near undulating barren mountain plain with wetlands | DIK·2&BP·1&VP·2&VEG·4&AI·1&JP·1 | 2 | 14.2 |
| | | **Coastal fjords** | | | |
| 195 | KF-1 | Open fjord | RED·1&VP·1&AI·1&JP·1 | 175 | 1648.6 |
| 196 | KF-2 | Open fjord with settlements/infrastructure | RED·1&VP·1&AI·2&JP·1 | 219 | 2065.0 |
| 197 | KF-3 | Open fjord with settlements/infrastructure and agriculture | RED·1&VP·1&AI·2&JP·2 | 5 | 48.6 |
| 198 | KF-4 | Open fjord with village/small town | RED·1&VP·1&AI·3&JP·1 | 48 | 453.2 |
| 199 | KF-5 | Open fjord with city | RED·1&VP·1&AI·4&JP·1 | 1 | 4.4 |
| 200 | KF-6 | Open fjord with wetlands | RED·1&VP·2&AI·1&JP·1 | 1 | 10.4 |
| 201 | KF-7 | Open fjord with wetlands and settlements/infrastructure | RED·1&VP·2&AI·2&JP·1 | 4 | 26.1 |
| 202 | KF-8 | Relatively open fjord | RED·2&VP·1&AI·1&JP·1 | 1003 | 9133.6 |
| 203 | KF-9 | Relatively open fjord with settlements/infrastructure | RED·2&VP·1&AI·2&JP·1 | 1060 | 9660.5 |
| 204 | KF-10 | Relatively open fjord with settlements/infrastructure and agriculture | RED·2&VP·1&AI·2&JP·2 | 17 | 102.1 |
| 205 | KF-11 | Relatively open fjord with village/small town | RED·2&VP·1&AI·3&JP·1 | 117 | 1182.3 |
| 206 | KF-12 | Relatively open fjord with village/small town and agriculture | RED·2&VP·1&AI·3&JP·2 | 9 | 88.3 |
| 207 | KF-13 | Relatively open fjord with city | RED·2&VP·1&AI·4&JP·1 | 4 | 43.5 |
| 208 | KF-14 | Relatively open fjord with wetlands | RED·2&VP·2&AI·1&JP·1 | 12 | 133.8 |
| 209 | KF-15 | Relatively open fjord with wetlands and settlements/infrastructure | RED·2&VP·2&AI·2&JP·1 | 8 | 91.0 |
| 210 | KF-16 | Relatively open fjord with wetlands and village/small town | RED·2&VP·2&AI·3&JP·1 | 1 | 12.7 |
| 211 | KF-17 | Narrow fjord | RED·3&VP·1&AI·1&JP·1 | 587 | 4934.8 |

| MiT no. | MiT Code | Minor type – descriptive name | CLG segment combination | n | Total area km² |
|---|---|---|---|---|---|
| 212 | KF-18 | Narrow fjord with settlements/infrastructure | RED·3&VP·1&AI·2&JP·1 | 267 | 2355.0 |
| 213 | KF-19 | Narrow fjord with settlements/infrastructure and agriculture | RED·3&VP·1&AI·2&JP·2 | 2 | 16.2 |
| 214 | KF-20 | Narrow fjord with village/small town | RED·3&VP·1&AI·3&JP·1 | 27 | 301.4 |
| 215 | KF-21 | Narrow fjord with village/small town and agriculture | RED·3&VP·1&AI·3&JP·2 | 1 | 8.4 |
| 216 | KF-22 | Narrow fjord with wetlands | RED·3&VP·2&AI·1&JP·1 | 2 | 11.4 |
| 217 | KF-23 | Narrow fjord with wetlands and settlements/infrastructure | RED·3&VP·2&AI·2&JP·1 | 1 | 12.6 |
| 218 | KF-24 | Deeply cut fjord | RED·4&VP·1&AI·1&JP·1 | 127 | 988.3 |
| 219 | KF-25 | Deeply cut fjord with settlements/infrastructure | RED·4&VP·1&AI·2&JP·1 | 16 | 143.1 |
| 220 | KF-26 | Deeply cut fjord with village/small town | RED·4&VP·1&AI·3&JP·1 | 1 | 5.5 |
| | | **Coastal plains** | | | |
| 221 | KS-1 | Sheltered inner coastal plain | IYK·1,2&RE-KS·1&VP·1&AI·1,2&JP·1 | 69 | 640.4 |
| 222 | KS-2 | Sheltered inner coastal plain with agriculture | IYK·1,2&RE-KS·1&VP·1&AI·1,2&JP·2 | 22 | 169.2 |
| 223 | KS-3 | Sheltered inner coastal plain with village/small town | IYK·1,2&RE-KS·1&VP·1&AI·3&JP·1 | 19 | 144.6 |
| 224 | KS-4 | Sheltered inner coastal plain with village/small town and agriculture | IYK·1,2&RE-KS·1&VP·1&AI·3&JP·2 | 6 | 47.3 |
| 225 | KS-5 | Sheltered inner coastal plain with city | IYK·1,2&RE-KS·1&VP·1&AI·4&JP·1 | 2 | 9.1 |
| 226 | KS-6 | Sheltered inner coastal plain with wetlands | IYK·1,2&RE-KS·1&VP·2&AI·1,2&JP·1 | 61 | 545.0 |
| 227 | KS-7 | Sheltered inner coastal plain with wetlands and agriculture | IYK·1,2&RE-KS·1&VP·2&AI·1,2&JP·2 | 2 | 15.8 |
| 228 | KS-8 | Sheltered inner coastal plain with wetlands with village/small town | IYK·1,2&RE-KS·1&VP·2&AI·3&JP·1 | 1 | 4.2 |
| 229 | KS-9 | Sheltered inner flat coastal plain | IYK·1,2&RE-KS·2&VP·1&AI·1,2&JP·1 | 674 | 5140.7 |
| 230 | KS-10 | Sheltered inner undulating coastal plain with agriculture | IYK·1,2&RE-KS·2&VP·1&AI·1,2&JP·2 | 58 | 449.8 |
| 231 | KS-11 | Sheltered inner undulating coastal plain with village/small town | IYK·1,2&RE-KS·2&VP·1&AI·3&JP·1 | 176 | 1418.6 |
| 232 | KS-12 | Sheltered inner undulating coastal plain with village/small town and agriculture | IYK·1,2&RE-KS·2&VP·1&AI·3&JP·2 | 21 | 164.4 |
| 233 | KS-13 | Sheltered inner undulating coastal plain with city | IYK·1,2&RE-KS·2&VP·1&AI·4&JP·1 | 10 | 96.8 |
| 234 | KS-14 | Sheltered inner undulating coastal plain with city and agriculture | IYK·1,2&RE-KS·2&VP·1&AI·4&JP·2 | 1 | 6.6 |
| 235 | KS-15 | Sheltered inner undulating coastal plain with wetlands | IYK·1,2&RE-KS·2&VP·2&AI·1,2&JP·1 | 58 | 444.0 |
| 236 | KS-16 | Sheltered inner undulating coastal plain with wetlands and agriculture | IYK·1,2&RE-KS·2&VP·2&AI·1,2&JP·2 | 1 | 8.9 |
| 237 | KS-17 | Sheltered inner rugged coastal plain | IYK·1,2&RE-KS·3&VP·1&AI·1,2&JP·1 | 48 | 443.4 |
| 238 | KS-18 | Sheltered inner rugged coastal plain with agriculture | IYK·1,2&RE-KS·3&VP·1&AI·1,2&JP·2 | 1 | 4.4 |
| 239 | KS-19 | Sheltered inner rugged coastal plain with village/small town | IYK·1,2&RE-KS·3&VP·1&AI·3&JP·1 | 3 | 32.9 |
| 240 | KS-20 | Sheltered inner rugged coastal plain with wetlands | IYK·1,2&RE-KS·3&VP·2&AI·1,2&JP·1 | 2 | 25.2 |
| 241 | KS-21 | Moderately wave-exposed flat coastal plain | IYK·3&RE-KS·1&VP·1&AI·1,2&JP·1 | 124 | 1064.8 |
| 242 | KS-22 | Moderately wave-exposed flat coastal plain with agriculture | IYK·3&RE-KS·1&VP·1&AI·1,2&JP·2 | 16 | 142.5 |
| 243 | KS-23 | Moderately wave-exposed flat coastal plain with village/small town | IYK·3&RE-KS·1&VP·1&AI·3&JP·1 | 15 | 135.9 |
| 244 | KS-24 | Moderately wave-exposed flat coastal plain with village/small town and agriculture | IYK·3&RE-KS·1&VP·1&AI·3&JP·2 | 8 | 71.4 |
| 245 | KS-25 | Moderately wave-exposed flat coastal plain with city | IYK·3&RE-KS·1&VP·1&AI·4&JP·1 | 1 | 8.4 |
| 246 | KS-26 | Moderately wave-exposed flat coastal plain with wetlands | IYK·3&RE-KS·1&VP·2&AI·1,2&JP·1 | 44 | 452.4 |
| 247 | KS-27 | Moderately wave-exposed undulating coastal plain | IYK·3&RE-KS·2&VP·1&AI·1,2&JP·1 | 634 | 4869.9 |
| 248 | KS-28 | Moderately wave-exposed undulating coastal plain with agriculture | IYK·3&RE-KS·2&VP·1&AI·1,2&JP·2 | 19 | 127.6 |

| MiT no. | MiT Code | Minor type – descriptive name | CLG segment combination | n | Total area km² |
|---|---|---|---|---|---|
| 249 | KS-29 | Moderately wave-exposed undulating coastal plain with village/small town | IYK·3&RE-KS·2&VP·1&AI·3&JP·1 | 76 | 596.1 |
| 250 | KS-30 | Moderately wave-exposed undulating coastal plain with village/small town and agriculture | IYK·3&RE-KS·2&VP·1&AI·3&JP·2 | 3 | 25.2 |
| 251 | KS-31 | Moderately wave-exposed undulating coastal plain with city | IYK·3&RE-KS·2&VP·1&AI·4&JP·1 | 7 | 61.5 |
| 252 | KS-32 | Moderately wave-exposed undulating coastal plain with wetlands | IYK·3&RE-KS·2&VP·2&AI·1,2&JP·1 | 21 | 172.4 |
| 253 | KS-33 | Moderately wave-exposed undulating coastal plain with wetlands with village/small town | IYK·3&RE-KS·2&VP·2&AI·3&JP·1 | 1 | 10.0 |
| 254 | KS-34 | Moderately wave-exposed rugged coastal plain | IYK·3&RE-KS·3&VP·1&AI·1,2&JP·1 | 52 | 466.5 |
| 255 | KS-35 | Moderately wave-exposed rugged coastal plain with agriculture | IYK·3&RE-KS·3&VP·1&AI·1,2&JP·2 | 2 | 27.8 |
| 256 | KS-36 | Moderately wave-exposed rugged coastal plain with village/small town | IYK·3&RE-KS·3&VP·1&AI·3&JP·1 | 4 | 43.0 |
| 257 | KS-37 | Moderately wave-exposed rugged coastal plain with wetlands and agriculture | IYK·3&RE-KS·3&VP·2&AI·1,2&JP·2 | 1 | 6.4 |
| 258 | KS-38 | Very wave-exposed outer flat coastal plain | IYK·4&RE-KS·1&VP·1&AI·1,2&JP·1 | 218 | 1961.7 |
| 259 | KS-39 | Very wave-exposed outer flat coastal plain with agriculture | IYK·4&RE-KS·1&VP·1&AI·1,2&JP·2 | 23 | 187.8 |
| 260 | KS-40 | Very wave-exposed outer flat coastal plain with village/small town | IYK·4&RE-KS·1&VP·1&AI·3&JP·1 | 6 | 48.5 |
| 261 | KS-41 | Very wave-exposed outer flat coastal plain with village/small town and agriculture | IYK·4&RE-KS·1&VP·1&AI·3&JP·2 | 3 | 31.2 |
| 262 | KS-42 | Very wave-exposed outer flat coastal plain with wetlands | IYK·4&RE-KS·1&VP·2&AI·1,2&JP·1 | 22 | 212.4 |
| 263 | KS-43 | Very wave-exposed outer flat coastal plain with wetlands with village/small town | IYK·4&RE-KS·1&VP·2&AI·3&JP·1 | 1 | 22.1 |
| 264 | KS-44 | Very wave-exposed outer undulating coastal plain | IYK·4&RE-KS·2&VP·1&AI·1,2&JP·1 | 844 | 6488.2 |
| 265 | KS-45 | Very wave-exposed outer undulating coastal plain with agriculture | IYK·4&RE-KS·2&VP·1&AI·1,2&JP·2 | 6 | 50.0 |
| 266 | KS-46 | Very wave-exposed outer undulating coastal plain with village/small town | IYK·4&RE-KS·2&VP·1&AI·3&JP·1 | 44 | 394.6 |
| 267 | KS-47 | Very wave-exposed outer undulating coastal plain with village/small town and agriculture | IYK·4&RE-KS·2&VP·1&AI·3&JP·2 | 2 | 13.8 |
| 268 | KS-48 | Very wave-exposed outer undulating coastal plain with city | IYK·4&RE-KS·2&VP·1&AI·4&JP·1 | 1 | 11.1 |
| 269 | KS-49 | Very wave-exposed outer undulating coastal plain with wetlands | IYK·4&RE-KS·2&VP·2&AI·1,2&JP·1 | 13 | 100.4 |
| 270 | KS-50 | Very wave-exposed outer rugged coastal plain | IYK·4&RE-KS·3&VP·1&AI·1,2&JP·1 | 58 | 540.9 |
| 271 | KS-51 | Very wave-exposed outer rugged coastal plain with village/small town | IYK·4&RE-KS·3&VP·1&AI·3&JP·1 | 6 | 68.6 |
| 272 | KS-52 | Extremely wave-exposed outer flat coastal plain | IYK·5&RE-KS·1&VP·1&AI·1,2&JP·1 | 198 | 1775.5 |
| 273 | KS-53 | Extremely wave-exposed outer flat coastal plain with agriculture | IYK·5&RE-KS·1&VP·1&AI·1,2&JP·2 | 4 | 33.6 |
| 274 | KS-54 | Extremely wave-exposed outer flat coastal plain with village/small town | IYK·5&RE-KS·1&VP·1&AI·3&JP·1 | 5 | 37.1 |
| 275 | KS-55 | Extremely wave-exposed outer flat coastal plain with wetlands | IYK·5&RE-KS·1&VP·2&AI·1,2&JP·1 | 5 | 36.2 |
| 276 | KS-56 | Extremely wave-exposed outer flat coastal plain with wetlands and village/small town | IYK·5&RE-KS·1&VP·2&AI·3&JP·1 | 1 | 5.6 |
| 277 | KS-57 | Extremely wave-exposed outer flat undulating coastal plain | IYK·5&RE-KS·2&VP·1&AI·1,2&JP·1 | 720 | 5809.2 |
| 278 | KS-58 | Extremely wave-exposed outer undulating coastal plain with village/small town | IYK·5&RE-KS·2&VP·1&AI·3&JP·1 | 1 | 10.1 |
| 279 | KS-59 | Extremely wave-exposed outer undulating coastal plain with village/small town and agriculture | IYK·5&RE-KS·2&VP·1&AI·3&JP·2 | 1 | 5.5 |
| 280 | KS-60 | Extremely wave-exposed outer undulating coastal plain with city | IYK·5&RE-KS·2&VP·1&AI·4&JP·1 | 1 | 9.8 |
| 281 | KS-61 | Extremely wave-exposed outer undulating coastal plain with wetlands | IYK·5&RE-KS·2&VP·2&AI·1,2&JP·1 | 2 | 11.5 |
| 282 | KS-62 | Extremely wave-exposed outer rugged coastal plain | IYK·5&RE-KS·3&VP·1&AI·1,2&JP·1 | 27 | 282.2 |
| 283 | KS-63 | Extremely wave-exposed outer rugged coastal plain with wetlands | IYK·5&RE-KS·3&VP·2&AI·1,2&JP·1 | 1 | 5.4 |
| | | **Coastal hills and mountains** | | | |
| 284 | KA-1 | Coastal hills- and mountains | Not appliccable | 77 | 273.6 |

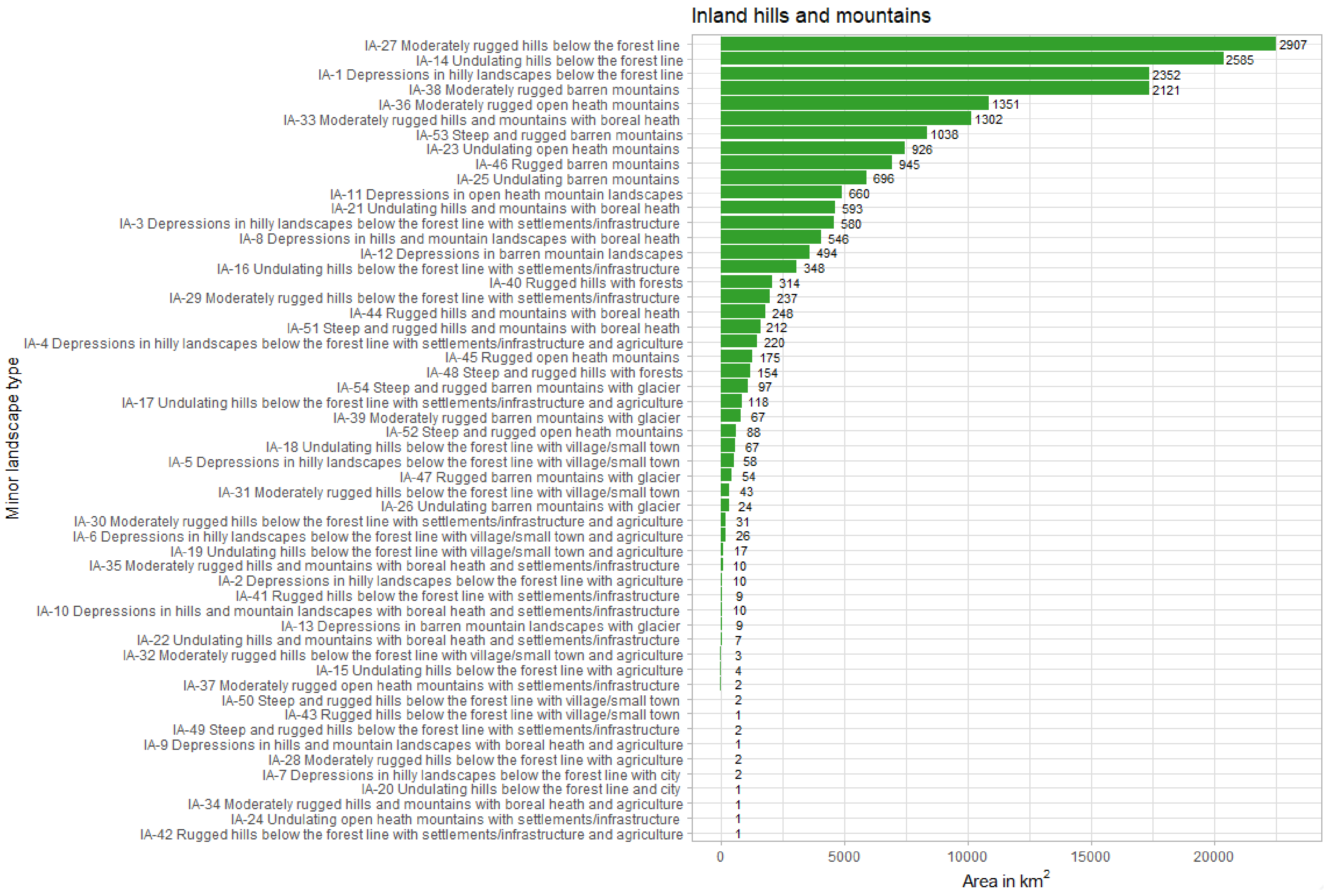


**Fig. S9.1.** Bar plot showing total area for each minor landscape type within inland hills and mountains. The number outside the bar refers to number of spatial landscape units for each minor landscape type.

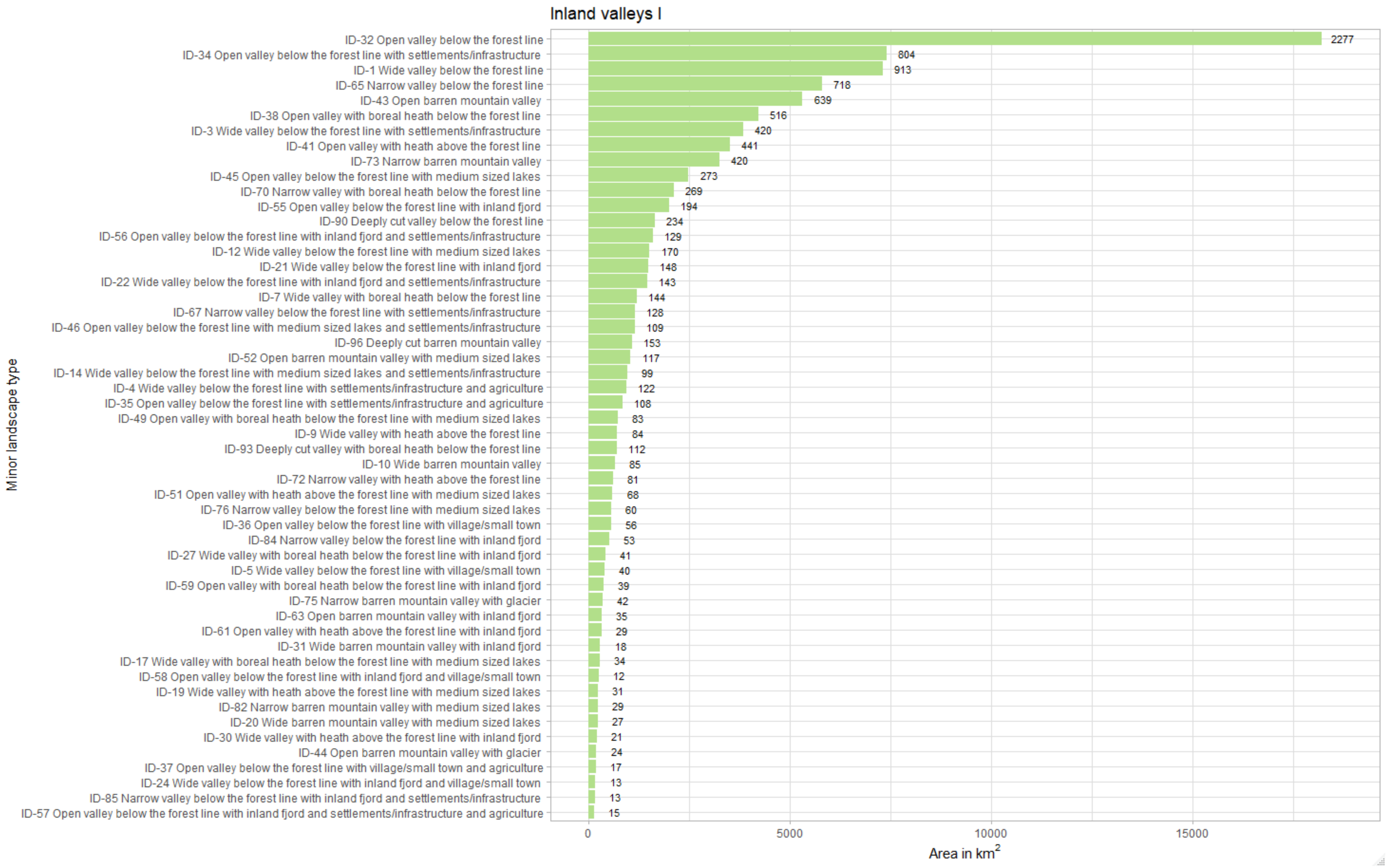


**Fig. S9.2.** Bar plot showing total area for each minor landscape type within inland valleys (1 of 2, see also next page). The number outside the bar refers to number of spatial landscape units for each minor landscape type.

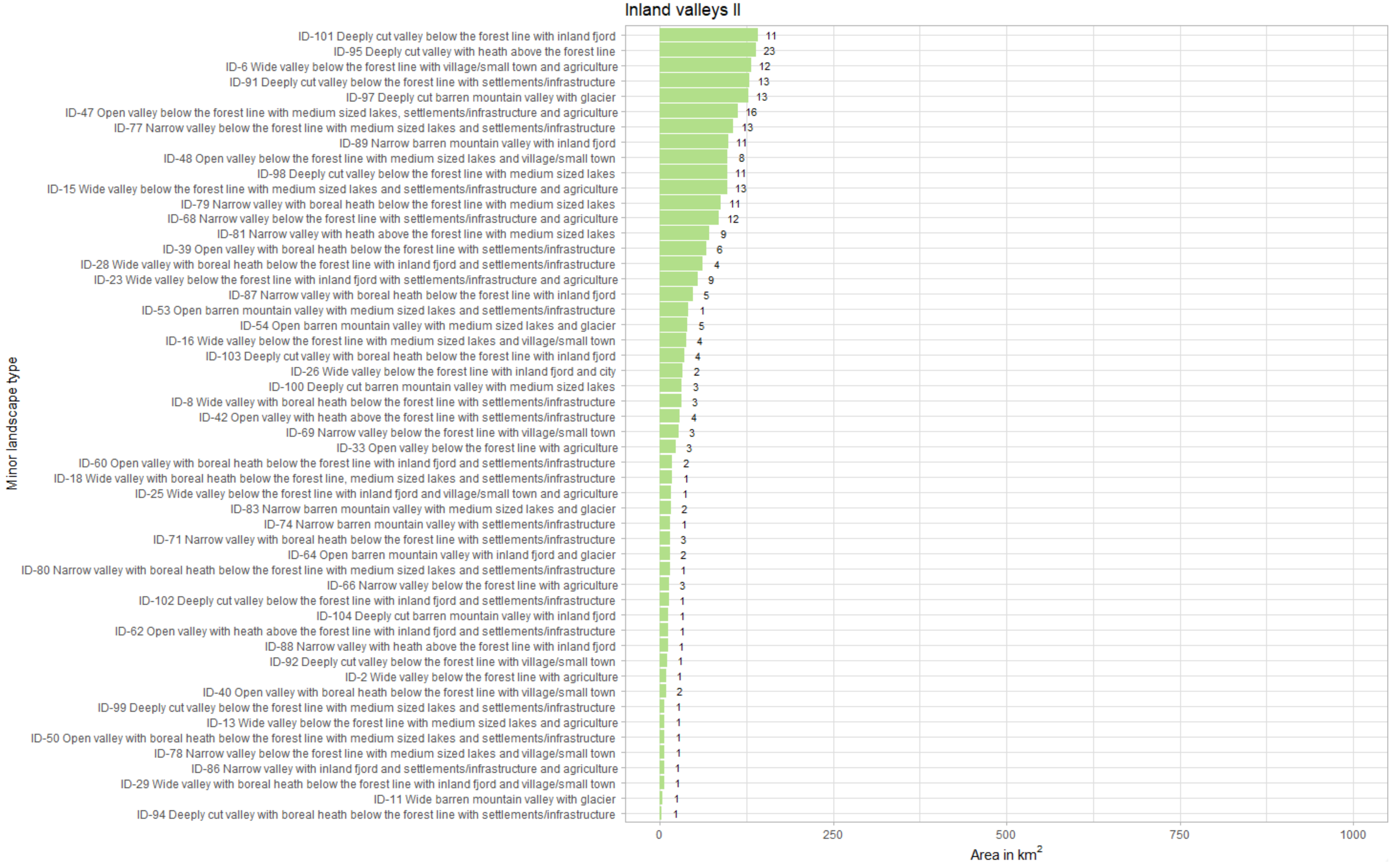


**Fig. S9.3.** Bar plot showing total area for each minor landscape type within inland valleys (2 of 2, see also previous page). The number outside the bar refers to number of spatial landscape units for each minor landscape type.

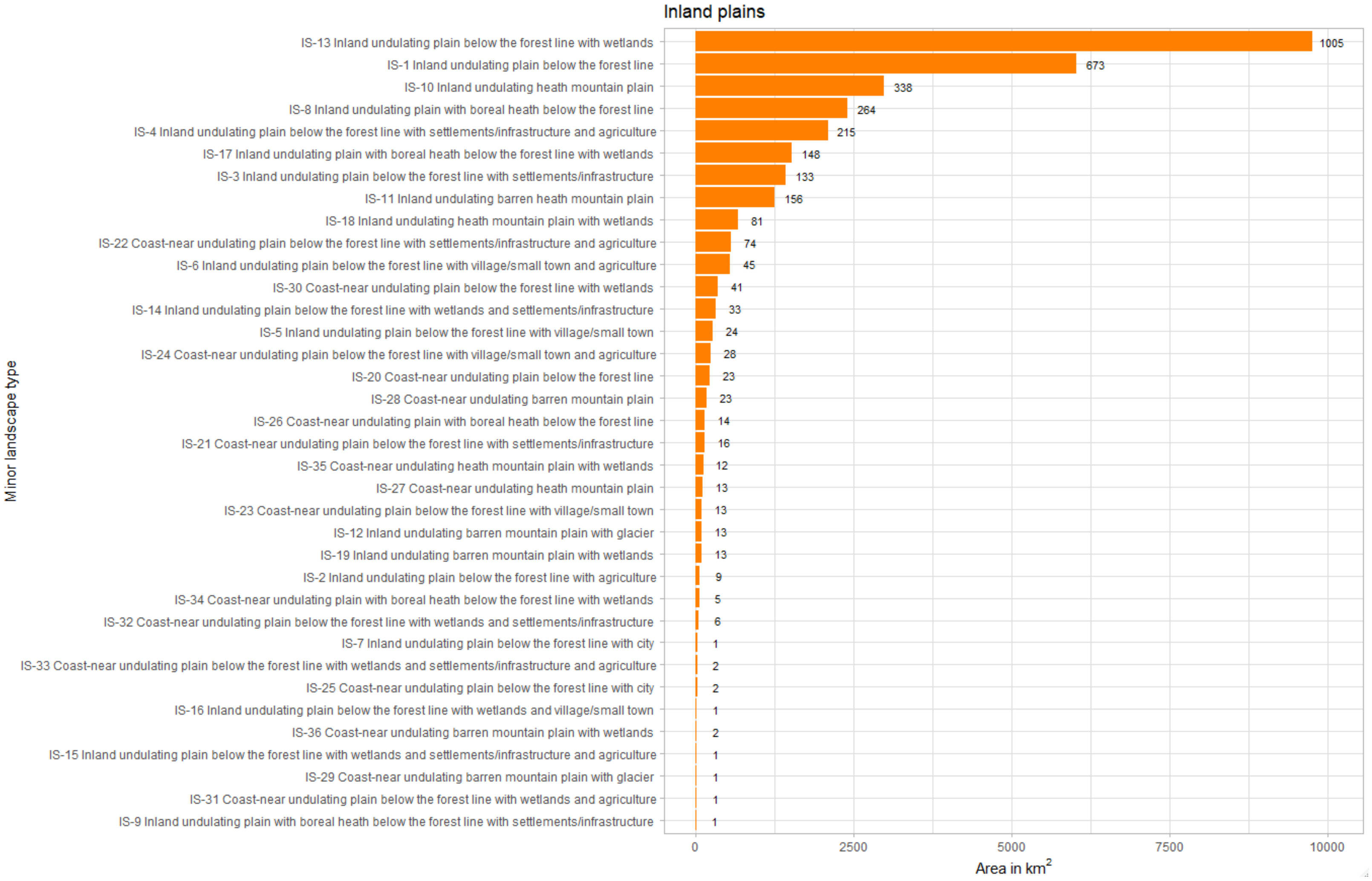


**Fig. S9.4.** Bar plot showing total area for each minor landscape type within inland plains. The number outside the bar refers to number of spatial landscape units for each minor landscape type.

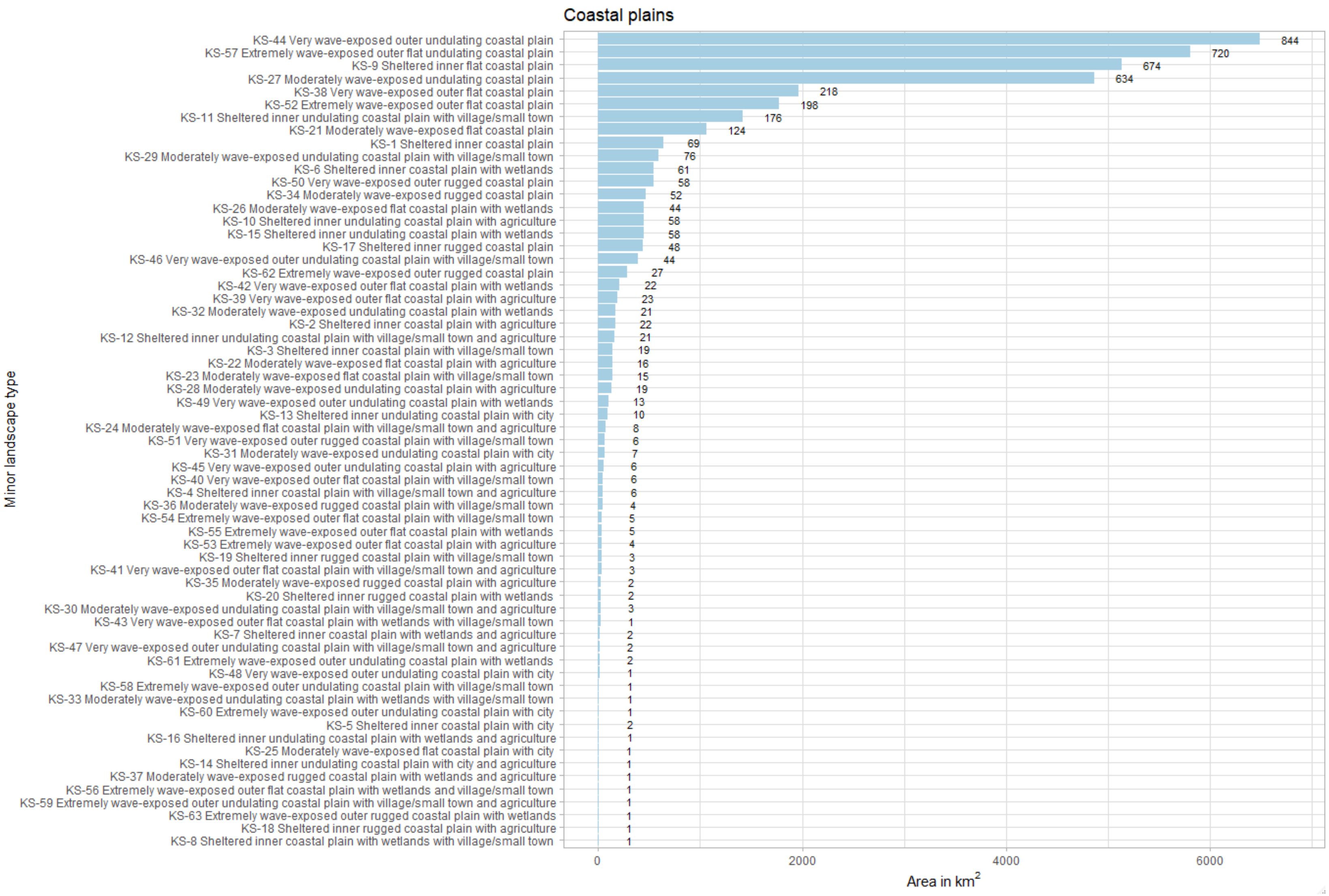


**Fig. S9.5.** Bar plot showing total area for each minor landscape type within coastal plains. The number outside the bar refers to number of spatial landscape units for each minor landscape type.

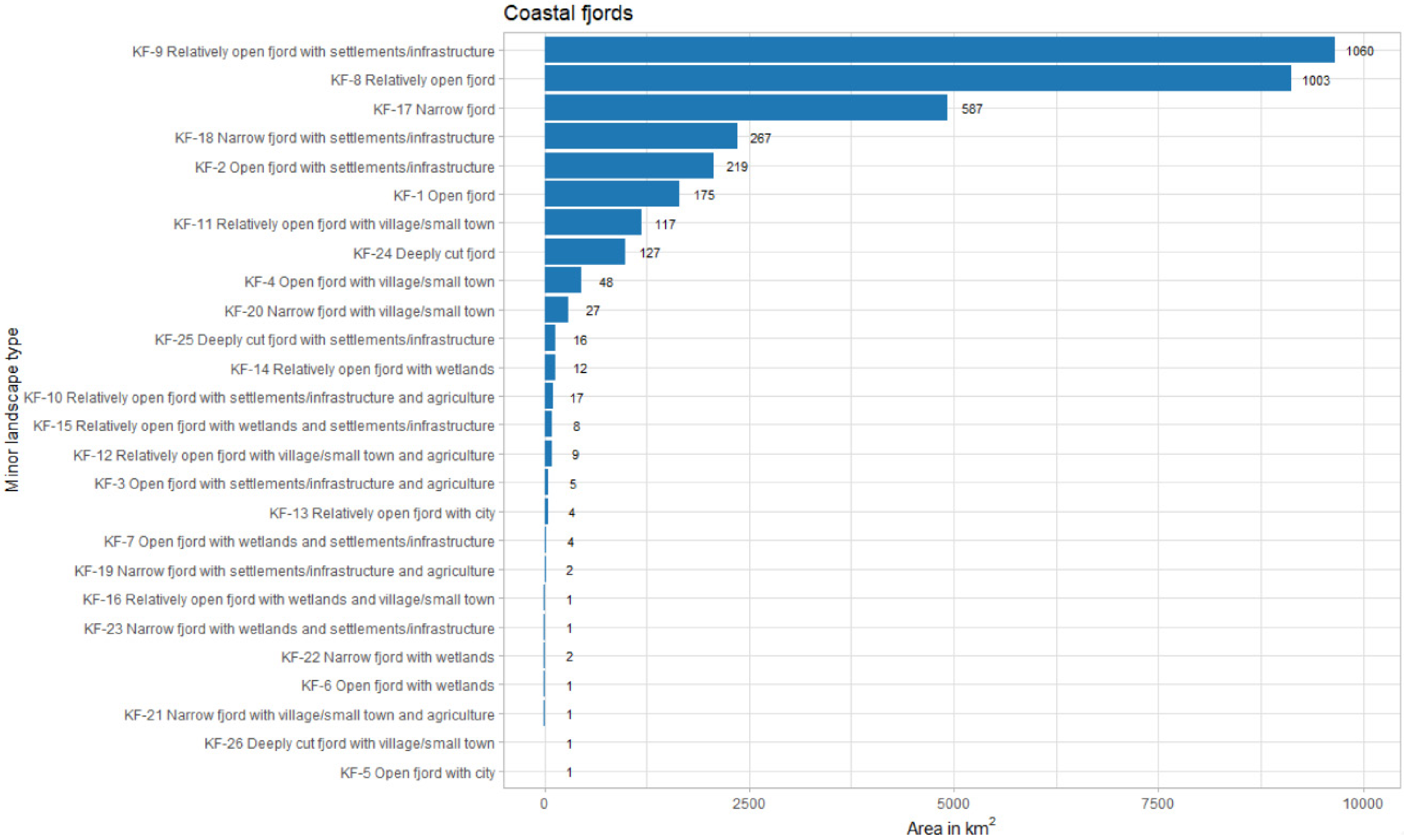


**Fig. S9.6.** Bar plot showing total area for each minor landscape type within coastal fjords. The number outside the bar refers to number of spatial landscape units for each minor landscape type.

### S10 Validation

Table. S9. Comparison of number of minor landscape types in the tentative type system derived from statistical analyses from a sample of observation units throughout Norway (‘analysis’) and realised minor types in the applied landscape type mapping (‘mapping’).

| **Major type** | **Minor types** | |
|---|---|---|
| | Analysis | Mapping |
| **Coastal plains** | 51 | 63 |
| **Coastal fjords** | 31 | 26 |
| **Coastal hills and mountains** | NA | 1 |
| **Inland valleys** | 98 | 104 |
| **Inland hills and mountains** | 41 | 54 |
| **Inland plains** | 81 | 36 |
| Total | **302** | **284** |
